# Trait-based paleontological niche prediction demonstrates deep time parallel ecological occupation in specialized ant predators

**DOI:** 10.1101/2022.06.09.495514

**Authors:** Christine E. Sosiak, Tyler Janovitz, Vincent Perrichot, John Paul Timonera, Phillip Barden

## Abstract

Understanding the paleoecology of species is fundamental to reconstructions of paleoecological communities, analyses of changing paleoenvironments, and the evolutionary history of many lineages. One method of establishing paleoecology is through comparing the morphology of extant analogs to extinct species; this method has been applied to many vertebrate groups using predictive linear models, but has been rarely applied using invertebrate taxa or non-linear frameworks. The ant fossil record, which frequently preserves specimens in three-dimensional fidelity, chronicles putative faunal turnover during the Late Cretaceous and into Cenozoic. The earliest fossil species comprise enigmatic stem-groups that underwent extinction concomitant with crown ant diversification. Here, we apply a wide-ranging extant ecomorphological dataset to demonstrate the utility of Random Forest machine learning classification in predicting the ecology of stem-group “hell ants”. We reconstruct a predicted ecomorphological assemblage of this phenotypically aberrant group of extinct ants, and compare predicted hell ant ecologies to the ecological occupations of their closest living analogs, lineages of solitary predators with highly specialized mandibular morphology. In contrast to previous hypotheses, we find that hell ants were primarily leaf litter or ground-nesting and foraging taxa, and that the ecological breadth of this unusual lineage mirrored that of living groups. Results suggest ecological coherence between the Mesozoic and modern communities, even as the earliest occupants of predatory niches were phylogenetically and morphologically distinct.

## Introduction

Estimating the ecological niche occupation of extinct taxa is a central component of paleontology. The putative ecologies of extinct organisms are routinely incorporated into analyses of extinction risk, paleoenvironmental reconstruction, and lineage evolutionary history (Palmqvist et al. 2003; Benson et al. 2014; Frederickson et al. 2018). Even as aspects of extinct niche occupation may be reliably inferred by the preservation of individual traits in fossil specimens, organismal ecology remains multifaceted. Morphology frequently reflects ecology across entire phenotypes (Williams 1972; Losos 1992) and phenotypic syndromes may be linked to multiple aspects of an organism’s ecological niche spanning habitat, diet, and interactions. These ecologically-linked body plans - ecomorphs - are found in such disparate taxa as fish, reptiles, arthropods, and mammals (Barton et al. 2011; Gerry et al. 2011; Sanders et al. 2013; Figueirido et al. 2019). The relationship between ecological niche and multi-trait morphology can also be leveraged to estimate paleoecologies. In lineages with surviving relatives, extant taxa may serve as analogs for ecological niche estimation, however a partial fossil record often reduces the utility of extant-to-extinct comparisons.

Across vertebrate species, limb anatomy is a predictor of locomotion, prey items, and substrate behaviour. In particular, the forelimb anatomy of carnivores has been used to predict the likely predatory habits and prey size of extinct carnivorous mammals and mammaliaforms (Dunn et al. 2019; Ercoli et al. 2012; Figueirido et al. 2016; Jenkins et al. 2020; Lungmus and Angielczyk 2021; Meloro and Louys 2014). Other examples of extant morphology used in the prediction of extinct ecology include estimating prey type in extinct raptor species from present-day birds of prey (Hertel 1995); predicting prey items and modes of scavenging in extinct crocodyliforms from extant crocodyliform snout morphology (Drumheller and Wilberg 2019); approximating arboreal behaviour in extinct primates (Rector and Vergamini 2018); and predicting habitat preferences in fossil Anolis lizards using ear canal shape (Dickson et al. 2017). In most cases, potentially because of the difficulty of obtaining many complete skeletons, these methods incorporate isolated body parts rather than full-body morphology; however, morphometric analyses of mammalian skeletons have been used to predict the locomotion mode of various Mesozoic mammaliaforms (Chen and Wilson 2015; Meng et al. 2017).

Many attempts to predict paleoecology from extant morphology use techniques such as canonical correlations analysis or canonical variate analysis, and in particular linear discriminant analysis (Dickson et al. 2017; Dunn et al. 2019; Hertel 1995; Janis and Figueirido 2014; Meloro and Louys 2014; Rector and Vergamini 2018). These methods maximize variation in the measured traits between predetermined classes in morphospace. The fossil specimen’s most likely ecology is then determined by proximity to each class mean in this constructed morphospace (Strauss 2010). These approaches are powerful tools for establishing sets of traits most strongly associated with predetermined classes, but are limited to linear relationships only among measured traits, which may restrict accuracy by failing to incorporate non-linear predictive relationships between traits.

While most predictive paleoecology studies have focused on vertebrate paleoecology, minimal attention has been paid to these approaches in invertebrates, particularly insects. Many extant insect lineages reach back to the Mesozoic or Paleozoic, and are highly ecologically diverse. One example are the ants, which arose between ~150 and 100 Ma (Mega-annum) (Moreau et al. 2006; Brady et al. 2006; Borowiec et al. 2019). With over 13,000 species comprising a significant component of terrestrial biomass, ants are globally ubiquitous, speciose, and are present in most post-producer ecological niches (Hölldobler and Wilson 1990). Importantly, despite this ecological diversity, ants are also morphologically conserved with respect to broad bauplan functionality and possess a rich fossil record extending back 100 Ma to the mid-Cretaceous. A majority of ant fossils are known from fossil amber, which often preserves entire specimens with high fidelity. Because of their well-defined homology and uniquely preserved fossil history, extinct ants are strongly suited for testing paleoecological niche prediction methods.

The earliest known ant fossils date to the Early–Late Cretaceous transition and evidence extinct stem-lineages that began to diversify prior to the most recent common ancestor of all living ants. Cretaceous fossils have fueled speculation on the ecological occupation of the earliest ants because these taxa have bearing on the evolution of eusociality more broadly. It was posited that these early species were unlikely to construct nests, but instead used already-present cavities in soil and wood (Dlussky 1996; Wilson 1967; Wilson 1987a; Wilson 1987b; Grimaldi and Agosti 2000; Engel and Grimaldi 2005). Based on their presumed wasp ancestors, they were additionally argued to be predators (Dlussky 1996; Wilson 1987a; Wilson 1987b). As the taxonomic diversity of extinct ant species increased, concordant with the discovery of new fossil occurrences, paleoclimate reconstructions and contextualization with other fossils and phylogenetic relationships suggested soil and leaf litter microhabitats in newly emerging angiosperm forests (Wilson and Hölldobler 2005, Perrichot et al. 2008, Moreau et al. 2006, Moreau and Bell 2013). Phylogenetic reconstructions of extant lineages have recovered ant ancestors as potentially hypogaeic soil-dwellers (Lucky et al. 2013).

Haidomyrmecines, or hell ants, are an enigmatic and highly specialized extinct subfamily of ants comprising 16 described species and 10 genera (Perrichot et al. 2020). Hell ants occupy a stem-group position relative to modern ants and are frequently recovered as sister to all other extinct and extant ants (Barden et al. 2016; 2020). They persisted throughout the mid-to-late Cretaceous as evidenced by amber fossils ranging from 100 to 78 Ma on three different continents in Canada, Myanmar, and France (Dlussky 1996; Perrichot et al. 2008; McKellar et al. 2013) and are hypothesized to have undergone extinction concomitant with the diversification of extant lineages. These ants are morphologically unique among the Formicidae in having vertically-articulating mandibles, unlike the horizontal alignment of modern ants. Remarkably, hell ants possess an array of horn-like cranial appendages that have been directly observed to facilitate solitary predation through fossil remains (Barden et al. 2020): haidomyrmecines captured prey individually by articulating their mandibles against their horns. Hell ants have been hypothesized as arboreal predators, considering the potential difficulty of substrate manipulation with their vertically-aligned mandibles, and frequent preservation in amber, potentially indicating close proximity to tree resin (Dlussky 1996; Barden and Grimaldi 2012; Lattke and Melo 2020).

What functional niches did the earliest ant predators occupy and how does this occupation compare with extant lineages? Here, we use a wide-ranging extant morphometric dataset spanning over 160 species that has previously demonstrated a quantitative link between morphology and ecology to predict the paleoecology of several hell ant species. Using a supervised machine learning classification algorithm, Random Forest, we predict foraging niche, nesting niche, and functional role for hell ants. With these predictions, we reconstruct known ecomorphological space for haidomyrmecines and compare these ecological occupations to those of extant lineages of specialized solitary predators. Our results demonstrate repeated filling of functional niche space across phylogenetically and morphologically distant lineages.

## Methods

We applied a supervised machine learning approach by training multiple random forest models on a comprehensive ecomorphological dataset of extant ant taxa. Briefly, for each ecological niche aspect, we constructed a total of four overarching models trained on four subsets of extant data to evaluate alternative predictive schemes based on body size and dataset completeness in anticipation of partial fossil specimens: i) a complete morphometric dataset comprising raw measurements; ii) a complete morphometric dataset comprising size-corrected ratio measurements; iii) a subset of the complete dataset comprising raw measurements; iv) a subset of the complete dataset comprising size-corrected ratio measurements. We then used each model to predict ecological niche aspects based on two alternate versions of input fossil data: one including measurements based on assumed homology of haidomyrmine ants and another assuming haidomyrmecine measurements corresponding with functional analogs of extant ants. Niche aspects of each specimen were then predicted using the appropriate model; the complete-dataset trained model for complete specimens, and subset-trained model for incomplete specimens. All analyses were performed in R v.4.0.3 (R Core Team, 2021).

### Development of the predictive model

#### Training dataset

We developed a predictive machine learning model using a training dataset of extant ant morphometrics with sampling following Sosiak & Barden (2021). Our extant dataset spans 15 subfamilies, 113 genera, and 167 species, including three specimens per species where possible, and measuring as many conspecifics as were present in museum collections otherwise. Polymorphic species are represented by the media caste, and species with specialized castes are represented by non-specialized “minor” workers. There is currently no evidence for specialized hell ant worker castes and specialized worker castes are not a synapomorphy of crown ants.

Our morphometric sampling comprised linear measurements of 12 cephalic traits and five post-cephalic traits. Most sampled traits have been previously linked to ecology (Weiser and Kaspari 2006; Yates and Andrew 2011; Gibb et al. 2015; Yates et al. 2014). All measurements were conducted on point-mounted specimens under stereo microscopy. Because body size in diverse species can drive the overwhelming majority of variation in a dataset and potentially mask other important contributors, we created two datasets: one comprising raw measurements and one comprising only size-corrected ratios. A list of all traits measured with associated definitions may be found in Supplementary Tables S1 and S2.

All specimens were assigned a binning from each of three ecological niche aspect categories: functional role (six binnings), nesting niche (five binnings), and foraging niche (four binnings). Specimen binnings were assigned based on literature surveys. When any particular aspect of a species’ ecological niche was uncertain, the species was assigned an “unknown” binning and excluded from further model development. We collapsed the 35 observed combinations of niche binnings across all niche aspects into 10 ecomorph syndromes, defined after preliminary analysis indicated that some niche combinations showed extensive morphological overlap. A list of all ecological niche aspect binnings and syndromes with associated definitions may be found in Supplementary Tables S3 and S4.

#### Model development

We implemented Random Forest analysis, a supervised machine learning algorithm, to delimit species into ecological niches by morphology using our extant ant morphometric dataset. Random Forest models are classification algorithms that partition morphospace according to predefined ecological niche binnings through an assemblage of decision trees (Breiman 2001). The algorithm builds a series of decision trees which each provide a “vote” on a given specimen’s predicted category; the vote consensus from the decision trees is then used as the final predicted category for the specimen. At each split in each decision tree, a predetermined number of variables are randomly selected and tested to evaluate their contribution to the accuracy of the model. During each iteration of the model, one-third of the data is randomly removed and used to estimate the error rate of that particular iteration; the converged test error rate across the iterations is considered the out-of-bag error rate for the model. This removal of testing data during bootstrapping eliminates the need for a priori separation of a testing and training dataset, incorporating all collected data.

While other supervised machine learning or dimension reduction techniques have been used in morphology-based paleoecological prediction, Random Forest has recently been shown to outperform linear predictive approaches with respect to morphology (Pigot et al. 2020; Sosiak & Barden 2021). This is the first application of Random Forest in predicting extinct ecology to our knowledge. Random Forest models were trained on our morphometric data, using both raw measurement datasets and size-corrected ratio measurement datasets separately for each ecological niche aspect. We constructed Random Forest models for functional role, nesting niche, foraging niche, and ecomorph syndrome separately, allowing for both granular classification of ecological niche and more synthesized classification. Model parameters were selected based on initial sensitivity tests: mtry=4 (number of variables tested at each split) and ntree=5000 (number of trees iterated). Out-of-bag error rate estimates ranged from 15-22%, reflecting accuracies of 78-85%; OOB error rates for each model are shown in Table S5. All Random Forest analyses were implemented in the R package “randomForest” (Liaw and Weiner 2018).

#### Fossil data

We measured fossil hell ant specimens using a combination of stereo microscopy and reconstructions of X-ray micro-computed tomography (micro-CT) scans. Twenty specimens from 16 species and morphospecies were measured under stereo microscopy. We submerged the amber specimens in water to reduce light distortion; some measurements were not possible due to specimen positioning. Three specimens were micro-CT scanned and reconstructed for subsequent measurements; two species of hell ant and one species of *Pseudomyrmex macrops* from Dominican amber to assess the reliability of CT-scan based data. Congeners of the Dominican *Pseudomyrmex* fossil are extant today and their ecology is well consistent across the genus and well-characterized. Two specimens, *Haidomyrmex scimitarus* (specimen AMNH Bu-FB80) and *Linguamyrmex vladi* (specimen AMNH BuPH-1) were scanned at the American Museum of Natural History Microscopy and Imaging Facility using a GE phoenix v|tome|x s240 60kV CT-scanner. Specimen AMNH Bu-FB80 was imaged at 180 μA for 5 second exposures and a voxel size of ~8μm; specimen AMNH BuPH-1 at 250 μA for 1 second exposures and a voxel size of ~3μm. The *Pseudomyrmex macrops* specimen (AMNH DR-14-1021) was imaged at the New Jersey Institute of Technology York Center using a Bruker SkyScan 1275 at 60kV and 150μA for 1 second exposures with a subsequent voxel size of ~3.5μm. Volume reconstruction of the x-ray images was conducted in 3D Slicer v4.11 (Fedorov et al. 2012) using the Segmentation modules; still images of the reconstructed specimens were subsequently imported into ImageJ (Abràmoff et al. 2004) for linear measurements to retain consistency with measurements taken under stereo microscopy.

Most hell ant traits were distinguished as homologous to extant species; however, because haidomyrmecine cranial morphology is highly modified, eye position is difficult to assess in the context of modern ant variation. Extant ants (and most extinct ants, including the Dominican *Pseudomyrmex* fossil) have a prognathous head posture, in which the long axis of the head is held parallel to the ground with the oral opening pointing anteriad. In many other insect taxa, such as wasps and grasshoppers, the head is held in a hypognathous posture; the long axis of the head is held perpendicular to the ground, with the oral opening positioned ventrally. A hypognathous-like head posture is also present in hell ants, a configuration not seen in any extant ant. This difference in head posture means that, interpreted strictly on the basis of homology, the dorsoventral axis of the extant ant head is the anteroposterior axis of the hell ant head, and vice versa; that eye positioning along each axis is similarly swapped. When interpreted this way, the eye position of the hell ant is more similar to many *Camponotus* species: positioned extremely posteriorly and more dorsally.

Functionally, however, the hell ant would have been moving with its head held hypognathously, as evidenced by fossilized posture as well as preserved predation (Perrichot et al 2008; Barden & Grimaldi 2012; Perrichot et al. 2016; Barden et al. 2020) thus the functional anteroposterior axis of the hell ant head would be from its clypeal protrusion or horn to the occipital foramen, rather than the homologous anteroposterior axis of the head running perpendicular to the ground from vertex to oral opening. Considering that our question is a question of functional ecology - what is the ecological occupation of hell ants? - we measured traits related to eye position both functionally and homologously, to incorporate this possible variation.

Due to limitations measuring specimens directly from amber fossils, we produced two fossil morphometric datasets: one ‘incomplete dataset’ which excluded a subset of traits for all specimens; and one with all measurements included. The incomplete dataset lacked the frontal head length, head width, frontal mandible length, and pronotal width measurements. These measurements are often difficult to accurately capture because amber fossils are typically prepared to expose a clear lateral profile of any inclusions, leaving the dorsal and frontal margins of the amber curved and distorted. Twenty hell ant specimens were included in this incomplete dataset.

The complete dataset comprised the proof-of-concept fossil *Pseudomyrmex macrops* specimen and three hell ant specimens: *Dhagnathos autokrator, Haidomyrmex scimitarus,* and *Linguamyrmex vladi.* While the majority of specimens were workers, two of the specimens - the complete *Dhagnathos autokrator* and *Haidomyrmex scimitarus* were represented by alate and dealate queens, respectively. We include these here for two reasons: one, hell ant queens are hypothesized to have actively foraged and hunted in early colony foundation, and so likely occupied a similar ecological niche to the workers of the species; and two, worker specimens are limited in many hell ant species and entirely absent in the genus *Dhagnathos.* While we have no comparison to *Dhagnathos* workers, we include a *Haidomyrmex scimitarus* worker in the incomplete morphometric dataset, allowing us to compare the accuracy of the model in predicting alate and worker ecology. A full list of all specimens included with information pertaining to their sampling methods and castes can be found in Table S6; all morphometric data for fossil specimens may be found in Supplementary Data_Fossil Morphometrics. To ensure that morphological diversity for traits measured from fossil species are within the bounds of extant morphological diversity, we conducted principal component analysis (PCA) to compare morphospace occupation of hell ants relative to extant lineages. We conducted separate PCAs for both the raw measurements and size-corrected ratio measurements. All PCAs were implemented in R packages “corrplot” (Wei et al. 2017) and “FactoMineR” (Lê et al. 2008).

### Random Forest model implementation

We implemented different sets of Random Forest models, given the missing traits in our incomplete fossil dataset; once with the missing traits eliminated from the extant ant morphometric training dataset, and once including all traits for the complete fossil dataset. For each dataset, we implemented two models using the raw measurements and the size-corrected ratio measurements. Using these two models, we predicted ecological niche aspects of hell ants twice; once using functional morphology, and once using homologous morphology. Thus, each specimen’s ecological niche was predicted four times: using raw measurements with functional morphology, raw measurements with homologous morphology, size-corrected ratio measurements with functional morphology, and size-corrected ratio measurements with homologous morphology. We compiled all posterior probabilities of each prediction into a heatmap of model predictions for each specimen with the R package “ggplot2” (Wickham 2016) to better visualize the consensus among models for each predicted ecological niche aspect.

### Comparative ecomorphological niche occupation

To assess the specificity and breadth of niche occupations in haidomyrmecines, we generated three-dimensional ecomorphological matrices comprising living and extinct predatory taxa. Our taxonomic sampling included five lineages: hell ants as well as all four distantly related extant groups with trap-jaw like morphology and behavior, wherein workers act as solitary hunters that capture prey; many, but not all, do so through rapid closure of specialized mandibles (Gronenberg and Ehmer 1996, Hölldobler & Wilson 1990). While the speed of prey capture is not known in hell ants, haidomyrmecines are united with some extant trap jaw taxa by the presence of elongate setae (interpreted as trigger hairs) in the path of mandible closure and dramatic morphological adaptations related to predation (Dlussky 1996; Barden & Grimaldi 2012; Perrichot et al. 2016; Barden et al. 2017). Importantly, all species within our sampling are primarily solitary hunters (Larabee et al. 2014; Barden et al. 2020) in contrast to group raiding or collective prey capture that typify many other predatory ant lineages (Dornhaus & Powell 2010). Trap jaw mechanisms have evolved at least ten times in living ants (Larabee et al. 2014; Booher et al. 2021); these origins are distributed among four monophyletic lineages. Extant trap jaw predation has evolved once within each of the subfamilies Ponerinae and Formicinae, thus our sampling included species within relevant genera: *Anochetus* + *Odontomachus* and *Myrmoteras,* respectively. There are at least eight trap jaw origins within the subfamily Myrmicinae and seven of these have occurred within the genus *Strumigenys*. Our myrmicine sampling therefore included Strumigenys as well as the five “dacetine” trap-jaw genera: *Acanthognathus, Daceton, Epopostruma, Microdaceton,* and *Orectognathus.* Although it is not yet clear whether all dacetine trap jaws are the product of a single origin, we grouped these taxa in analyses as they are more closely related to each other than any are to *Strumigenys* (Ward et al. 2015).

To estimate the total number of unique ecomorphological combinations for each specialized predatory lineage, we gathered niche occupation and size data for a total of 982 species, including 15 hell ant species with ecologies estimated under random forest. Our extant sampling represents a minimum of 50% species sampling for each genus. Each species was assigned one of three foraging and three nesting niche aspects according to our modeling results for haidomyrmecines or published natural history observations for extant taxa (Supplementary Data_Trap Jaw Ecomorphospace). Because observational data do not exist for all species, we applied published generalizations in some cases (e.g. *Strumigenys* are noted as almost always leaf litter nesting and foraging, we assumed this occupation by default except when otherwise noted in the literature).

To estimate body size, we gathered minimum and maximum Weber’s length (a measurement of mesosoma length and traditional metric of ant size) measurements from taxonomic descriptions and revisions. To include taxa without published morphometric data, we collected Weber’s length measurements from publicly available images on AntWeb (AntWeb 2021) using ImageJ. We discretized species sizes by delimiting Weber’s length ranges for each species into at least one of twelve equal size binnings. Size binning ranges were defined as one half of the standard deviation of Weber’s length measurements across all species. In cases where a species Weber’s length range exceeded any one size binning, we assigned multiple size binnings for that species.

We generated three-dimensional ecological disparity values for each lineage following a modification of Chen et al. (2019). We assigned each ecological binning a numerical value from 1 to 3 based on inferred ecological proximity (nesting niche: leaf litter = 1, ground = 2, lignicolous = 3; foraging niche: leaf litter = 1, epigaeic = 2, arboreal = 3), while values for the third ecological dimension, body size, were continuous from 1 to 12. To reduce the impact of species sampling bias between fossil and extant lineages, we calculated disparity only among unique occupations, not between species. We created a matrix of unique niche occupations for each lineage and calculated intra-lineage disparity values by summing the distances between niche aspects for all pairwise combinations of unique occupations using the equation. For example, the ecological disparity between unique occupation 1 (uo1) and unique occupation 2 (uo2) would be: | Nesting Nicheu_o1_ - Nesting Nicheu_o2_ | + | Foraging Niche_uoi_ - Foraging Niche_uo2_ | + | Body Size_uoi_ - Body Size_uo2_ |. We summarized mean and standard deviations for each lineage using ggplot. Visual representations of lineage specific ecomorphological occupations (ecospaces) were generated using the R package “rgl” (Adler et al. 2021) and redrawn in Adobe Illustrator.

## Results

### Model performance and prediction sensitivity

Visualization of extant and extinct morphospace through principal component analysis illustrated that extinct morphospace fully overlaps with extant morphospace representing raw trait measurements, and mostly overlaps with extant morphospace representing size-corrected ratio measurements (Supplemental Figures S3, S4). Although hell ant morphology is distinct, the hell ant morphospace represented by measurements incorporated in our models is primarily within the bounds of extant diversity. Principal component 1 in the size-corrected ratio morphospace is primarily driven by mandible size relative to body size, which is greater in many hell ants compared to extant ants, likely resulting in the small portion of unique hell ant morphospace. Predictions from size-corrected ratio measurements are often not significantly different from predictions using raw measurements, however, and this along with a great degree of morphospace overlap suggests that extant morphology is an appropriate analog for extinct ecomorphology.

The fossil *Pseudomyrmex* macrops specimen used as a proof-of-concept was consistently and accurately predicted as a lignicolous arboreal-foraging phytophagivore (Figure 2). The arboreal foraging niche was predicted with the highest confidence, while the phytophagous functional role was predicted with the lowest confidence (Tables S7-S22). This specimen was preserved in Dominican amber, approximately 16 Ma, and measurements were taken from a microCT scan: the accuracy of the model prediction suggests that neither taphonomy nor microCT scan-based measurements distort morphology in prediction.

**Figure 1.**
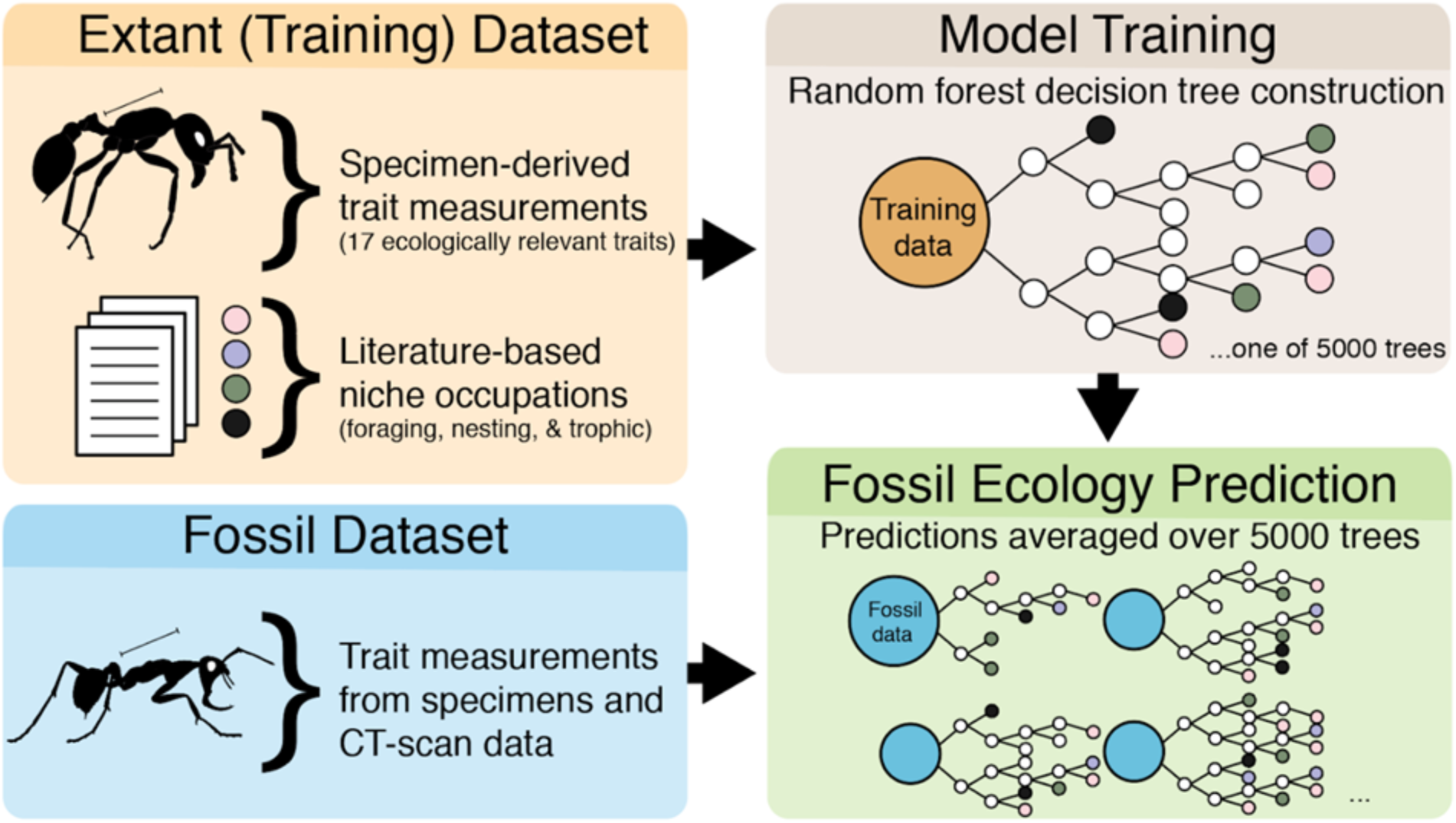
Diagrammatic workflow of predictive model development and testing. A comprehensive morphometric dataset of extant ants was compiled; species were binned according to various ecological niche aspects based on surveys of the literature. Random Forest models were then trained on subsets of the original dataset. Homologous traits were measured on fossil ant specimens; when available, traits were measured from CT reconstructions, and otherwise were measured under light microscopy. Finally, the pre-trained Random Forest models were used to predict extinct ecology from fossil morphometric datasets.

**Figure 2.**
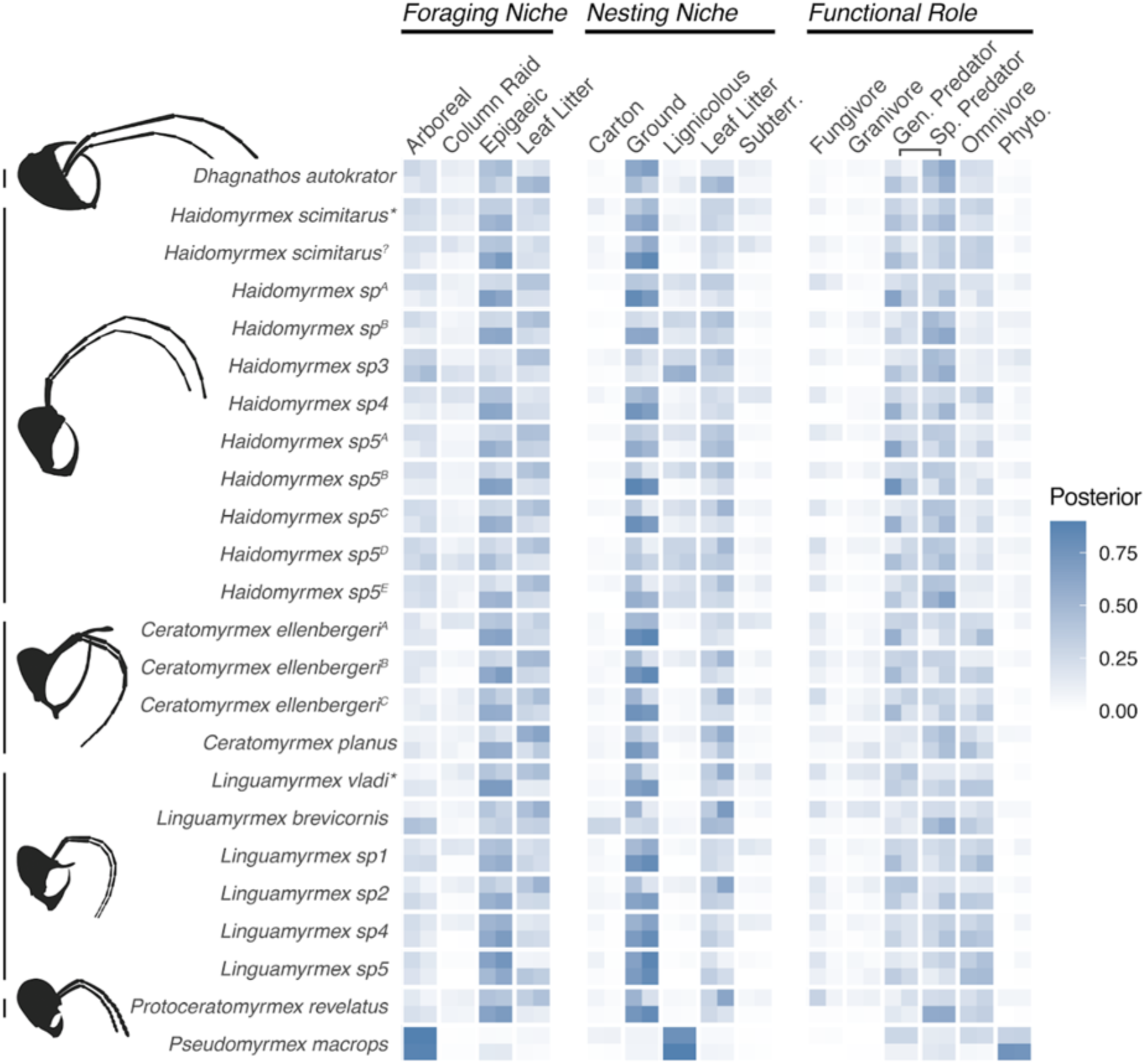
Posterior probabilities of niche aspect predictions. Each species’ niche aspect predictions are represented by four models derived from alternate datasets; from top left, clockwise: raw functional measurements, raw homologous measurements, size-corrected ratio functional measurements, size-corrected ratio homologous measurements. Taxonomic sampling includes described (named) taxa as well as putative morphospecies. Alternate specimens of the same species are denoted with superscripts, *denotes specimens included through CT-scan reconstruction data. All posterior probabilities are available in Supplemental Tables S7-38; full specimen information is available in Supplemental Table 6.

Hell ants are primarily recovered as epigaeic foragers that nested directly in the ground, though several species are predicted as leaf litter nesters and foragers (Figure 2). Additionally, one *Haidomyrmex* morphospecies *(Haidomyrmex sp3)* was partially predicted as a lignicolous nester and arboreal forager; with *Linguamyrmex brevicornis* also partially predicted as an arboreal forager. Hell ants were primarily predicted as predators, both specialist and generalist, though some species were additionally predicted to be omnivorous (Figure 2, Supplemental Tables S7-S38). Given the ecological plasticity in many extant ant species - many species are primarily predaceous but may supplement their diet with plant material - we may expect hell ants to follow this same pattern. Additionally, even when species were predicted as omnivores, the probabilities of specialist predation and generalist predation when summed were greater than the probability of omnivory, suggesting that atomization of predator classes may have played some role in the prediction of omnivory. Supporting the general accuracy of our models, we find broad congruence across species and also do not recover any strong predictions of unlikely ecological niches: hell ants were not predicted as subterranean nesters, column-raiding foragers, or fungivorous, granivorous, or phytophagous functional roles (Figure 2).

Posterior probabilities for each prediction (an indicator of Random Forest model vote consensus) were highest with foraging niche and nesting niche predictions, indicating overall greater confidence in these predictions (Figure 2, Supplemental Tables S7-S38). Predictions for functional role and ecomorph had lower posterior probabilities (Figure 2, Supplemental Tables S7-S38). This may be a function of a greater number of potential aspects for each (six possible functional roles and ten possible ecomorphs), necessarily splitting the model votes. Extant morphology may also have less predictive power for extinct ecology with respect to functional role and ecomorph, resulting in lower predictive accuracy.

Intraspecific consensus among the four models used was variable. For some specimens, there was very strong agreement, with all four models predicting the same ecological niche aspect binning; however, there were also cases where two models predicted one niche aspect binning, and two predicted another (Figure 2, Supplemental Tables S7-S38). There were rarely scenarios where more than two niche aspect binnings were predicted for a single specimen. More frequently in situations of evenly split predictions, the size-corrected ratio models would predict the same aspect binning while the raw measurement models would predict another; this suggests that the difference between functional morphology and homologous morphology with respect to eye positioning did not matter as much to niche prediction. This may be because the model ranked eye positioning traits relatively low on its scale of variable importance, thus, the difference between functional and homologous morphology was negligible (Supplemental Figure S4, S5).

There was a limited drop in accuracy from the models trained on the full morphometric dataset to the models trained on the limited morphometrics dataset, with missing traits removed (Table S5). In most cases, the difference was only a few percentage points, at most five percent. This limited decrease in accuracy suggests that the traits removed were not essential to be measured, and that there is a core set of most relevant traits. Variable importance plots from the models rank traits related to eyes, antennae, legs, and mandibles as most crucial to model accuracy (Supplemental Figures S5, S6). The traits that were removed primarily dealt with dimensions of the head and thorax; while their inclusion improves model accuracy, it is not necessary to include all of them in order to obtain a reasonably accurate prediction of extinct ecological niches.

We found no robust differences in niche predictions between specimens measured directly and microCT-measured specimens; specimens measured from microCT scans were also predicted as epigaeic or leaf litter foragers and nesters (Figure 2, Tables S7-S38). Additionally, we find that the dealate *Haidomyrmex scimitarus* (measured from a microCT scan) and the worker *Haidomyrmex scimitarus*(measured through light microscopy) were both predicted to be ground-nesting epigaeic predators (Figure 2, Supplemental Tables S7-S38); illustrating consensus between the two types of input data. This congruent ecological assignment also supports the hypothesis that stem-ant queens did not establish nests through claustral founding – queens of many extant species do not actively forage themselves – but occupied the same general ecological niche as workers of their species (Barden & Grimaldi 2012).

### Niche occupation in extinct and extant specialized predators

Our most conservative estimates of ecological niche occupation suggest that hell ants occupied primarily ground-nesting epigaeic niches with some leaf litter occupation, while across-model results recover hell ants within arboreal, ground, and leaf litter niches across a moderate body size range. In comparing predicted hell ant ecospace to solitary predator ant ecospace, we find that hell ants occupied at least part of the ecomorphological spaces occupied by extant lineages (Figure 3). The sister ponerine genera *Anochetus* and *Odontomachus* exhibit the greatest extant ecospace diversity and disparity, occupying most potential ecospace, with species ranging from ~3mm to ~1.7cm spanning arboreal, ground, and leaf litter niches (Brown 1978; Hoenle et al. 2020). The most restricted ecospace is occupied by species within the formicine genus Myrmoteras, which are minute leaf litter dwellers.

**Figure 3.**
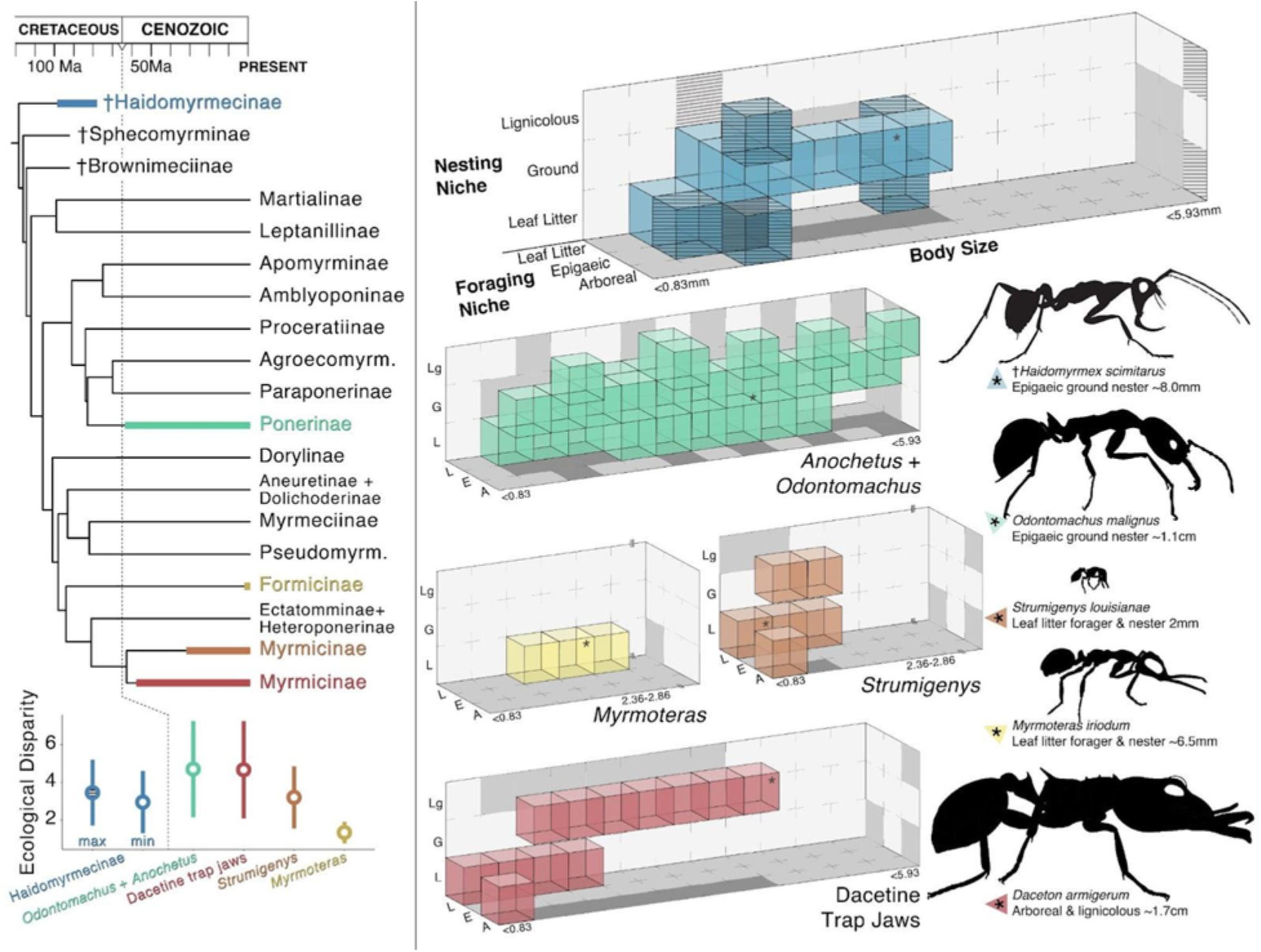
Ecospace occupation of haidomyrmecine and extant specialist predator lineages. (Top left) Subfamily-level time-calibrated phylogeny of ants with divergence dates from Borowiec et al. (2019), Haidomyrmecinae and extant lineages denoted with colored bars; haidomyrmecine range derived from oldest and youngest deposit ages, extant ranges based on available crown age estimations for each lineage: Ponerinae – *Anochetus*+*Odontomachus* (Fernandes et al. 2021); Myrmicinae – *Strumigenys*(Booher et al. 2021); Myrmicinae - dacetine trap-jaws (Ward et al. 2015). No divergence date estimates are available for the formicine trap-jaw genus *Myrmoteras.*(Right) Lineage-specific ecomorphological niche occupations. Each colored cube represents a unique occupied niche. Hashed cubes in haidomyrmecine ecospace indicate “maximum” hypothetical niche occupation based on all unique combinations estimated across all four random forest models while remaining cubes reflect only the majority aspect from the raw functional measurement model. All extant ecospaces compiled from literature. (Bottom left) Within-lineage disparity calculated as average pairwise distance between each unique three-dimensional occupation. Maximum and minimum haidomyrmecine values represent alternate niche occupations described in the right panel. Disparity values are listed in Table S39.

Ecological disparity does not appear to be linked to species diversity per se: the most taxonomically diverse trapjaw lineage is *Strumigenys* with over 850 species, but the constrained size and primarily leaf litter habits of the genus produce a within-group ecological disparity that is low relative to *Anochetus* and *Odontomachus* with 188 total species (Figure 3) (Bolton 2021). Similarly, while total species diversity of hell ants is unknown, even within our limited fossil sample, haidomyrmecines are found to be relatively ecologically disparate and diverse compared to extant lineages.

## Discussion

### Hell ants as epigaeic or leaf litter predators

We recover broad consensus across models for hell ant ecological niche occupations: models consistently predicted haidomyrmecine taxa as leaf litter foraging or epigaeic ground-nesting predators, with few outliers (Figure 4). Variations within the testing datasets, such as homologous vs functional morphology or missing traits, did not greatly affect predictions. Our results are in contrast to previous hypotheses suggesting a primarily arboreal lifestyle among hell ants. Initial hypotheses were based on qualitative assessments of morphology (Barden & Grimaldi 2012) and an assertion that hell ants’ vertically aligned mandibles might have precluded the fine manipulation of soil required to create ground nests (Dlussky 1996). However, extant and fossilized behavioral evidence provide support for ground and leaf litter leaf litter habits among in haidomyrmecines. Many extant trap-jaw ant species are capable of manipulating soil with their highly specialized mandibles, allowing for nesting directly in the ground (Cerquera & Tschinkel 2010). Soil nesting is also estimated as the ancestral state among all crown ants, although fossils have not yet been included in such reconstructions (Lucky et al. 2013). Additionally, two fossilized examples of hell ant prey reflect leaf litter and surficial habitats: a beetle larva in association with a *Linguamyrmex vladi* worker (Barden et al. 2017), likely reflecting a humid leaf litter habitat; and a cockroach relative *Caputoraptor elegans* in association with *Ceratomyrmex ellenbergi* (Barden et al. 2020), possibly living in leaf litter or surficial areas, although arboreal habitats have been proposed (Bai et al. 2018). This reconstructed nesting ecology aligns with proposed “extrinsic factors” related to the evolution of eusociality (Evans 1977).

**Figure 4.**
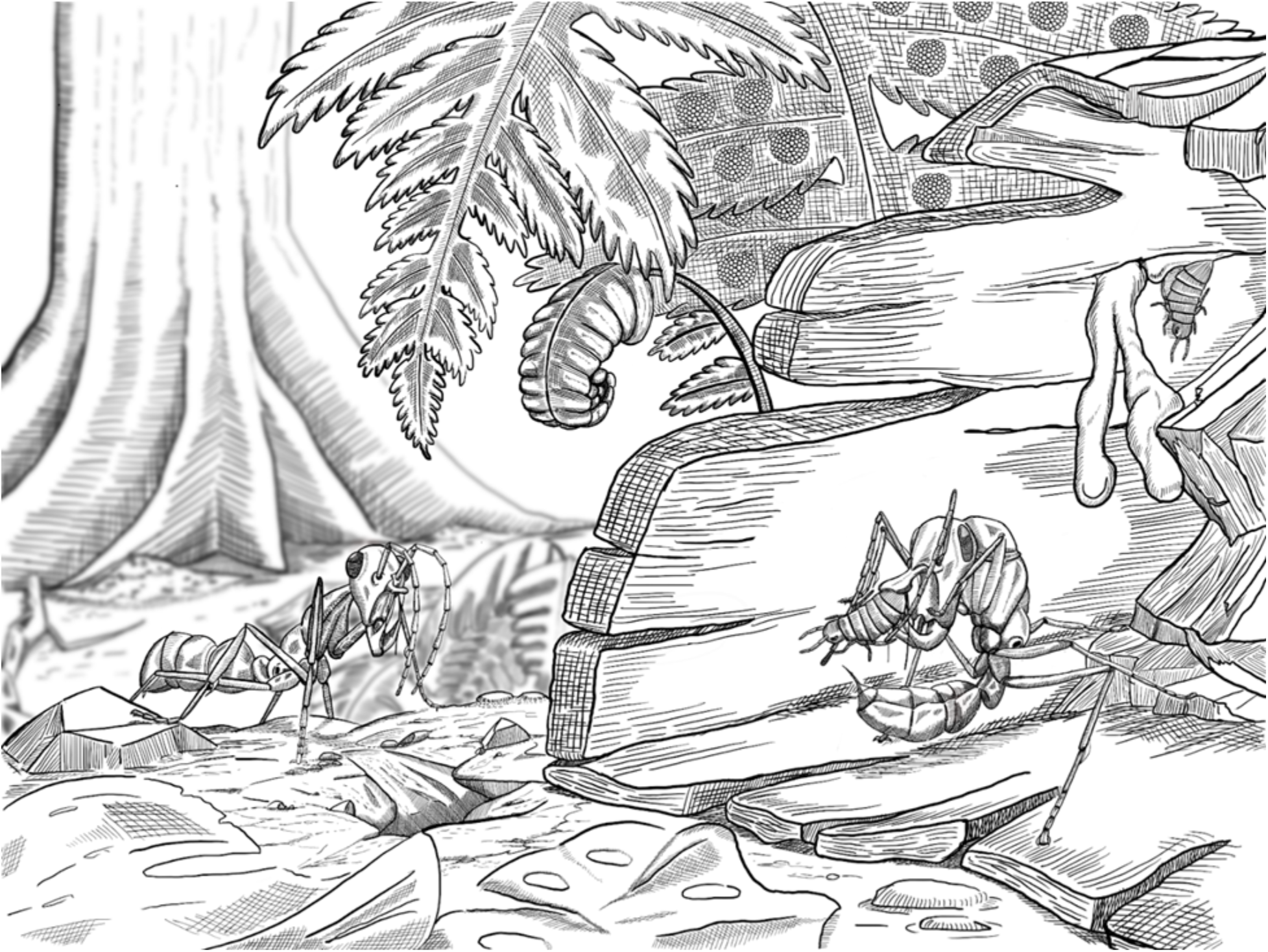
Reconstruction of the putative nesting and predatory foraging habits of the hell ant *Linguamyrmex vladi.* Artist: John Paul Timonera.

Entrapment bias of ants in amber tends to skew towards arboreal species, though preservation is contingent on the position of resin secretions on the source plant (Solórzano-Kraemer et al. 2015, 2018). Experiments conducted using sticky traps as a resin analogue found that sticky traps placed higher up in the trees collected arboreal species, while traps placed within one meter of the ground collected leaf litter species, possibly foraging for prey caught in the traps (Solórzano-Kraemer et al 2018). While we do not recover hell ants primarily as arboreal species, it is possible that hell ants foraging around the base of trees, perhaps attracted by struggling prey, would have themselves become entrapped in amber. This may be similar to fossilization in asphalt and tar traps, where predatory species are overrepresented because they are attracted to dying prey animals trapped in the tar (Stock 2001). Considering the broad diversity of specimens entrapped in Cretaceous amber (Sánchez-García and Engel 2016; Selden and Ren 2017; Stoev et al. 2019; Yu et al. 2019; Ngô-Muller et al. 2020), it is likely that at least some amber secretions occurred lower on trunks or root buttresses, entrapping ecologically diverse segments of the overall amber biota. Additionally, evidence from Cenozoic amber ant assemblages finds that ground-dwelling ant species are frequently entrapped in resin, lending further support to the likelihood of entrapment for non-arboreal hell ants (Guénard et al. 2015).

### Ecospace occupation and ecological succession

Many extant predatory ants hunt in groups, however some lineages capture prey alone (Beckers et al. 1989) and certain morphological adaptations necessitate solo foraging. Workers of extant trap-jaw ants capture prey through rapid closure of their specialized mandibles – often a power-amplified mechanism is employed following the activation of elongate trigger setae in the path of mandible movement. This specialized prey capture typically precludes group predation; workers individually subdue a single prey item before returning to the nest. An instance of preserved predation has demonstrated that hell ants captured prey alone in a manner analogous to modern trap-jaw taxa (Barden et al. 2020). Hell ant workers pinned prey between their vertically-expanded tusk-like mandibles and cranial appendages. Our results suggest that hell ants may have been functional equivalents to many modern-day trapjaws in surficial and leaf litter arthropod communities; solitary foraging hunters seeking out prey across the forest floor and in interstitial leaf litter spaces. While there are some morphological traits that support a trap-jaw mechanism in hell ants, including trigger hairs and a structurally-reinforced clypeal paddle at the point of mandible articulation, it is unknown whether or not hell ants were trap-jaw predators. Nevertheless, they were most likely solitary-foraging predators, considering fossil paleoethological evidence and consistent individual fossilization of specimens, in contrast to worker aggregations known in other stem-ant genera (Barden & Grimaldi 2016; Barden et al. 2020).

We recover a pattern of repeated ecological niche occupation across lineages of solitary-foraging specialized ant predators. Even as molecular-based divergence estimates place the origin of crown ants during the Cretaceous (Borowiec et al. 2019; Moreau et al. 2006), the earliest extant trap-jaw predators originated in the Cenozoic (Figure 3) (Booher et al. 2021; Fernandes et al. 2021; Ward et al. 2015). Molecular divergence-date estimates for crown-group origins of trap-jaw lineages range from ~65 Ma in ponerine ants (Fernandes et al. 2021) to ~35 Ma in the myrmicine genus Strumigenys (Booher et al. 2021). The last hell ant fossil dates to 78 Ma in Campanian age Canadian amber (McKellar et al. 2013). It is unclear precisely when hell ants went extinct, however it seems plausible that a mass extinction event may have contributed to their demise. The overlap of ecospace occupation between hell ants and the lineages arising soon after the Cretaceous-Paleogene extinction (KPg), ponerine and dacetine trap jaw ants, suggests faunal turnover in niche occupation. The ecological breadth of modern trap jaw ants may represent echoes of their Cretaceous counterparts.

Our reconstructed hell ant ecospace overlaps with occupations of all extant groups sampled. Hell ants occupied approximately 10% of total potential sampled ecomorphospace and radiated into arboreal, leaf litter, and surficial habitats. Subsequently, other specialized predatory ant lineages radiated into ecomorphospace that partially overlaps with hell ant ecomorphospace. Trap-jaw dacetines and ponerines emerged relatively rapidly after the Cretaceous end extinction, while other trap-jaw lineages emerged later in the Cenozoic; however, all lineages occupy at least part of the hell ant ecomorphospace. This pattern illustrates potential ecological restriction of solitary specialized predators. Additionally, while we included most known species in extant ecospace reconstructions, our fossil sampling was much more limited; thus the full ecospace occupation of hell ants was probably more broad than our current reconstruction suggests.

### Pitfalls and potential of paleoecological niche reconstructions

We present here a pipeline for paleoecological niche prediction using machine learning algorithms and broad ecomorphological sampling. To our knowledge, this is the first attempt at predicting extinct ecology using Random Forest. Because this class of supervised machine learning incorporates non-linear modeling and has been shown to outperform various other discriminant function methods (Pigot et al. 2020; Sosiak & Barden 2021), it represents a powerful tool in reconstruction of fossil niche occupations and communities. The method requires a taxonomic group with both fossil and extant representatives, which exhibit consistent body plans, allowing for homologous trait measurements across time series. The application of extant trait data in extinct ecological estimation is best suited among lineages that exhibit a high degree of extant diversity relative to fossil samples: the more ecologies that the extant group occupies, the more likely it will be that all potential ecological niches of the extinct group are represented.

Predicted ecologies may also be evaluated in the context of other fossil evidence. For example, several extant ant taxa are primarily granivorous seed-eaters (Cole 1968; Plowes et al. 2013), thus the functional role of extinct species could potentially be predicted as granivorous. However, while grasses first evolved in the early Cretaceous, grassland ecosystems did not develop broadly until later in the Cenozoic, making it unlikely that Mesozoic ants would have been granivorous (Stromberg 2011; Boyce and Lee 2017). Indeed, while our model allowed for six functional role binnings, haidomyrmecines were never predicted as belonging to temporally inappropriate ecologies with any certainty, even as our included Cenozoic fossil specimen was correctly estimated as an uncommon phytophagivore.

Quantitative predictions of ecological niches allow for new approaches to many paleoecological questions. We present here one such application; a relative comparison of ecospace occupation between extinct and extant lineages. By reconstructing the ecological community of the earliest ants, we find repeated lineage occupation of ecospace, consistent with dynastic succession across Earth’s last mass extinction event.

## Supporting information

Supplementary Data: Fossil Morphometrics

Supplementary Data: Specimen Images

Supplementary Data: Trap Jaw Morphospace

Supplementary Data: Random Forest Code

## Acknowledgements

We thank Christine Johnson and Christine Lebeau for facilitating access to specimens at the American Museum of Natural History, in addition to Morgan Hill and Andrew Smith for facilitating access to imaging equipment at the AMNH; Stefan Cover and David Lubertazzi for facilitating specimen access and hospitality at the Museum of Comparative Zoology; Eugenia Okonski and Ted Schultz for facilitating specimen access and hospitality at the Smithsonian National Museum of Natural History; all of whom without which we would not have been able to develop the initial model.

## Data and materials availability

All data and scripts needed to reproduce the analyses and evaluate the conclusions in the paper are present in the paper and/or in the Supplementary Materials.

## Contributions

Conceptualization: CES, PB Methodology: CES, PB

Data compilation and collection: CES, VP, TJ

Data visualization: CES, PB, JPT

Writing–original draft: CES, PB

Writing–review & editing: CES, PB, VP, TJ, JPT

## Ethics

The fossil specimens used in this research are preserved primarily within Burmese amber. We affirm that all specimens were acquired prior to June 2017, pursuant to the proposed boycott of Burmese amber by the Society of Vertebrate Paleontologists (Rayfield et al. 2020). Some specimens are located in a private collection, while others are located in institutional museums: specimen repositories are indicated in the Supplementary Information associated with this manuscript, and we have provided photomicrographs of all specimens residing in a private collection. All data associated with this manuscript are available as Supplementary Information, including all morphometric measurements for privately-owned specimens.

