## Supplemental Information for "Trait-based paleontological niche prediction demonstrates deep time parallel ecological occupation in specialized ant predators"

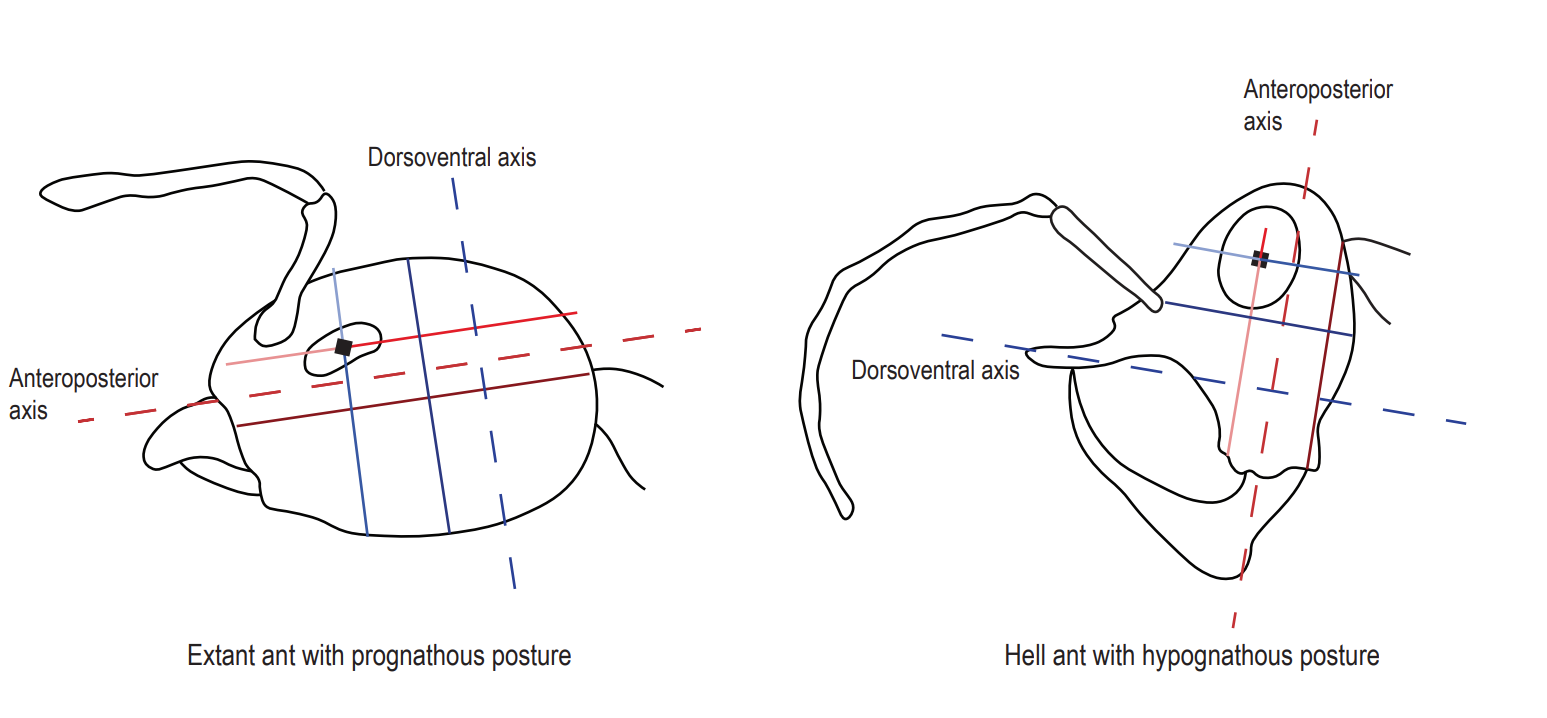

**Figure S1.**

Eye positioning measurements consistent with homologous morphology.

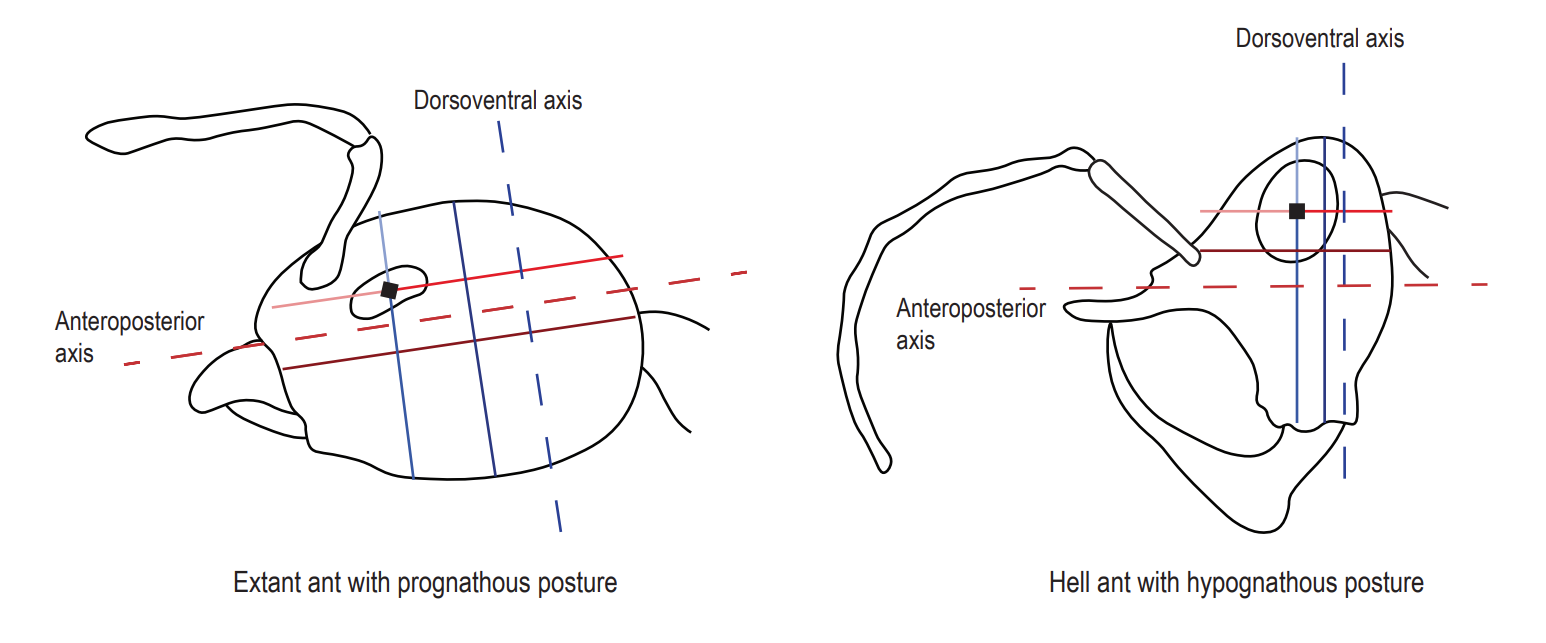

**Figure S2.**

Eye positioning measurements consistent with functional morphology.

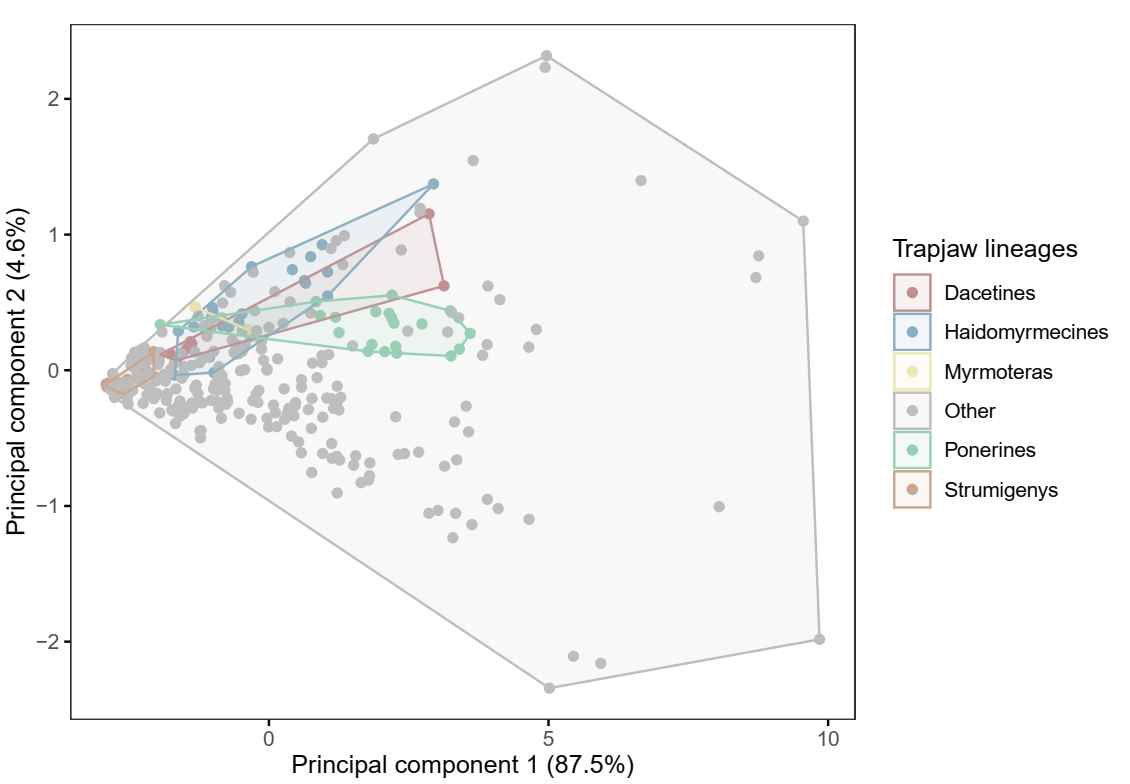

**Figure S3.**

Principal component analysis of extant and extinct morphospace using raw trait measurements. The morphospace of each lineage with an independent origin of trap-jaw mechanics is delineated.

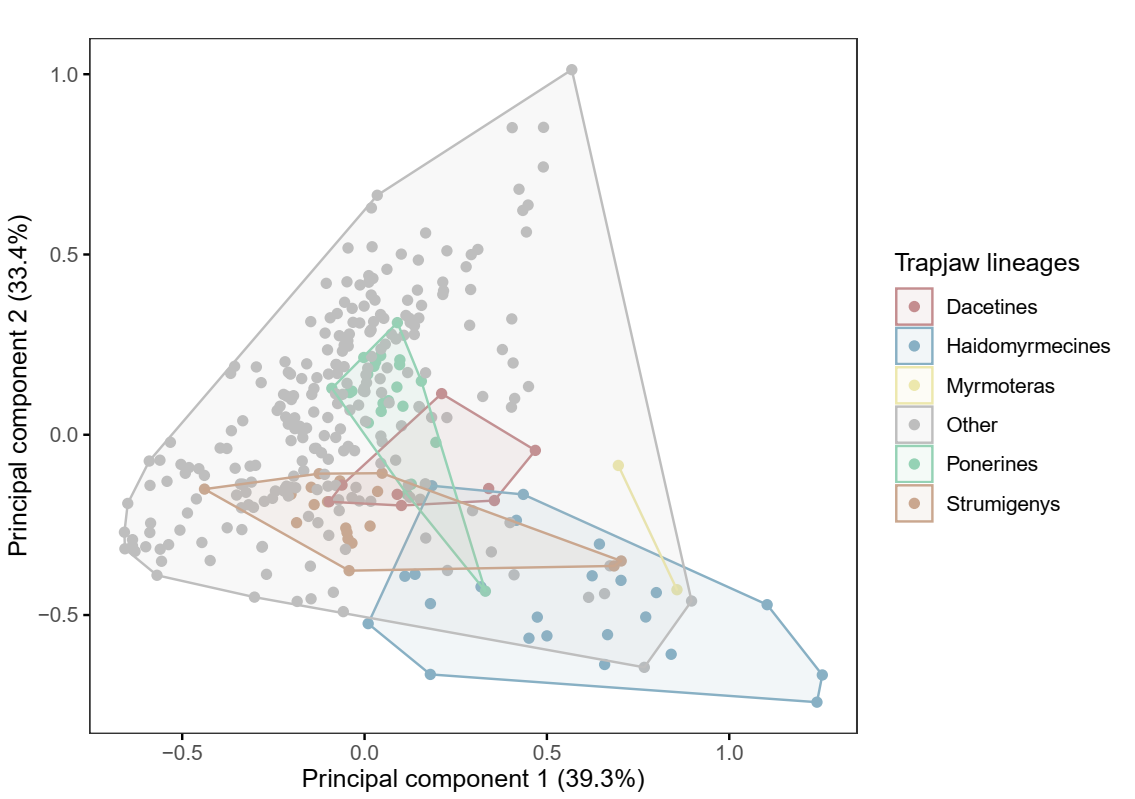

**Figure S4.**

Principal component analysis of extant and extinct morphospace using size-corrected ratio measurements. The morphospace of each lineage with an independent origin of trap-jaw mechanics is delineated.

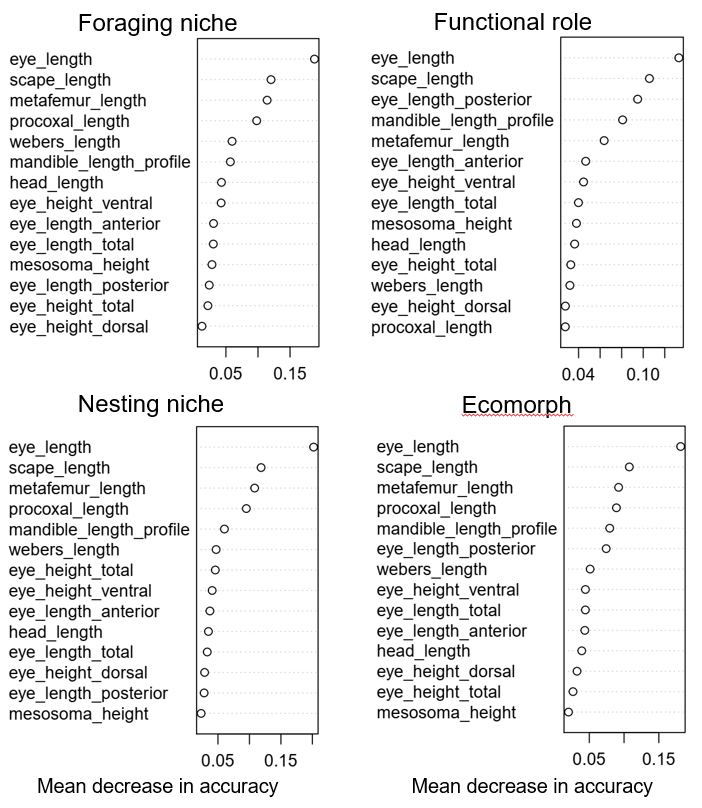

**Figure S5.**

Variable importance plots for each Random Forest classification model (functional role; nesting niche; foraging niche; ecomorph syndrome) using raw morphological trait measurement data. Variable importance is calculated as the mean decrease in accuracy across splits when the variable was eliminated; a higher decrease in accuracy when that variable was omitted indicates greater importance to the model.

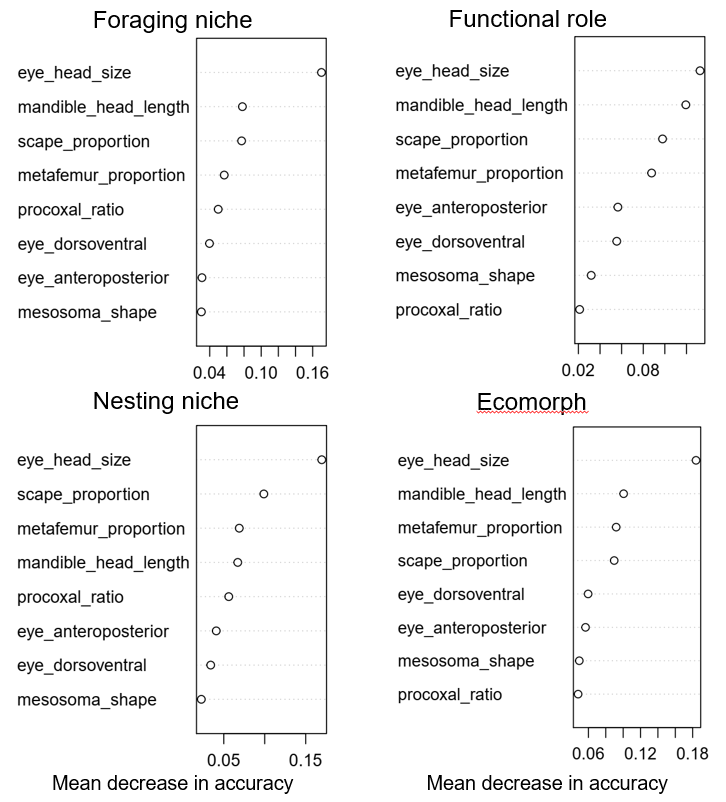

**Figure S6.**

Variable importance plots for each Random Forest classification model (functional role; nesting niche; foraging niche; ecomorph syndrome) using size-corrected ratio measurement data. Variable importance is calculated as the mean decrease in accuracy across splits when the variable was eliminated; a higher decrease in accuracy when that variable was omitted indicates greater importance to the model.

| **Trait** | **Description of measurement** | **Known ecological significance** |
| --- | --- | --- |
| Head width (HW) | Taken in frontal view along widest axis of head capsule excluding eyes | Mandibular musculature of workers (Kaspari 1993); size of spaces workers can move through (Sarty et al. 2006) |
| Head length (HL) | Medially from anterior margin of clypeus to vertex of head capsule in frontal view | Size of spaces workers can move through (Kaspari and Weiser 1999) |
| Eye length (EL) | Measured along longest axis of eye | Foraging behaviour and foraging period (Weiser and Kaspari 2006) |
| Mandible length (lateral profile view) (MLP) | From point of insertion to apical-most tooth of mandible | Diet (Fowler et al. 1991) |
| Anteroposterior eye position (3 measurements taken) (LHL; ELA; ELP) | Taken in lateral view: length from midpoint of eye to anterior clypeal margin (ELA); length of head from midpoint of eye to posterior margin (ELP); total lateral head length (LHL) used to calculate eye position ratios | Foraging and diet (Fowler et al. 1991); habitat stratum (Gibb and Parr 2013) |
| Dorsoventral eye position (3 measurements taken) (HH; EHD; EHV) | Taken in lateral view: height of head from midpoint of eye to dorsal margin of head (EHD); height of head from midpoint of eye to ventral margin of head (EHV); total head height (HH) used to calculate eye position ratios | Related to foraging and diet (Fowler et al. 1991); habitat stratum occupied (Gibb and Parr 2013) |
| Mandible length (frontal view) (MLF) | From point of clypeal insertion to apical-most tooth of mandible | Diet (Fowler et al. 1991) |
| Scape length (SL) | From antennal socket to distal margin of scape | Chemosensory; detection of pheromone trails (Weiser and Kaspari 2006) |
| Weber’s length (WL) | Taken in lateral view from the anterodorsal margin of the pronotum to the posteroventral margin of the mesosoma | Established proxy for worker body size (Weber 1938) |
| Procoxal length (PL) | From articulation point with propleuron to the distal tip of the procoxa | N/A |
| Mesosoma height (MH) | Taken at a right angle to Weber’s length from the ventral margin of propleuron to dorsal margin of pronotum | N/A |
| Pronotal width (PW) | Measured at the widest point of the pronotum when viewed dorsally | Body mass of workers (Kaspari and Weiser 1999) |
| Metafemur length (ML) | Measured from articulation point with trochanter to distal tip of the metafemur | Foraging speed and habitat complexity (Feener et al. 1988) |

**Table S1.**

All morphological traits measured, with description of measurements taken, and known ecological significance of traits.

| **Trait** | **Formula** | **Interpretation** |
| --- | --- | --- |
| Head shape (HS) | HW/HL | Larger value = more boxy, square head  Smaller value = more elongate along anteroposterior axis |
| Procoxal proportion (PP) | PL/WL | Larger value = procoxa longer in comparison to body length  Smaller value = procoxa shorter in comparison to body length |
| Eye proportion (EP) | EL/HL | Larger value = eyes large in comparison to head  Smaller value = eyes small in comparison to head |
| Pronotum expansion (PE) | PW/WL | Larger value = pronotum laterally expanded in relation to body size  Smaller value = pronotum laterally narrow in relation to body size |
| Pronotum flattening (PF) | PW/MH | Larger value = pronotum broad and dorsoventrally flattened  Smaller value = pronotum narrow and dorsoventrally expanded |
| Mesosoma shape (MS) | MH/WL | Larger value = overall elongate narrow body  Smaller value = overall stocky squarish body |
| Mandible curvature (MC) | MLP/MLF | Larger value = low degree of mandible curvature  Smaller value = high degree of mandible curvature |
| Mandible proportion (MP) | MLP/HL | Larger value = elongate mandibles in comparison with head size  Smaller value = short mandibles in comparison with head size |
| Dorsoventral eye position (DEP) | EHD/HH | Larger value = eye positioned more ventrally  Smaller value = eye positioned more dorsally |
| Anteroposterior eye position (AEP) | ELA/LHL | Larger value = eye positioned more posteriorly  Smaller value = eye positioned more anteriorly |
| Scape proportion (SP) | SL/HL | Larger value = scape elongate in comparison with head size  Smaller value = scape short in comparison with head size |
| Metafemur proportion (MeP) | ML/WL | Larger value = elongate leg in relation to body size  Smaller value = shorter leg in relation to body size |

**Table S2.**

Size-corrected ratios of traits measured, formulae for calculation, and interpretation of ratio value.

| **Ecomorph designation based on granular binnings** | **Functional role binnings collapsed** | **Final ecomorph syndromes** |
| --- | --- | --- |
| fg_gr_eg | om_gr_eg | **Ground or leaf litter-nesting epigaeic omnivore**  om_grll_eg  Exemplars: *Camponotus; Dinoponera; Aphaenogaster* |
| gn_gr_eg |  |  |
| mh_gr_ll | om_gr_ll |  |
| om_gr_eg | om_gr_eg |  |
| om_gr_ab | om_gr_ab |  |
| om_gr_ll | om_gr_ll |  |
| om_ll_eg | om_ll_eg |  |
| gn_leaf | om_leaf | **Leaf litter omnivore**  om_leaf  Exemplars: *Aneuretus; Acanthomyrmex; Cyphomyrmex* |
| om_leaf |  |  |
| om_lg_ab | om_lg_ab | **Lignicolous arboreal omnivore**  om_lg_ab  Exemplars: *Pseudomyrmex; Cataulacus; Cephalotes* |
| py_lg_ab |  |  |
| tr_lg_ab |  |  |
| om_cn_ab | om_cn_ab | **Carton-nesting arboreal omnivore**  om_cn_ab  Exemplars: *Crematogaster; Oceophylla; Azteca* |
| om_cn_eg | om_cn_eg |  |
| om_sub | om_sub | **Subterranean omnivore**  om_sub  Exemplars: *Acropyga* |
| gp_gr_eg | pred_gr_eg | **Ground-nesting epigaeic or leaf litter predator**  pred_gr_egll  Exemplars: *Odontomachus; Diacamma; Harpegnathos* |
| sp_gr_eg | pred_gr_ll |  |
| gp_gr_ll |  |  |
| gp_leaf | pred_leaf | **Leaf litter predator**  pred_leaf  Exemplars: *Hypoponera; Mystrium; Strumigenys* |
| sp_leaf |  |  |
| sp_lg_ab | pred_lg_ab | **Lignicolous arboreal predator**  pred_lg_ab  Exemplars: *Simopone; Cylindromyrmex; Daceton* |
| gp_lg_ab |  |  |
| gp_ll_cr | pred_ll_cr | **Column-raiding predator**  pred_cr  Exemplars: *Simopelta; Neivamyrmex; Cheliomyrmex* |
| gp_sb_cr | pred_sb_cr |  |
| sp_sb_cr |  |  |
| sp_gr_cr | pred_gr_cr |  |
| gp_ll_eg | pred_ll_eg | **Leaf litter-nesting epigaeic predator**  pred_ll_eg  Exemplars: *Leptogenys; Platythyrea* |
| sp_ll_eg |  |  |
| gp_sb_ll | pred_sb_ll | **Subterranean predator**  pred_sub  Exemplars: *Proceratium; Stigmatomma; Leptanilla* |
| sp_sb_ll |  |  |

**Table S3.**

Schematic of ecomorph syndrome collapse: ecomorphs were collapsed based on overlap between groups, confusion/misclassification rates between groups, and qualitative assessment of likely ecological overlap (leaf litter foragers may also forage on the surface of the ground, for example). Final column shows ecomorph syndrome abbreviations, definitions, and exemplar taxa (abbreviations used in raw dataset available on Dryad (DOI: https://doi.org/10.5061/dryad.kh1893243).

| **Binning designator** | **Definition** | **Exemplar taxa** |
| --- | --- | --- |
| ***Functional role*** | | |
| GP | Generalist predator – broad taxonomic diet | *Odontomachus; Diacamma; Harpegnathos* |
| SP | Specialist predator – obligate feeding on specific taxon (e.g. termites) | *Acanthostichus; Megaponera; Simopelta* |
| Om | Omnivorous – prey items, plant matter, etc. | *Paraponera; Camponotus; Iridomyrmex* |
| Py | Phytophagous – extrafloral nectaries, herbivory, etc. | *Pseudomyrmex; Tetraponera; Myrmelachista* |
| Fg | Fungus-growing | *Cyphomyrmex; Trachymyrmex; Atta* |
| Tr | Trophobiotic – symbiotic relationship with other insects (homopteran secretions, etc.) | *Acropyga; Melissotarsus; Rhopalomastix* |
| Gn | Granivorous – seed-harvesting | *Acanthomyrmex; Pogonomyrmex; Veromessor* |
| Mh | Mushroom-foraging | *Euprenolepis* |
| ***Nesting niche*** | | |
| Cn | Carton-nesting – structured nests from plant material in trees and shrubs | *Oecophylla; Azteca; Liometopum* |
| Gr | Ground-nesting – nests in dirt mounds, under stones, rock cracks, etc. | *Platythyrea; Formica; Pheidole* |
| Lg | Lignicolous – nests in twig and tree cavities | *Pseudomyrmex; Simopone; Cylindromyrmex* |
| Ll | Leaf litter-nesting – nests in leaf litter interstitial space, rotten wood, etc. | *Strumigenys; Discothyrea; Typhlomyrmex* |
| Sb | Subterranean-nesting | *Leptanilloides; Leptanilla* |
| ***Foraging niche*** | | |
| Ab | Arboreal – in and on trees and shrubs | *Daceton; Tetraponera; Crematogaster* |
| CR | Column-raiding – cooperative, nomadic or raiding predation | *Simopelta; Dorylus; Eciton* |
| Eg | Epigaeic – active foraging on the ground surface | *Leptomyrmex; Rhytidoponera; Myrmecocystus* |
| Ll | Leaf litter – within interstitial spaces in leaf litter | *Discothyrea; Amblyopone; Heteroponera* |
| Sb | Subterranean – underground | *Acropyga* |

**Table S4.**

Ecological niche aspect binning abbreviations, definitions, and exemplar taxa (abbreviations used in dataset available on Dryad, DOI: https://doi.org/10.5061/dryad.kh1893243)

| **Complete morphometric dataset** | | | | |
| --- | --- | --- | --- | --- |
|  | Nesting niche | Foraging niche | Functional role | Ecomorph |
| Raw trait measurements | 85.77 | 83.63 | 85.88 | 81.78 |
| Size-corrected ratio trait measurements | 82.92 | 85.05 | 80.92 | 79.84 |
| **Subset of complete morphometric dataset** | | | | |
| Raw trait measurements | 85.41 | 82.92 | 80.53 | 81.78 |
| Size-corrected ratio trait measurements | 80.07 | 81.14 | 82.06 | 77.91 |

**Table S5.**

Out-of-bag (OOB) percent accuracies for each Random Forest model constructed.

| **Species** | **Specimen accession number** | **Caste** | **Sampling method** |
| --- | --- | --- | --- |
| *Ceratomyrmex planus* | BALTJ_025 | Worker | Measured under light microscopy |
| *Ceratomyrmex ellenbergeri* | BALTJ_016 | Worker | Measured under light microscopy |
| *Ceratomyrmex ellenbergeri* | BALTJ_027 | Worker | Measured under light microscopy |
| *Ceratomyrmex ellenbergeri* | BALTJ_029 | Worker | Measured under light microscopy |
| *Protoceratomyrmex revelatus* | BALTJ_043 | Worker | Measured under light microscopy |
| *Linguamyrmex sp5* | BALTJ_050 | Worker | Measured under light microscopy |
| *Linguamyrmex sp4* | BALTJ_031 | Worker | Measured under light microscopy |
| *Linguamyrmex brevicornis* | BALTJ_030 | Worker | Measured under light microscopy |
| *Linguamyrmex sp2* | BALTJ_059 | Worker | Measured under light microscopy |
| *Linguamyrmex sp1* | BALTJ_056 | Worker | Measured under light microscopy |
| *Haidomyrmex sp3* | BALTJ_022 | Worker | Measured under light microscopy |
| *Haidomyrmex sp5* | BALTJ_017 | Worker | Measured under light microscopy |
| *Haidomyrmex sp5* | BALTJ_009 | Worker | Measured under light microscopy |
| *Haidomyrmex sp5* | BALTJ_066 | Worker | Measured under light microscopy |
| *Haidomyrmex sp5* | BALTJ_007 | Worker | Measured under light microscopy |
| *Haidomyrmex scimitarus* | BALTJ_004 | Worker | Measured under light microscopy |
| *Haidomyrmex sp5* | BALTJ_011 | Worker | Measured under light microscopy |
| *Haidomyrmex sp4* | BALTJ_003 | Worker | Measured under light microscopy |
| *Haidomyrmex indet1* | BALTJ_010 | Worker | Measured under light microscopy |
| *Haidomyrmex indet2* | BALTJ_065 | Worker | Measured under light microscopy |
| *Linguamyrmex vladi* | BuPH-1 | Worker | Measured from micro-CT reconstruction |
| *Haidomyrmex scimitarus* | Bu-FB80 | Dealate queen | Measured from micro-CT reconstruction |
| *Pseudomyrmex c.f. macrops* | DR-14-1021 | Worker | Measured from micro-CT reconstruction |
| *Dhagnathos autokrator* | IGR.BU-003 | Alate queen | Measured under light microscopy |

**Table S6.**

List of all fossil specimens included in the study, with caste and measurement type information included. All specimens with accession numbers beginning in BALTJ are from a private collection, while specimens with alternate accession numbers are housed at the American Museum of Natural History (*Linguamyrmex vladi* and *Haidomyrmex scimitarus*); the New Jersey Institute of Technology (*Pseudomyrmex macrops*); and at the University of Rennes (*Dhagnathos autokrator*).

| **Species** | **Cn** | **Gr** | **Lg** | **Ll** | **Sb** | **Predicted aspect** |
| --- | --- | --- | --- | --- | --- | --- |
| *Linguamyrmex vladi* | 0.0322 | 0.502 | 0.0182 | 0.35 | 0.0976 | Gr |
| *Haidomyrmex scimitarus* | 0.136 | 0.3546 | 0.0466 | 0.3012 | 0.1616 | Gr |
| *Pseudomyrmex macrops* | 0.0892 | 0.0846 | 0.7778 | 0.0416 | 0.0068 | Lg |
| *Dhagnathos autokrator* | 0.0424 | 0.6124 | 0.0658 | 0.1712 | 0.1082 | Gr |

**Table S7.**

Nesting niche prediction posterior probabilities when using model incorporating full morphometric dataset; raw trait measurements; functional morphology.

| **Species** | **Ab** | **CR** | **Eg** | **Ll** | **Predicted aspect** |
| --- | --- | --- | --- | --- | --- |
| *Linguamyrmex vladi* | 0.075 | 0.1058 | 0.4062 | 0.413 | Ll |
| *Haidomyrmex scimitarus* | 0.2614 | 0.1728 | 0.3334 | 0.2324 | Eg |
| *Pseudomyrmex macrops* | 0.9048 | 0.0074 | 0.0322 | 0.0556 | Ab |
| *Dhagnathos autokrator* | 0.2752 | 0.1276 | 0.43 | 0.1672 | Eg |

**Table S8.**

Foraging niche prediction posterior probabilities when using model incorporating full morphometric dataset; raw trait measurements; functional morphology.

| **Species** | **Fg** | **Gn** | **GP** | **Om** | **Py** | **SP** | **Predicted aspect** |
| --- | --- | --- | --- | --- | --- | --- | --- |
| *Linguamyrmex vladi* | 0.2072 | 0.1658 | 0.3222 | 0.1426 | 0.0072 | 0.155 | GP |
| *Haidomyrmex scimitarus* | 0.0994 | 0.0576 | 0.304 | 0.277 | 0.0062 | 0.2558 | GP |
| *Pseudomyrmex macrops* | 0.018 | 0 | 0.2766 | 0.2616 | 0.3034 | 0.1404 | Py |
| *Dhagnathos autokrator* | 0.043 | 0.0496 | 0.1808 | 0.232 | 0.0114 | 0.4832 | SP |

**Table S9.**

Functional role prediction posterior probabilities when using model incorporating full morphometric dataset; raw trait measurements; functional morphology.

| **Species** | **om_cn_ab** | **om_grll_eg** | **om_leaf** | **om_lg_ab** | **pred_cr** | **pred_gr_egll** | **pred_leaf** | **pred_lg_ab** | **pred_ll_eg** | **pred_sub** | **Predicted aspect** |
| --- | --- | --- | --- | --- | --- | --- | --- | --- | --- | --- | --- |
| *Linguamyrmex vladi* | 0.0416 | 0.4052 | 0.085 | 0.0102 | 0.0718 | 0.0836 | 0.2484 | 0.0122 | 0.0352 | 0.0068 | om_grll_eg |
| *Haidomyrmex scimitarus* | 0.096 | 0.338 | 0.015 | 0.01 | 0.0656 | 0.1426 | 0.1952 | 0.013 | 0.1214 | 0.0032 | om_grll_eg |
| *Pseudomyrmex macrops* | 0.159 | 0.054 | 0.0154 | 0.4104 | 0.0104 | 0.0078 | 0.048 | 0.287 | 6.00E-04 | 0.0074 | om_lg_ab |
| *Dhagnathos autokrator* | 0.0144 | 0.248 | 0 | 0.0102 | 0.0178 | 0.4646 | 0.1218 | 0.1052 | 0.0174 | 0 | pred_gr_egll |

**Table S10.**

Ecomorph prediction posterior probabilities when using model incorporating full morphometric dataset; raw trait measurements; functional morphology.

| **Species** | **Cn** | **Gr** | **Lg** | **Ll** | **Sb** | **Predicted aspect** |
| --- | --- | --- | --- | --- | --- | --- |
| *Linguamyrmex vladi* | 0.0232 | 0.1886 | 0.03 | 0.5856 | 0.1726 | Ll |
| *Haidomyrmex scimitarus* | 0.039 | 0.4696 | 0.0574 | 0.2966 | 0.1374 | Gr |
| *Pseudomyrmex macrops* | 0.0924 | 0.084 | 0.7782 | 0.04 | 0.0054 | Lg |
| *Dhagnathos autokrator* | 0.0088 | 0.68 | 0.0474 | 0.1728 | 0.091 | Gr |

**Table S11.**

Nesting niche prediction posterior probabilities when using model incorporating full morphometric dataset; raw trait measurements, homologous morphology.

| **Species** | **Ab** | **CR** | **Eg** | **Ll** | **Predicted aspect** |
| --- | --- | --- | --- | --- | --- |
| *Linguamyrmex vladi* | 0.0528 | 0.158 | 0.3324 | 0.4568 | Ll |
| *Haidomyrmex scimitarus* | 0.2258 | 0.1488 | 0.334 | 0.2914 | Eg |
| *Pseudomyrmex macrops* | 0.9014 | 0.0084 | 0.0304 | 0.0598 | Ab |
| *Dhagnathos autokrator* | 0.2124 | 0.1082 | 0.4696 | 0.2098 | Eg |

**Table S12.**

Foraging niche prediction posterior probabilities when using model incorporating full morphometric dataset; raw trait measurements, homologous morphology.

| **Species** | **Fg** | **Gn** | **GP** | **Om** | **Py** | **SP** | **Predicted aspect** |
| --- | --- | --- | --- | --- | --- | --- | --- |
| *Linguamyrmex vladi* | 0.055 | 0.1854 | 0.3942 | 0.1782 | 0.0268 | 0.1604 | GP |
| *Haidomyrmex scimitarus* | 0.019 | 0.0436 | 0.2066 | 0.3202 | 0.0476 | 0.363 | SP |
| *Pseudomyrmex macrops* | 0.0188 | 2.00E-04 | 0.2766 | 0.2478 | 0.313 | 0.1436 | Py |
| *Dhagnathos autokrator* | 0.0248 | 0.0418 | 0.0614 | 0.2192 | 0.0254 | 0.6274 | SP |

**Table S13.**

Functional role prediction posterior probabilities when using model incorporating full morphometric dataset; raw trait measurements, homologous morphology.

| **Species** | **om_cn_ab** | **om_grll_eg** | **om_leaf** | **om_lg_ab** | **pred_cr** | **pred_gr_egll** | **pred_leaf** | **pred_lg_ab** | **pred_ll_eg** | **pred_sub** | **Predicted aspect** |
| --- | --- | --- | --- | --- | --- | --- | --- | --- | --- | --- | --- |
| *Linguamyrmex vladi* | 0.0308 | 0.3514 | 0.0834 | 0.0508 | 0.116 | 0.0496 | 0.283 | 0.0048 | 0.0244 | 0.0058 | om_grll_eg |
| *Haidomyrmex scimitarus* | 0.0216 | 0.3756 | 0.0134 | 0.0598 | 0.0558 | 0.112 | 0.2732 | 0.0128 | 0.0708 | 0.005 | om_grll_eg |
| *Pseudomyrmex macrops* | 0.1624 | 0.0568 | 0.0116 | 0.4136 | 0.0142 | 0.0092 | 0.0456 | 0.279 | 2.00E-04 | 0.0074 | om_lg_ab |
| *Dhagnathos autokrator* | 0.0048 | 0.2558 | 0 | 0.0328 | 0.0202 | 0.475 | 0.1918 | 0.0134 | 0.0052 | 0 | pred_gr_egll |

**Table S14.**

Ecomorph prediction posterior probabilities when using model incorporating full morphometric dataset; raw trait measurements, homologous morphology.

| **Species** | **Cn** | **Gr** | **Lg** | **Ll** | **Sb** | **Predicted aspect** |
| --- | --- | --- | --- | --- | --- | --- |
| *Linguamyrmex vladi* | 0.049 | 0.6252 | 0.0074 | 0.2694 | 0.049 | Gr |
| *Haidomyrmex scimitarus* | 0.0266 | 0.5414 | 0.0972 | 0.2838 | 0.051 | Gr |
| *Pseudomyrmex macrops* | 0.0086 | 0.0686 | 0.843 | 0.0678 | 0.012 | Lg |
| *Dhagnathos autokrator* | 0.0184 | 0.4248 | 0.0814 | 0.42 | 0.0554 | Gr |

**Table S15.**

Nesting niche prediction posterior probabilities when using model incorporating full morphometric dataset; size-corrected ratio trait measurements; functional morphology.

| **Species** | **Ab** | **CR** | **Eg** | **Ll** | **Predicted aspect** |
| --- | --- | --- | --- | --- | --- |
| *Linguamyrmex vladi* | 0.1572 | 0.062 | 0.6564 | 0.1244 | Eg |
| *Haidomyrmex scimitarus* | 0.2086 | 0.108 | 0.4724 | 0.211 | Eg |
| *Pseudomyrmex macrops* | 0.8186 | 0.0142 | 0.1376 | 0.0296 | Ab |
| *Dhagnathos autokrator* | 0.1462 | 0.067 | 0.3526 | 0.4342 | Ll |

**Table S16.**

Foraging niche prediction posterior probabilities when using model incorporating full morphometric dataset; size-corrected ratio trait measurements; functional morphology.

| **Species** | **Fg** | **Gn** | **GP** | **Om** | **Py** | **SP** | **Predicted aspect** |
| --- | --- | --- | --- | --- | --- | --- | --- |
| *Linguamyrmex vladi* | 0.037 | 0.0328 | 0.3172 | 0.362 | 8.00E-04 | 0.2502 | Om |
| *Haidomyrmex scimitarus* | 0.0074 | 0.02 | 0.4008 | 0.1708 | 0.0202 | 0.3808 | GP |
| *Pseudomyrmex macrops* | 8.00E-04 | 0.0044 | 0.0866 | 0.051 | 0.7116 | 0.1456 | Py |
| *Dhagnathos autokrator* | 0.0458 | 0.0158 | 0.383 | 0.1612 | 0.0322 | 0.362 | GP |

**Table S17.**

Functional role prediction posterior probabilities when using model incorporating full morphometric dataset; size-corrected ratio trait measurements; functional morphology.

| **Species** | **om_cn_ab** | **om_grll_eg** | **om_leaf** | **om_lg_ab** | **pred_cr** | **pred_gr_egll** | **pred_leaf** | **pred_lg_ab** | **pred_ll_eg** | **pred_sub** | **Predicted aspect** |
| --- | --- | --- | --- | --- | --- | --- | --- | --- | --- | --- | --- |
| *Linguamyrmex vladi* | 0.0684 | 0.2738 | 0.0552 | 6.00E-04 | 0.0558 | 0.4122 | 0.1088 | 0.0058 | 0.0192 | 2.00E-04 | pred_gr_egll |
| *Haidomyrmex scimitarus* | 0.0114 | 0.1738 | 0.0102 | 0.0146 | 0.0308 | 0.4666 | 0.1072 | 0.0624 | 0.1136 | 0.0094 | pred_gr_egll |
| *Pseudomyrmex macrops* | 0.0066 | 0.0354 | 0.0048 | 0.6404 | 0.0072 | 0.0222 | 0.0098 | 0.231 | 0.0406 | 0.002 | om_lg_ab |
| *Dhagnathos autokrator* | 0.0136 | 0.1948 | 0.025 | 0.0136 | 0.0216 | 0.2822 | 0.3438 | 0.0352 | 0.059 | 0.0112 | pred_leaf |

**Table S18.**

Ecomorph prediction posterior probabilities when using model incorporating full morphometric dataset; size-corrected ratio trait measurements; functional morphology.

| **Species** | **Cn** | **Gr** | **Lg** | **Ll** | **Sb** | **Predicted aspect** |
| --- | --- | --- | --- | --- | --- | --- |
| *Linguamyrmex vladi* | 0.0366 | 0.674 | 0.0064 | 0.2228 | 0.0602 | Gr |
| *Haidomyrmex scimitarus* | 0.0206 | 0.6094 | 0.0786 | 0.234 | 0.0574 | Gr |
| *Pseudomyrmex macrops* | 0.0138 | 0.0708 | 0.8422 | 0.0638 | 0.0094 | Lg |
| *Dhagnathos autokrator* | 0.0202 | 0.2926 | 0.1466 | 0.485 | 0.0556 | Ll |

**Table S19.**

Nesting niche prediction posterior probabilities when using model incorporating full morphometric dataset, size-corrected ratio trait measurements; homologous morphology.

| **Species** | **Ab** | **CR** | **Eg** | **Ll** | **Predicted aspect** |
| --- | --- | --- | --- | --- | --- |
| *Linguamyrmex vladi* | 0.1468 | 0.0678 | 0.6546 | 0.1308 | Eg |
| *Haidomyrmex scimitarus* | 0.1956 | 0.1006 | 0.509 | 0.1948 | Eg |
| *Pseudomyrmex macrops* | 0.8224 | 0.0124 | 0.1394 | 0.0258 | Ab |
| *Dhagnathos autokrator* | 0.1982 | 0.062 | 0.261 | 0.4788 | Ll |

**Table S20.**

Foraging niche prediction posterior probabilities when using model incorporating full morphometric dataset, size-corrected ratio trait measurements; homologous morphology.

| **Species** | **Fg** | **Gn** | **GP** | **Om** | **Py** | **SP** | **Predicted aspect** |
| --- | --- | --- | --- | --- | --- | --- | --- |
| *Linguamyrmex vladi* | 0.0276 | 0.0546 | 0.2514 | 0.3588 | 0.001 | 0.3066 | Om |
| *Haidomyrmex scimitarus* | 0.003 | 0.0196 | 0.2828 | 0.1952 | 0.037 | 0.4624 | SP |
| *Pseudomyrmex macrops* | 0 | 0.005 | 0.0872 | 0.0454 | 0.6976 | 0.1648 | Py |
| *Dhagnathos autokrator* | 0.0114 | 0.017 | 0.2314 | 0.1764 | 0.046 | 0.5178 | SP |

**Table S21.**

Functional role prediction posterior probabilities when using model incorporating full morphometric dataset, size-corrected ratio trait measurements; homologous morphology.

| **Species** | **om_cn_ab** | **om_grll_eg** | **om_leaf** | **om_lg_ab** | **pred_cr** | **pred_gr_egll** | **pred_leaf** | **pred_lg_ab** | **pred_ll_eg** | **pred_sub** | **Predicted aspect** |
| --- | --- | --- | --- | --- | --- | --- | --- | --- | --- | --- | --- |
| *Linguamyrmex vladi* | 0.05 | 0.303 | 0.0564 | 2.00E-04 | 0.0574 | 0.4042 | 0.1094 | 0.0082 | 0.011 | 2.00E-04 | pred_gr_egll |
| *Haidomyrmex scimitarus* | 0.0058 | 0.2034 | 0.0114 | 0.0124 | 0.0298 | 0.5468 | 0.091 | 0.0528 | 0.0294 | 0.0172 | pred_gr_egll |
| *Pseudomyrmex macrops* | 0.0062 | 0.04 | 0.003 | 0.6426 | 0.0074 | 0.0238 | 0.0102 | 0.2282 | 0.0372 | 0.0014 | om_lg_ab |
| *Dhagnathos autokrator* | 0.0156 | 0.1462 | 0.0214 | 0.0344 | 0.0212 | 0.1492 | 0.4522 | 0.1158 | 0.0172 | 0.0268 | pred_leaf |

**Table S22.**

Ecomorph prediction posterior probabilities when using model incorporating full morphometric dataset, size-corrected ratio trait measurements; homologous morphology.

| **Species** | **Cn** | **Gr** | **Lg** | **Ll** | **Sb** | **Predicted aspect** |
| --- | --- | --- | --- | --- | --- | --- |
| *Ceratomyrmex planus* | 0.0598 | 0.3838 | 0.0762 | 0.4338 | 0.0464 | Ll |
| *Ceratomyrmex ellenbergeri* | 0.057 | 0.485 | 0.0416 | 0.2394 | 0.177 | Gr |
| *Ceratomyrmex ellenbergeri* | 0.0382 | 0.4644 | 0.1014 | 0.3522 | 0.0438 | Gr |
| *Ceratomyrmex ellenbergeri* | 0.0736 | 0.519 | 0.0402 | 0.2826 | 0.0846 | Gr |
| *Protoceratomyrmex revelatus* | 0.0762 | 0.5304 | 0.0466 | 0.2896 | 0.0572 | Gr |
| *Linguamyrmex sp5* | 0.0912 | 0.6572 | 0.0166 | 0.202 | 0.033 | Gr |
| *Linguamyrmex sp4* | 0.0646 | 0.5374 | 0.0234 | 0.2556 | 0.119 | Gr |
| *Linguamyrmex brevicornis* | 0.0552 | 0.5154 | 0.0342 | 0.36 | 0.0352 | Gr |
| *Linguamyrmex sp2* | 0.0572 | 0.4372 | 0.0872 | 0.3666 | 0.0518 | Gr |
| *Linguamyrmex sp1* | 0.0662 | 0.4752 | 0.0368 | 0.2536 | 0.1682 | Gr |
| *Haidomyrmex sp3* | 0.0508 | 0.3036 | 0.2456 | 0.3598 | 0.0402 | Ll |
| *Haidomyrmex sp5* | 0.0426 | 0.355 | 0.1906 | 0.361 | 0.0508 | Ll |
| *Haidomyrmex sp5* | 0.0352 | 0.3664 | 0.2152 | 0.3422 | 0.041 | Gr |
| *Haidomyrmex sp5* | 0.0468 | 0.3816 | 0.1986 | 0.3056 | 0.0674 | Gr |
| *Haidomyrmex sp5* | 0.0376 | 0.3426 | 0.2926 | 0.2666 | 0.0606 | Cn |
| *Haidomyrmex scimitarus* | 0.0806 | 0.442 | 0.0288 | 0.249 | 0.1996 | Gr |
| *Haidomyrmex sp5* | 0.0412 | 0.4184 | 0.1834 | 0.2888 | 0.0682 | Gr |
| *Haidomyrmex sp4* | 0.0752 | 0.4584 | 0.0474 | 0.2482 | 0.1708 | Gr |
| *Haidomyrmex indet1* | 0.0294 | 0.3846 | 0.1996 | 0.3448 | 0.0416 | Gr |
| *Haidomyrmex indet2* | 0.0192 | 0.2474 | 0.2866 | 0.3914 | 0.0554 | Ll |

**Table S23.**

Nesting niche prediction posterior probabilities when using model incorporating subset of complete morphometrics dataset; raw trait measurements; functional morphology.

| **Species** | **Ab** | **CR** | **Eg** | **Ll** | **Predicted aspect** |
| --- | --- | --- | --- | --- | --- |
| *Ceratomyrmex planus* | 0.0986 | 0.0744 | 0.2564 | 0.5706 | Ll |
| *Ceratomyrmex ellenbergeri* | 0.1966 | 0.1784 | 0.384 | 0.241 | Eg |
| *Ceratomyrmex ellenbergeri* | 0.172 | 0.0578 | 0.2922 | 0.478 | Ll |
| *Ceratomyrmex ellenbergeri* | 0.13 | 0.0842 | 0.4284 | 0.3574 | Eg |
| *Protoceratomyrmex revelatus* | 0.1536 | 0.089 | 0.4234 | 0.334 | Eg |
| *Linguamyrmex sp5* | 0.252 | 0.019 | 0.653 | 0.076 | Eg |
| *Linguamyrmex sp4* | 0.238 | 0.0954 | 0.512 | 0.1546 | Eg |
| *Linguamyrmex brevicornis* | 0.1 | 0.0648 | 0.3994 | 0.4358 | Ll |
| *Linguamyrmex sp2* | 0.1608 | 0.0972 | 0.3546 | 0.3874 | Ll |
| *Linguamyrmex sp1* | 0.2018 | 0.1692 | 0.425 | 0.204 | Eg |
| *Haidomyrmex sp3* | 0.3014 | 0.0798 | 0.1928 | 0.426 | Ll |
| *Haidomyrmex sp5* | 0.2274 | 0.0466 | 0.2878 | 0.4382 | Ll |
| *Haidomyrmex sp5* | 0.2136 | 0.076 | 0.3052 | 0.4052 | Ll |
| *Haidomyrmex sp5* | 0.2506 | 0.0792 | 0.2868 | 0.3834 | Ll |
| *Haidomyrmex sp5* | 0.2774 | 0.0734 | 0.2738 | 0.3754 | Ll |
| *Haidomyrmex scimitarus* | 0.2156 | 0.1968 | 0.3992 | 0.1884 | Eg |
| *Haidomyrmex sp5* | 0.2306 | 0.1016 | 0.2492 | 0.4186 | Ll |
| *Haidomyrmex sp4* | 0.216 | 0.2006 | 0.4204 | 0.163 | Eg |
| *Haidomyrmex indet1* | 0.2432 | 0.1048 | 0.2476 | 0.4044 | Ll |
| *Haidomyrmex indet2* | 0.2582 | 0.0602 | 0.2704 | 0.4112 | Ll |

**Table S24.**

Foraging niche prediction posterior probabilities when using model incorporating subset of complete morphometrics dataset; raw trait measurements; functional morphology.

| **Species** | **Fg** | **Gn** | **GP** | **Om** | **Py** | **SP** | **Predicted aspect** |
| --- | --- | --- | --- | --- | --- | --- | --- |
| *Ceratomyrmex planus* | 0.1016 | 0.0506 | 0.1618 | 0.253 | 0.0566 | 0.3764 | SP |
| *Ceratomyrmex ellenbergeri* | 0.1676 | 0.0496 | 0.2944 | 0.2472 | 0.014 | 0.2272 | GP |
| *Ceratomyrmex ellenbergeri* | 0.1592 | 0.0742 | 0.3004 | 0.1192 | 0.0364 | 0.3106 | SP |
| *Ceratomyrmex ellenbergeri* | 0.2452 | 0.0546 | 0.2304 | 0.152 | 0.0104 | 0.3074 | SP |
| *Protoceratomyrmex revelatus* | 0.3096 | 0.08 | 0.0948 | 0.2104 | 0.0666 | 0.2386 | Fg |
| *Linguamyrmex sp5* | 0.1842 | 0.055 | 0.1616 | 0.3338 | 0.0028 | 0.2626 | Om |
| *Linguamyrmex sp4* | 0.1486 | 0.0734 | 0.2288 | 0.268 | 0.0068 | 0.2744 | SP |
| *Linguamyrmex brevicornis* | 0.229 | 0.1706 | 0.135 | 0.1818 | 0.0452 | 0.2384 | SP |
| *Linguamyrmex sp2* | 0.1612 | 0.0706 | 0.3864 | 0.161 | 0.0234 | 0.1974 | GP |
| *Linguamyrmex sp1* | 0.1314 | 0.0716 | 0.2904 | 0.2572 | 0.0114 | 0.238 | GP |
| *Haidomyrmex sp3* | 0.11 | 0.042 | 0.151 | 0.1024 | 0.1224 | 0.4722 | SP |
| *Haidomyrmex sp5* | 0.1528 | 0.047 | 0.2392 | 0.1324 | 0.0618 | 0.3668 | SP |
| *Haidomyrmex sp5* | 0.1302 | 0.0378 | 0.2434 | 0.1506 | 0.0624 | 0.3756 | SP |
| *Haidomyrmex sp5* | 0.1756 | 0.06 | 0.1452 | 0.1302 | 0.0696 | 0.4194 | SP |
| *Haidomyrmex sp5* | 0.2224 | 0.0808 | 0.1288 | 0.101 | 0.0666 | 0.4004 | SP |
| *Haidomyrmex scimitarus* | 0.108 | 0.0286 | 0.2866 | 0.3028 | 0.0176 | 0.2564 | Om |
| *Haidomyrmex sp5* | 0.1756 | 0.0372 | 0.187 | 0.1384 | 0.0416 | 0.4202 | SP |
| *Haidomyrmex sp4* | 0.1548 | 0.0384 | 0.275 | 0.278 | 0.0132 | 0.2406 | Om |
| *Haidomyrmex indet1* | 0.208 | 0.0376 | 0.2088 | 0.1938 | 0.0544 | 0.2974 | SP |
| *Haidomyrmex indet2* | 0.113 | 0.075 | 0.1456 | 0.0944 | 0.0948 | 0.4772 | SP |

**Table S25.**

Functional role prediction posterior probabilities when using model incorporating subset of complete morphometrics dataset; raw trait measurements; functional morphology.

| **Species** | **om_cn_ab** | **om_grll_eg** | **om_leaf** | **om_lg_ab** | **pred_cr** | **pred_gr_egll** | **pred_leaf** | **pred_lg_ab** | **pred_ll_eg** | **pred_sub** | **Predicted aspect** |
| --- | --- | --- | --- | --- | --- | --- | --- | --- | --- | --- | --- |
| *Ceratomyrmex planus* | 0.0578 | 0.2526 | 0.158 | 0.0544 | 0.0754 | 0.064 | 0.2268 | 0.0848 | 0.0038 | 0.0224 | om_grll_eg |
| *Ceratomyrmex ellenbergeri* | 0.0656 | 0.3268 | 0.0108 | 0.0118 | 0.0564 | 0.1744 | 0.2716 | 0.017 | 0.0654 | 0 | om_grll_eg |
| *Ceratomyrmex ellenbergeri* | 0.0436 | 0.2244 | 0.0762 | 0.0712 | 0.0444 | 0.0384 | 0.4158 | 0.0436 | 0.0324 | 0.01 | pred_leaf |
| *Ceratomyrmex ellenbergeri* | 0.0506 | 0.2992 | 0.0264 | 0.026 | 0.0462 | 0.0866 | 0.3874 | 0.012 | 0.0626 | 0.003 | pred_leaf |
| *Protoceratomyrmex revelatus* | 0.1194 | 0.405 | 0.108 | 0.0892 | 0.0718 | 0.028 | 0.111 | 0.0174 | 0.0446 | 0.0056 | om_grll_eg |
| *Linguamyrmex sp5* | 0.0838 | 0.396 | 0.0028 | 0.0014 | 0.012 | 0.2324 | 0.1946 | 0.017 | 0.0598 | 0 | om_grll_eg |
| *Linguamyrmex sp4* | 0.0814 | 0.3842 | 0.0034 | 0.0026 | 0.0236 | 0.1794 | 0.2432 | 0.0106 | 0.0714 | 0 | om_grll_eg |
| *Linguamyrmex brevicornis* | 0.0716 | 0.4046 | 0.2012 | 0.065 | 0.0378 | 0.0432 | 0.138 | 0.0076 | 0.0254 | 0.0056 | om_grll_eg |
| *Linguamyrmex sp2* | 0.0468 | 0.2786 | 0.0312 | 0.0276 | 0.064 | 0.0502 | 0.4588 | 0.0112 | 0.0298 | 0.0018 | pred_leaf |
| *Linguamyrmex sp1* | 0.079 | 0.3368 | 0.008 | 0.0062 | 0.043 | 0.2066 | 0.234 | 0.0132 | 0.073 | 0 | om_grll_eg |
| *Haidomyrmex sp3* | 0.1102 | 0.1584 | 0.0758 | 0.1518 | 0.0394 | 0.0286 | 0.2254 | 0.189 | 0.0086 | 0.0128 | pred_leaf |
| *Haidomyrmex sp5* | 0.0472 | 0.1746 | 0.0562 | 0.1264 | 0.0416 | 0.0444 | 0.3034 | 0.1356 | 0.0572 | 0.0134 | pred_leaf |
| *Haidomyrmex sp5* | 0.0438 | 0.207 | 0.044 | 0.1276 | 0.071 | 0.0358 | 0.2872 | 0.1122 | 0.0628 | 0.0086 | pred_leaf |
| *Haidomyrmex sp5* | 0.06 | 0.1958 | 0.0538 | 0.1508 | 0.0646 | 0.0372 | 0.2004 | 0.147 | 0.0746 | 0.0158 | pred_leaf |
| *Haidomyrmex sp5* | 0.0418 | 0.2396 | 0.0426 | 0.1212 | 0.0542 | 0.0428 | 0.209 | 0.1694 | 0.0548 | 0.0246 | om_grll_eg |
| *Haidomyrmex scimitarus* | 0.082 | 0.3636 | 0.0036 | 0.0108 | 0.0596 | 0.2104 | 0.1606 | 0.0096 | 0.0998 | 0 | om_grll_eg |
| *Haidomyrmex sp5* | 0.0688 | 0.2302 | 0.091 | 0.0952 | 0.0628 | 0.0496 | 0.176 | 0.1854 | 0.0224 | 0.0186 | om_grll_eg |
| *Haidomyrmex sp4* | 0.0898 | 0.3548 | 0.0016 | 0.0156 | 0.0662 | 0.2046 | 0.1214 | 0.011 | 0.1344 | 0 | om_grll_eg |
| *Haidomyrmex indet1* | 0.0544 | 0.2744 | 0.106 | 0.109 | 0.053 | 0.0436 | 0.1388 | 0.1768 | 0.0294 | 0.0146 | om_grll_eg |
| *Haidomyrmex indet2* | 0.0358 | 0.2108 | 0.0304 | 0.162 | 0.0486 | 0.0794 | 0.1852 | 0.1034 | 0.125 | 0.0194 | om_grll_eg |

**Table S26.**

Ecomorph prediction posterior probabilities when using model incorporating subset of complete morphometrics dataset; raw trait measurements; functional morphology.

| **Species** | **Cn** | **Gr** | **Lg** | **Ll** | **Sb** | **Predicted aspect** |
| --- | --- | --- | --- | --- | --- | --- |
| *Ceratomyrmex planus* | 0.0384 | 0.2442 | 0.0898 | 0.5854 | 0.0422 | Ll |
| *Ceratomyrmex ellenbergeri* | 0.0074 | 0.4942 | 0.0608 | 0.2844 | 0.1532 | Gr |
| *Ceratomyrmex ellenbergeri* | 0.0574 | 0.1966 | 0.1402 | 0.507 | 0.0988 | Ll |
| *Ceratomyrmex ellenbergeri* | 0.034 | 0.2198 | 0.0466 | 0.5722 | 0.1274 | Ll |
| *Protoceratomyrmex revelatus* | 0.033 | 0.2098 | 0.0432 | 0.6314 | 0.0826 | Ll |
| *Linguamyrmex sp5* | 0.0084 | 0.8596 | 0.0304 | 0.0828 | 0.0188 | Gr |
| *Linguamyrmex sp4* | 0.0058 | 0.6526 | 0.046 | 0.1858 | 0.1098 | Gr |
| *Linguamyrmex brevicornis* | 0.0482 | 0.1406 | 0.0376 | 0.708 | 0.0656 | Ll |
| *Linguamyrmex sp2* | 0.019 | 0.184 | 0.1028 | 0.6352 | 0.059 | Ll |
| *Linguamyrmex sp1* | 0.0088 | 0.577 | 0.0556 | 0.2204 | 0.1382 | Gr |
| *Haidomyrmex sp3* | 0.0662 | 0.2 | 0.2552 | 0.4328 | 0.0458 | Ll |
| *Haidomyrmex sp5* | 0.043 | 0.3078 | 0.2038 | 0.401 | 0.0444 | Ll |
| *Haidomyrmex sp5* | 0.0374 | 0.1636 | 0.2776 | 0.4484 | 0.073 | Ll |
| *Haidomyrmex sp5* | 0.0424 | 0.1274 | 0.234 | 0.5152 | 0.081 | Ll |
| *Haidomyrmex sp5* | 0.0202 | 0.1118 | 0.2906 | 0.4878 | 0.0896 | Ll |
| *Haidomyrmex scimitarus* | 0.0134 | 0.5824 | 0.0606 | 0.214 | 0.1296 | Gr |
| *Haidomyrmex sp5* | 0.062 | 0.1338 | 0.223 | 0.4516 | 0.1296 | Ll |
| *Haidomyrmex sp4* | 0.0292 | 0.5052 | 0.079 | 0.2054 | 0.1812 | Gr |
| *Haidomyrmex indet1* | 0.0266 | 0.3602 | 0.2272 | 0.3438 | 0.0422 | Gr |
| *Haidomyrmex indet2* | 0.0136 | 0.2024 | 0.2682 | 0.446 | 0.0698 | Ll |

**Table S27.**

Nesting niche prediction posterior probabilities when using model incorporating subset of complete morphometrics dataset; raw trait measurements; homologous morphology.

| **Species** | **Ab** | **CR** | **Eg** | **Ll** | **Predicted aspect** |
| --- | --- | --- | --- | --- | --- |
| *Ceratomyrmex planus* | 0.0864 | 0.0914 | 0.1778 | 0.6444 | Ll |
| *Ceratomyrmex ellenbergeri* | 0.0928 | 0.1676 | 0.4108 | 0.3288 | Eg |
| *Ceratomyrmex ellenbergeri* | 0.1808 | 0.075 | 0.2518 | 0.4924 | Ll |
| *Ceratomyrmex ellenbergeri* | 0.0866 | 0.1364 | 0.3178 | 0.4592 | Ll |
| *Protoceratomyrmex revelatus* | 0.0776 | 0.102 | 0.3904 | 0.43 | Ll |
| *Linguamyrmex sp5* | 0.1526 | 0.0194 | 0.7488 | 0.0792 | Eg |
| *Linguamyrmex sp4* | 0.1634 | 0.1106 | 0.5472 | 0.1788 | Eg |
| *Linguamyrmex brevicornis* | 0.0778 | 0.0826 | 0.2944 | 0.5452 | Ll |
| *Linguamyrmex sp2* | 0.065 | 0.1212 | 0.2836 | 0.5302 | Ll |
| *Linguamyrmex sp1* | 0.1474 | 0.1606 | 0.4448 | 0.2472 | Eg |
| *Haidomyrmex sp3* | 0.3078 | 0.0824 | 0.1786 | 0.4312 | Ll |
| *Haidomyrmex sp5* | 0.217 | 0.0488 | 0.3048 | 0.4294 | Ll |
| *Haidomyrmex sp5* | 0.2186 | 0.0836 | 0.2414 | 0.4564 | Ll |
| *Haidomyrmex sp5* | 0.232 | 0.059 | 0.263 | 0.446 | Ll |
| *Haidomyrmex sp5* | 0.232 | 0.1114 | 0.2186 | 0.438 | Ll |
| *Haidomyrmex scimitarus* | 0.2052 | 0.13 | 0.414 | 0.2508 | Eg |
| *Haidomyrmex sp5* | 0.2532 | 0.0988 | 0.171 | 0.477 | Ll |
| *Haidomyrmex sp4* | 0.1976 | 0.1906 | 0.4222 | 0.1896 | Eg |
| *Haidomyrmex indet1* | 0.2514 | 0.085 | 0.2622 | 0.4014 | Ll |
| *Haidomyrmex indet2* | 0.208 | 0.0832 | 0.258 | 0.4508 | Ll |

**Table S28.**

Foraging niche prediction posterior probabilities when using model incorporating subset of complete morphometrics dataset; raw trait measurements; homologous morphology.

| **Species** | **Fg** | **Gn** | **GP** | **Om** | **Py** | **SP** | **Predicted aspect** |
| --- | --- | --- | --- | --- | --- | --- | --- |
| *Ceratomyrmex planus* | 0.0998 | 0.0466 | 0.1796 | 0.1574 | 0.0518 | 0.4648 | SP |
| *Ceratomyrmex ellenbergeri* | 0.0318 | 0.061 | 0.3106 | 0.2362 | 0.08 | 0.2804 | GP |
| *Ceratomyrmex ellenbergeri* | 0.0848 | 0.1064 | 0.322 | 0.1258 | 0.0582 | 0.3028 | GP |
| *Ceratomyrmex ellenbergeri* | 0.046 | 0.0978 | 0.3204 | 0.2192 | 0.037 | 0.2796 | GP |
| *Protoceratomyrmex revelatus* | 0.0794 | 0.0954 | 0.216 | 0.289 | 0.067 | 0.2532 | Om |
| *Linguamyrmex sp5* | 0.0396 | 0.0306 | 0.1186 | 0.4328 | 0.0542 | 0.3242 | Om |
| *Linguamyrmex sp4* | 0.0216 | 0.0636 | 0.1878 | 0.3358 | 0.0732 | 0.318 | Om |
| *Linguamyrmex brevicornis* | 0.0844 | 0.1938 | 0.194 | 0.2334 | 0.0452 | 0.2492 | SP |
| *Linguamyrmex sp2* | 0.0386 | 0.1178 | 0.3982 | 0.1722 | 0.0582 | 0.215 | GP |
| *Linguamyrmex sp1* | 0.0234 | 0.055 | 0.2526 | 0.3058 | 0.081 | 0.2822 | Om |
| *Haidomyrmex sp3* | 0.057 | 0.0388 | 0.1812 | 0.1338 | 0.1736 | 0.4156 | SP |
| *Haidomyrmex sp5* | 0.1042 | 0.0644 | 0.2236 | 0.1662 | 0.0912 | 0.3504 | SP |
| *Haidomyrmex sp5* | 0.0878 | 0.0904 | 0.2576 | 0.1102 | 0.1004 | 0.3536 | SP |
| *Haidomyrmex sp5* | 0.0544 | 0.0852 | 0.195 | 0.1636 | 0.0974 | 0.4044 | SP |
| *Haidomyrmex sp5* | 0.0322 | 0.0854 | 0.2618 | 0.1664 | 0.085 | 0.3692 | SP |
| *Haidomyrmex scimitarus* | 0.0564 | 0.0386 | 0.1404 | 0.3496 | 0.0724 | 0.3426 | SP |
| *Haidomyrmex sp5* | 0.0554 | 0.0408 | 0.2708 | 0.1428 | 0.0882 | 0.402 | SP |
| *Haidomyrmex sp4* | 0.044 | 0.0378 | 0.1976 | 0.3738 | 0.0844 | 0.2624 | Om |
| *Haidomyrmex indet1* | 0.1206 | 0.0274 | 0.2576 | 0.1944 | 0.1068 | 0.2932 | SP |
| *Haidomyrmex indet2* | 0.0356 | 0.113 | 0.2284 | 0.1366 | 0.0994 | 0.387 | SP |

**Table S29.**

Functional role prediction posterior probabilities when using model incorporating subset of complete morphometrics dataset; raw trait measurements; homologous morphology.

| **Species** | **om_cn_ab** | **om_grll_eg** | **om_leaf** | **om_lg_ab** | **pred_cr** | **pred_gr_egll** | **pred_leaf** | **pred_lg_ab** | **pred_ll_eg** | **pred_sub** | **Predicted aspect** |
| --- | --- | --- | --- | --- | --- | --- | --- | --- | --- | --- | --- |
| *Ceratomyrmex planus* | 0.0424 | 0.135 | 0.219 | 0.0462 | 0.0718 | 0.1054 | 0.2652 | 0.093 | 0.0024 | 0.0196 | pred_leaf |
| *Ceratomyrmex ellenbergeri* | 0.0082 | 0.2986 | 0.0144 | 0.0644 | 0.0508 | 0.1856 | 0.3356 | 0.0094 | 0.0322 | 0 | pred_leaf |
| *Ceratomyrmex ellenbergeri* | 0.0492 | 0.1868 | 0.0918 | 0.0972 | 0.0586 | 0.0274 | 0.4056 | 0.0386 | 0.0272 | 0.0176 | pred_leaf |
| *Ceratomyrmex ellenbergeri* | 0.0366 | 0.2668 | 0.0392 | 0.052 | 0.0848 | 0.0512 | 0.409 | 0.0092 | 0.0462 | 0.005 | pred_leaf |
| *Protoceratomyrmex revelatus* | 0.0812 | 0.4286 | 0.0738 | 0.0708 | 0.105 | 0.0378 | 0.1346 | 0.0224 | 0.0384 | 0.0074 | om_grll_eg |
| *Linguamyrmex sp5* | 0.0114 | 0.4306 | 0.0036 | 0.0464 | 0.016 | 0.2274 | 0.224 | 0.0142 | 0.026 | 0 | om_grll_eg |
| *Linguamyrmex sp4* | 0.0122 | 0.3858 | 0.0062 | 0.051 | 0.0414 | 0.1664 | 0.2982 | 0.0082 | 0.0302 | 0 | om_grll_eg |
| *Linguamyrmex brevicornis* | 0.0688 | 0.373 | 0.1954 | 0.0668 | 0.0672 | 0.0338 | 0.1634 | 0.0052 | 0.0162 | 0.0102 | om_grll_eg |
| *Linguamyrmex sp2* | 0.019 | 0.1966 | 0.0568 | 0.065 | 0.089 | 0.0274 | 0.517 | 0.0056 | 0.0182 | 0.0054 | pred_leaf |
| *Linguamyrmex sp1* | 0.0102 | 0.3314 | 0.0094 | 0.0618 | 0.0484 | 0.199 | 0.295 | 0.011 | 0.0332 | 0 | om_grll_eg |
| *Haidomyrmex sp3* | 0.1096 | 0.167 | 0.0658 | 0.1542 | 0.0406 | 0.0172 | 0.2366 | 0.1888 | 0.0072 | 0.013 | pred_leaf |
| *Haidomyrmex sp5* | 0.0464 | 0.212 | 0.0738 | 0.1242 | 0.0378 | 0.0362 | 0.2596 | 0.1398 | 0.0498 | 0.0204 | pred_leaf |
| *Haidomyrmex sp5* | 0.0406 | 0.163 | 0.0836 | 0.1436 | 0.0832 | 0.0212 | 0.2748 | 0.1196 | 0.0544 | 0.016 | pred_leaf |
| *Haidomyrmex sp5* | 0.057 | 0.2198 | 0.0486 | 0.1436 | 0.0682 | 0.0162 | 0.2008 | 0.1536 | 0.067 | 0.0252 | om_grll_eg |
| *Haidomyrmex sp5* | 0.0332 | 0.19 | 0.044 | 0.1394 | 0.1132 | 0.014 | 0.2506 | 0.1464 | 0.0426 | 0.0266 | pred_leaf |
| *Haidomyrmex scimitarus* | 0.0154 | 0.3938 | 0.008 | 0.0652 | 0.0406 | 0.2328 | 0.1854 | 0.009 | 0.0492 | 0 | om_grll_eg |
| *Haidomyrmex sp5* | 0.0706 | 0.1688 | 0.0652 | 0.1158 | 0.0928 | 0.0496 | 0.198 | 0.1944 | 0.0238 | 0.021 | pred_leaf |
| *Haidomyrmex sp4* | 0.0254 | 0.4188 | 0.0042 | 0.0784 | 0.0728 | 0.1626 | 0.163 | 0.011 | 0.0628 | 0.001 | om_grll_eg |
| *Haidomyrmex indet1* | 0.043 | 0.2726 | 0.0744 | 0.1262 | 0.0422 | 0.0516 | 0.1436 | 0.213 | 0.022 | 0.0114 | om_grll_eg |
| *Haidomyrmex indet2* | 0.0324 | 0.1804 | 0.0488 | 0.1762 | 0.1038 | 0.0574 | 0.1936 | 0.0774 | 0.1062 | 0.0238 | pred_leaf |

**Table S30.**

Ecomorph prediction posterior probabilities when using model incorporating subset of complete morphometrics dataset; raw trait measurements; homologous morphology.

| **Species** | **Cn** | **Gr** | **Lg** | **Ll** | **Sb** | **Predicted aspect** |
| --- | --- | --- | --- | --- | --- | --- |
| *Ceratomyrmex planus* | 0.0684 | 0.6814 | 0.0074 | 0.2158 | 0.027 | Gr |
| *Ceratomyrmex ellenbergeri* | 0.0424 | 0.7562 | 6.00E-04 | 0.1998 | 0.001 | Gr |
| *Ceratomyrmex ellenbergeri* | 0.0482 | 0.704 | 0.0018 | 0.2322 | 0.0138 | Gr |
| *Ceratomyrmex ellenbergeri* | 0.0326 | 0.7176 | 0.023 | 0.2146 | 0.0122 | Gr |
| *Protoceratomyrmex revelatus* | 0.0256 | 0.6942 | 0.0226 | 0.242 | 0.0156 | Gr |
| *Linguamyrmex sp5* | 0.0252 | 0.6544 | 0.002 | 0.3162 | 0.0022 | Gr |
| *Linguamyrmex sp4* | 0.0212 | 0.6548 | 0.0044 | 0.3188 | 0 | Gr |
| *Linguamyrmex brevicornis* | 0.2636 | 0.2216 | 0.0608 | 0.4474 | 0.0066 | Ll |
| *Linguamyrmex sp2* | 0.0634 | 0.6044 | 0.0108 | 0.3094 | 0.012 | Gr |
| *Linguamyrmex sp1* | 0.0434 | 0.7008 | 0.0024 | 0.239 | 0.0144 | Gr |
| *Haidomyrmex sp3* | 0.0206 | 0.1626 | 0.5006 | 0.2918 | 0.0244 | Lg |
| *Haidomyrmex sp5* | 0.004 | 0.548 | 0.0458 | 0.4 | 0.0022 | Gr |
| *Haidomyrmex sp5* | 0.0116 | 0.8092 | 0.0194 | 0.1556 | 0.0042 | Gr |
| *Haidomyrmex sp5* | 0.0022 | 0.743 | 0.0856 | 0.1658 | 0.0034 | Gr |
| *Haidomyrmex sp5* | 0.0304 | 0.3702 | 0.258 | 0.32 | 0.0214 | Gr |
| *Haidomyrmex scimitarus* | 0.055 | 0.7134 | 0.0088 | 0.2052 | 0.0176 | Gr |
| *Haidomyrmex sp5* | 0.0026 | 0.5238 | 0.2264 | 0.2288 | 0.0184 | Gr |
| *Haidomyrmex sp4* | 0.0066 | 0.707 | 0.0744 | 0.2032 | 0.0088 | Gr |
| *Haidomyrmex indet1* | 0.0022 | 0.7738 | 0.0692 | 0.1506 | 0.0042 | Gr |
| *Haidomyrmex indet2* | 0.003 | 0.5718 | 0.1326 | 0.2886 | 0.004 | Gr |

**Table S31.**

Nesting niche prediction posterior probabilities when using model incorporating subset of complete morphometrics dataset; size-corrected ratio trait measurements; functional morphology.

| **Species** | **Ab** | **CR** | **Eg** | **Ll** | **Predicted aspect** |
| --- | --- | --- | --- | --- | --- |
| *Ceratomyrmex planus* | 0.1752 | 0.0844 | 0.5354 | 0.205 | Eg |
| *Ceratomyrmex ellenbergeri* | 0.1584 | 0.0032 | 0.59 | 0.2484 | Eg |
| *Ceratomyrmex ellenbergeri* | 0.1658 | 0.0132 | 0.5646 | 0.2564 | Eg |
| *Ceratomyrmex ellenbergeri* | 0.11 | 0.0826 | 0.5422 | 0.2652 | Eg |
| *Protoceratomyrmex revelatus* | 0.2024 | 0.0676 | 0.6052 | 0.1248 | Eg |
| *Linguamyrmex sp5* | 0.1838 | 0.0022 | 0.4504 | 0.3636 | Eg |
| *Linguamyrmex sp4* | 0.1978 | 0.002 | 0.5542 | 0.246 | Eg |
| *Linguamyrmex brevicornis* | 0.403 | 0.0424 | 0.2646 | 0.29 | Ab |
| *Linguamyrmex sp2* | 0.2446 | 0.0182 | 0.502 | 0.2352 | Eg |
| *Linguamyrmex sp1* | 0.2264 | 0.0142 | 0.4928 | 0.2666 | Eg |
| *Haidomyrmex sp3* | 0.3572 | 0.2008 | 0.2432 | 0.1988 | Ab |
| *Haidomyrmex sp5* | 0.1086 | 0.0674 | 0.5246 | 0.2994 | Eg |
| *Haidomyrmex sp5* | 0.1166 | 0.0642 | 0.6328 | 0.1864 | Eg |
| *Haidomyrmex sp5* | 0.1594 | 0.0844 | 0.5338 | 0.2224 | Eg |
| *Haidomyrmex sp5* | 0.2292 | 0.162 | 0.4284 | 0.1804 | Eg |
| *Haidomyrmex scimitarus* | 0.165 | 0.0598 | 0.5762 | 0.199 | Eg |
| *Haidomyrmex sp5* | 0.1918 | 0.0992 | 0.4758 | 0.2332 | Eg |
| *Haidomyrmex sp4* | 0.1104 | 0.0872 | 0.584 | 0.2184 | Eg |
| *Haidomyrmex indet1* | 0.0934 | 0.057 | 0.6406 | 0.209 | Eg |
| *Haidomyrmex indet2* | 0.1078 | 0.0872 | 0.5662 | 0.2388 | Eg |

**Table S32.**

Foraging niche prediction posterior probabilities when using model incorporating subset of complete morphometrics dataset; size-corrected ratio trait measurements; functional morphology.

| **Species** | **Fg** | **Gn** | **GP** | **Om** | **Py** | **SP** | **Predicted aspect** |
| --- | --- | --- | --- | --- | --- | --- | --- |
| *Ceratomyrmex planus* | 0.0434 | 0.1026 | 0.1652 | 0.4172 | 2.00E-04 | 0.2714 | Om |
| *Ceratomyrmex ellenbergeri* | 0.1128 | 0.0178 | 0.5196 | 0.2742 | 0 | 0.075 | GP |
| *Ceratomyrmex ellenbergeri* | 0.0924 | 0.0332 | 0.4498 | 0.283 | 0.001 | 0.1406 | GP |
| *Ceratomyrmex ellenbergeri* | 0.066 | 0.0764 | 0.4536 | 0.2248 | 0.0022 | 0.177 | GP |
| *Protoceratomyrmex revelatus* | 0.0106 | 0.0128 | 0.1504 | 0.2706 | 0.001 | 0.5546 | SP |
| *Linguamyrmex sp5* | 0.0942 | 0.0364 | 0.3288 | 0.4298 | 0 | 0.1106 | Om |
| *Linguamyrmex sp4* | 0.0596 | 0.0104 | 0.2864 | 0.4004 | 0 | 0.243 | Om |
| *Linguamyrmex brevicornis* | 0.0314 | 0.024 | 0.1766 | 0.3368 | 0.0014 | 0.4298 | SP |
| *Linguamyrmex sp2* | 0.1124 | 0.0122 | 0.336 | 0.34 | 0 | 0.1994 | Om |
| *Linguamyrmex sp1* | 0.0606 | 0.0218 | 0.4288 | 0.331 | 0 | 0.1578 | GP |
| *Haidomyrmex sp3* | 0.009 | 8.00E-04 | 0.3848 | 0.101 | 0.0618 | 0.4426 | SP |
| *Haidomyrmex sp5* | 0.0032 | 0.0014 | 0.6024 | 0.1608 | 0.0156 | 0.2166 | GP |
| *Haidomyrmex sp5* | 0.0036 | 2.00E-04 | 0.7118 | 0.1194 | 0.0092 | 0.1558 | GP |
| *Haidomyrmex sp5* | 0.0038 | 0.0296 | 0.5182 | 0.1168 | 0.0202 | 0.3114 | GP |
| *Haidomyrmex sp5* | 0.0934 | 0.0184 | 0.4184 | 0.1162 | 0.0288 | 0.3248 | GP |
| *Haidomyrmex scimitarus* | 0.0352 | 0.0128 | 0.428 | 0.2884 | 0.0012 | 0.2344 | GP |
| *Haidomyrmex sp5* | 0.0064 | 0.0106 | 0.3556 | 0.094 | 0.0446 | 0.4888 | SP |
| *Haidomyrmex sp4* | 0.003 | 0.035 | 0.522 | 0.1148 | 0.0182 | 0.307 | GP |
| *Haidomyrmex indet1* | 0.002 | 0.0092 | 0.6058 | 0.0974 | 0.0182 | 0.2674 | GP |
| *Haidomyrmex indet2* | 0.0038 | 0.0374 | 0.2948 | 0.1204 | 0.0324 | 0.5112 | SP |

**Table S33.**

Functional role prediction posterior probabilities when using model incorporating subset of complete morphometrics dataset; size-corrected ratio trait measurements; functional morphology.

| **Species** | **om_cn_ab** | **om_grll_eg** | **om_leaf** | **om_lg_ab** | **pred_cr** | **pred_gr_egll** | **pred_leaf** | **pred_lg_ab** | **pred_ll_eg** | **pred_sub** | **Predicted aspect** |
| --- | --- | --- | --- | --- | --- | --- | --- | --- | --- | --- | --- |
| *Ceratomyrmex planus* | 0.055 | 0.328 | 0.1102 | 0 | 0.0292 | 0.3218 | 0.1372 | 0.0062 | 0.0116 | 0 | om_grll_eg |
| *Ceratomyrmex ellenbergeri* | 0.063 | 0.2064 | 0.0466 | 0 | 0.0028 | 0.5138 | 0.1148 | 0 | 0.0522 | 0 | pred_gr_egll |
| *Ceratomyrmex ellenbergeri* | 0.059 | 0.216 | 0.0606 | 0 | 0.0142 | 0.3868 | 0.2192 | 0.001 | 0.0428 | 0 | pred_gr_egll |
| *Ceratomyrmex ellenbergeri* | 0.02 | 0.2284 | 0.0272 | 0.0028 | 0.02 | 0.549 | 0.1062 | 0.009 | 0.0364 | 0.001 | pred_gr_egll |
| *Protoceratomyrmex revelatus* | 0.0142 | 0.2096 | 0.004 | 0.0026 | 0.0146 | 0.5278 | 0.0854 | 0.0326 | 0.1084 | 0 | pred_gr_egll |
| *Linguamyrmex sp5* | 0.0452 | 0.3486 | 0.061 | 0 | 0.0012 | 0.238 | 0.272 | 0.0036 | 0.0304 | 0 | om_grll_eg |
| *Linguamyrmex sp4* | 0.0426 | 0.2928 | 0.0306 | 0 | 0.001 | 0.32 | 0.2546 | 0.0018 | 0.0566 | 0 | pred_gr_egll |
| *Linguamyrmex brevicornis* | 0.1488 | 0.1564 | 0.0172 | 0.0018 | 0.0196 | 0.2324 | 0.3284 | 0.0674 | 0.0274 | 0 | pred_leaf |
| *Linguamyrmex sp2* | 0.0688 | 0.2716 | 0.032 | 0 | 0.0136 | 0.3018 | 0.2594 | 0.0064 | 0.0456 | 0 | pred_gr_egll |
| *Linguamyrmex sp1* | 0.0508 | 0.2064 | 0.0326 | 0 | 0.011 | 0.3674 | 0.283 | 0.001 | 0.0472 | 0 | pred_gr_egll |
| *Haidomyrmex sp3* | 0.0154 | 0.0296 | 0.001 | 0.088 | 0.024 | 0.147 | 0.1572 | 0.4262 | 0.1048 | 0.0068 | pred_lg_ab |
| *Haidomyrmex sp5* | 0.0038 | 0.1488 | 0.0034 | 0.0074 | 0.0046 | 0.5314 | 0.1946 | 0.0162 | 0.0892 | 0 | pred_gr_egll |
| *Haidomyrmex sp5* | 0.0078 | 0.0982 | 0.0012 | 0.003 | 0.006 | 0.6876 | 0.0702 | 0.0122 | 0.1134 | 0 | pred_gr_egll |
| *Haidomyrmex sp5* | 0.0016 | 0.1446 | 0.002 | 0.0246 | 0.0076 | 0.5886 | 0.1084 | 0.044 | 0.0768 | 0.0018 | pred_gr_egll |
| *Haidomyrmex sp5* | 0.0298 | 0.111 | 0.0222 | 0.0962 | 0.0434 | 0.2588 | 0.1262 | 0.1654 | 0.141 | 0.006 | pred_gr_egll |
| *Haidomyrmex scimitarus* | 0.0334 | 0.1938 | 0.0172 | 0.0014 | 0.02 | 0.4756 | 0.208 | 0.0042 | 0.046 | 0 | pred_gr_egll |
| *Haidomyrmex sp5* | 0.002 | 0.0712 | 0 | 0.1434 | 0.0086 | 0.3544 | 0.1126 | 0.1754 | 0.1244 | 0.008 | pred_gr_egll |
| *Haidomyrmex sp4* | 0.0062 | 0.1108 | 0.001 | 0.0096 | 0.022 | 0.6138 | 0.0988 | 0.0278 | 0.109 | 0.001 | pred_gr_egll |
| *Haidomyrmex indet1* | 0.0014 | 0.084 | 0 | 0.0096 | 0.0118 | 0.7518 | 0.084 | 0.0246 | 0.0304 | 0.0022 | pred_gr_egll |
| *Haidomyrmex indet2* | 0.002 | 0.095 | 0.0036 | 0.0882 | 0.0156 | 0.431 | 0.1304 | 0.1152 | 0.1168 | 0.0022 | pred_gr_egll |

**Table S34.**

Ecomorph prediction posterior probabilities when using model incorporating subset of complete morphometrics dataset; size-corrected ratio trait measurements; functional morphology.

| **Species** | **Cn** | **Gr** | **Lg** | **Ll** | **Sb** | **Predicted aspect** |
| --- | --- | --- | --- | --- | --- | --- |
| *Ceratomyrmex planus* | 0.0392 | 0.5364 | 0.0298 | 0.3762 | 0.0184 | Gr |
| *Ceratomyrmex ellenbergeri* | 0.0236 | 0.8092 | 0.0024 | 0.1628 | 0.002 | Gr |
| *Ceratomyrmex ellenbergeri* | 0.024 | 0.7864 | 0.0026 | 0.176 | 0.011 | Gr |
| *Ceratomyrmex ellenbergeri* | 0.0238 | 0.6812 | 0.0424 | 0.2208 | 0.0318 | Gr |
| *Protoceratomyrmex revelatus* | 0.0202 | 0.7678 | 0.0176 | 0.1806 | 0.0138 | Gr |
| *Linguamyrmex sp5* | 0.0102 | 0.7622 | 0.002 | 0.2246 | 0.001 | Gr |
| *Linguamyrmex sp4* | 0.0062 | 0.7626 | 0.0038 | 0.2252 | 0.0022 | Gr |
| *Linguamyrmex brevicornis* | 0.2566 | 0.2498 | 0.0588 | 0.4286 | 0.0062 | Ll |
| *Linguamyrmex sp2* | 0.048 | 0.6416 | 0.0176 | 0.2768 | 0.016 | Gr |
| *Linguamyrmex sp1* | 0.0272 | 0.7662 | 0.0028 | 0.187 | 0.0168 | Gr |
| *Haidomyrmex sp3* | 0.013 | 0.1366 | 0.5272 | 0.2864 | 0.0368 | Lg |
| *Haidomyrmex sp5* | 0.0052 | 0.4744 | 0.0716 | 0.4432 | 0.0056 | Gr |
| *Haidomyrmex sp5* | 0.0084 | 0.7316 | 0.0324 | 0.2228 | 0.0048 | Gr |
| *Haidomyrmex sp5* | 0.004 | 0.655 | 0.1158 | 0.202 | 0.0232 | Gr |
| *Haidomyrmex sp5* | 0.0328 | 0.2728 | 0.3176 | 0.341 | 0.0358 | Ll |
| *Haidomyrmex scimitarus* | 0.0308 | 0.7934 | 0.01 | 0.1502 | 0.0156 | Gr |
| *Haidomyrmex sp5* | 0.0024 | 0.4766 | 0.2318 | 0.2524 | 0.0368 | Gr |
| *Haidomyrmex sp4* | 0.008 | 0.6384 | 0.094 | 0.2388 | 0.0208 | Gr |
| *Haidomyrmex indet1* | 0.0034 | 0.6742 | 0.0928 | 0.2118 | 0.0178 | Gr |
| *Haidomyrmex indet2* | 0.0022 | 0.5782 | 0.149 | 0.2532 | 0.0174 | Gr |

**Table S35.**

Nesting niche prediction posterior probabilities when using model incorporating subset of complete morphometrics dataset; size-corrected ratio trait measurements; homologous morphology.

| **Species** | **Ab** | **CR** | **Eg** | **Ll** | **Predicted aspect** |
| --- | --- | --- | --- | --- | --- |
| *Ceratomyrmex planus* | 0.099 | 0.0394 | 0.5154 | 0.3462 | Eg |
| *Ceratomyrmex ellenbergeri* | 0.151 | 0.004 | 0.6054 | 0.2396 | Eg |
| *Ceratomyrmex ellenbergeri* | 0.0936 | 0.01 | 0.6584 | 0.238 | Eg |
| *Ceratomyrmex ellenbergeri* | 0.1374 | 0.0996 | 0.5312 | 0.2318 | Eg |
| *Protoceratomyrmex revelatus* | 0.1694 | 0.0638 | 0.6568 | 0.11 | Eg |
| *Linguamyrmex sp5* | 0.1136 | 0.0012 | 0.5652 | 0.32 | Eg |
| *Linguamyrmex sp4* | 0.1068 | 0.0036 | 0.6584 | 0.2312 | Eg |
| *Linguamyrmex brevicornis* | 0.3652 | 0.0286 | 0.3038 | 0.3024 | Ab |
| *Linguamyrmex sp2* | 0.177 | 0.0234 | 0.5522 | 0.2474 | Eg |
| *Linguamyrmex sp1* | 0.2352 | 0.014 | 0.5354 | 0.2154 | Eg |
| *Haidomyrmex sp3* | 0.4392 | 0.183 | 0.1976 | 0.1802 | Ab |
| *Haidomyrmex sp5* | 0.1836 | 0.0676 | 0.4668 | 0.282 | Eg |
| *Haidomyrmex sp5* | 0.1196 | 0.0638 | 0.611 | 0.2056 | Eg |
| *Haidomyrmex sp5* | 0.2378 | 0.079 | 0.4984 | 0.1848 | Eg |
| *Haidomyrmex sp5* | 0.31 | 0.2054 | 0.3108 | 0.1738 | Eg |
| *Haidomyrmex scimitarus* | 0.0926 | 0.0464 | 0.6744 | 0.1866 | Eg |
| *Haidomyrmex sp5* | 0.2252 | 0.0598 | 0.4966 | 0.2184 | Eg |
| *Haidomyrmex sp4* | 0.1356 | 0.0852 | 0.5698 | 0.2094 | Eg |
| *Haidomyrmex indet1* | 0.1418 | 0.0776 | 0.5776 | 0.203 | Eg |
| *Haidomyrmex indet2* | 0.1302 | 0.0806 | 0.5828 | 0.2064 | Eg |

**Table S36.**

Foraging niche prediction posterior probabilities when using model incorporating subset of complete morphometrics dataset; size-corrected ratio trait measurements; homologous morphology.

| **Species** | **Fg** | **Gn** | **GP** | **Om** | **Py** | **SP** | **Predicted aspect** |
| --- | --- | --- | --- | --- | --- | --- | --- |
| *Ceratomyrmex planus* | 0.0398 | 0.1744 | 0.1106 | 0.2534 | 0.0066 | 0.4152 | SP |
| *Ceratomyrmex ellenbergeri* | 0.036 | 0.0692 | 0.2244 | 0.432 | 0.0066 | 0.2318 | Om |
| *Ceratomyrmex ellenbergeri* | 0.035 | 0.0788 | 0.2684 | 0.32 | 0 | 0.2976 | Om |
| *Ceratomyrmex ellenbergeri* | 0.0192 | 0.1224 | 0.1888 | 0.3344 | 0.0098 | 0.3254 | Om |
| *Protoceratomyrmex revelatus* | 0.0102 | 0.0098 | 0.1038 | 0.3094 | 0.0104 | 0.5564 | SP |
| *Linguamyrmex sp5* | 0.0436 | 0.0836 | 0.2366 | 0.4418 | 0.0068 | 0.1876 | Om |
| *Linguamyrmex sp4* | 0.0288 | 0.0316 | 0.1772 | 0.3616 | 0.0072 | 0.3936 | SP |
| *Linguamyrmex brevicornis* | 0.0306 | 0.0304 | 0.1222 | 0.2642 | 0.0108 | 0.5418 | SP |
| *Linguamyrmex sp2* | 0.0474 | 0.0388 | 0.194 | 0.322 | 0.0074 | 0.3904 | SP |
| *Linguamyrmex sp1* | 0.0226 | 0.0386 | 0.236 | 0.4348 | 0.0076 | 0.2604 | Om |
| *Haidomyrmex sp3* | 0.0024 | 0.001 | 0.2646 | 0.1186 | 0.0788 | 0.5346 | SP |
| *Haidomyrmex sp5* | 0.0018 | 0.0062 | 0.3228 | 0.295 | 0.0144 | 0.3598 | SP |
| *Haidomyrmex sp5* | 0.0026 | 0.0036 | 0.3648 | 0.233 | 0.0166 | 0.3794 | SP |
| *Haidomyrmex sp5* | 0 | 0.0362 | 0.2278 | 0.2348 | 0.0382 | 0.4622 | SP |
| *Haidomyrmex sp5* | 0.0178 | 0.0274 | 0.2834 | 0.1914 | 0.0402 | 0.4398 | SP |
| *Haidomyrmex scimitarus* | 0.0136 | 0.0302 | 0.2974 | 0.316 | 0.0084 | 0.3344 | SP |
| *Haidomyrmex sp5* | 0 | 0.0166 | 0.1994 | 0.091 | 0.075 | 0.618 | SP |
| *Haidomyrmex sp4* | 0.0024 | 0.0464 | 0.2456 | 0.1742 | 0.054 | 0.4774 | SP |
| *Haidomyrmex indet1* | 0 | 0.0136 | 0.3086 | 0.2266 | 0.0214 | 0.4292 | SP |
| *Haidomyrmex indet2* | 0.003 | 0.044 | 0.195 | 0.139 | 0.0464 | 0.5726 | SP |

**Table S37.**

Functional role prediction posterior probabilities when using model incorporating subset of complete morphometrics dataset; size-corrected ratio trait measurements; homologous morphology.

| **Species** | **om_cn_ab** | **om_grll_eg** | **om_leaf** | **om_lg_ab** | **pred_cr** | **pred_gr_egll** | **pred_leaf** | **pred_lg_ab** | **pred_ll_eg** | **pred_sub** | **Predicted aspect** |
| --- | --- | --- | --- | --- | --- | --- | --- | --- | --- | --- | --- |
| *Ceratomyrmex planus* | 0.0384 | 0.2122 | 0.167 | 0.0048 | 0.0156 | 0.3348 | 0.1734 | 0.037 | 0.0086 | 0.0082 | pred_gr_egll |
| *Ceratomyrmex ellenbergeri* | 0.065 | 0.33 | 0.0928 | 0 | 0.0118 | 0.3258 | 0.159 | 0.0016 | 0.013 | 0 | om_grll_eg |
| *Ceratomyrmex ellenbergeri* | 0.042 | 0.275 | 0.0918 | 0 | 0.0188 | 0.4 | 0.1646 | 0.0022 | 0.005 | 0 | pred_gr_egll |
| *Ceratomyrmex ellenbergeri* | 0.0268 | 0.3134 | 0.0682 | 0.005 | 0.0436 | 0.3316 | 0.1656 | 0.018 | 0.0242 | 0.0036 | pred_gr_egll |
| *Protoceratomyrmex revelatus* | 0.0134 | 0.2224 | 0.003 | 0.003 | 0.012 | 0.5712 | 0.053 | 0.0196 | 0.102 | 0 | pred_gr_egll |
| *Linguamyrmex sp5* | 0.0264 | 0.3414 | 0.0824 | 0 | 0.004 | 0.2906 | 0.2442 | 0.0048 | 0.0058 | 0 | om_grll_eg |
| *Linguamyrmex sp4* | 0.021 | 0.3078 | 0.0362 | 0 | 0.0074 | 0.387 | 0.2282 | 0.0038 | 0.0078 | 0 | pred_gr_egll |
| *Linguamyrmex brevicornis* | 0.1364 | 0.1412 | 0.02 | 0.0016 | 0.0154 | 0.2688 | 0.3134 | 0.0702 | 0.0324 | 0 | pred_leaf |
| *Linguamyrmex sp2* | 0.0464 | 0.2742 | 0.0396 | 0.0012 | 0.0262 | 0.344 | 0.2514 | 0.01 | 0.0064 | 0 | pred_gr_egll |
| *Linguamyrmex sp1* | 0.0456 | 0.312 | 0.0502 | 0.0016 | 0.021 | 0.338 | 0.2114 | 0.0028 | 0.0172 | 0 | pred_gr_egll |
| *Haidomyrmex sp3* | 0.0076 | 0.0364 | 0 | 0.0766 | 0.0324 | 0.1066 | 0.1654 | 0.5032 | 0.0552 | 0.0158 | pred_lg_ab |
| *Haidomyrmex sp5* | 0.0122 | 0.2678 | 0.0028 | 0.0114 | 0.0102 | 0.3234 | 0.2796 | 0.0332 | 0.0586 | 0 | pred_gr_egll |
| *Haidomyrmex sp5* | 0.0046 | 0.209 | 0.0018 | 0.0068 | 0.013 | 0.5734 | 0.1688 | 0.0108 | 0.0102 | 0.0016 | pred_gr_egll |
| *Haidomyrmex sp5* | 0.008 | 0.2518 | 0 | 0.0352 | 0.02 | 0.3826 | 0.1464 | 0.0668 | 0.082 | 0.0068 | pred_gr_egll |
| *Haidomyrmex sp5* | 0.0284 | 0.1194 | 0.0216 | 0.0964 | 0.0712 | 0.1498 | 0.1526 | 0.2766 | 0.072 | 0.012 | pred_lg_ab |
| *Haidomyrmex scimitarus* | 0.0258 | 0.2276 | 0.0248 | 0.002 | 0.0206 | 0.5198 | 0.1688 | 0.0052 | 0.005 | 0 | pred_gr_egll |
| *Haidomyrmex sp5* | 0 | 0.0894 | 0 | 0.122 | 0.0102 | 0.3502 | 0.104 | 0.1666 | 0.1328 | 0.0236 | pred_gr_egll |
| *Haidomyrmex sp4* | 0.0102 | 0.1858 | 0.0022 | 0.016 | 0.0342 | 0.4946 | 0.1404 | 0.0396 | 0.0736 | 0.0034 | pred_gr_egll |
| *Haidomyrmex indet1* | 0.0036 | 0.2082 | 0 | 0.018 | 0.0268 | 0.5342 | 0.1424 | 0.0396 | 0.0224 | 0.0044 | pred_gr_egll |
| *Haidomyrmex indet2* | 0 | 0.1146 | 0.0048 | 0.0846 | 0.0274 | 0.426 | 0.1068 | 0.148 | 0.081 | 0.006 | pred_gr_egll |

**Table S38.**

Ecomorph prediction posterior probabilities when using model incorporating subset of complete morphometrics dataset; size-corrected ratio trait measurements; homologous morphology.

| **Lineage** | **Average ecological disparity** | **Standard deviation of ecological disparity** |
| --- | --- | --- |
| Ponerines (*Anochetus* and *Odontomachus*) | 4.701 | 2.553 |
| Dacetines (*Acanthognathus, Daceton, Epopostruma, Microdaceton,* and *Orectognathus*) | 4.667 | 2.570 |
| *Strumigenys* | 3.200 | 1.600 |
| *Myrmoteras* | 1.333 | 0.471 |
| Haidomyrmecines (conservative estimate) | 2.944 | 1.632 |
| Haidomyrmecines (comprehensive estimate) | 3.448 | 1.746 |

**Table S39.**

Average and standard deviation of ecological disparity calculated as pairwise distances between each unique three-dimensional occupation. Haidomyrmecine conservative estimates are derived from only the majority aspect predicted by the raw functional measurement model, while comprehensive estimates are derived from majority aspects predicted from all four RF models.

**Data S1.** Supplementary Data_Fossil Morphometrics (separate file)

All fossil morphometric data for all fossil specimens included in this study. The Excel file has four tabs: one with all raw trait measurements measured in a functional morphological framework (“raw functional”); one with all raw trait measurements measured in a homologous morphological framework (“raw homologous”); one with all size-corrected ratio trait measurements measured in a functional morphological framework (“ratio functional”); and one with all size-corrected ratio trait measurements measured in a homologous morphological framework (“raw homologous”).

**Data S2.** Supplementary Data_ Trap Jaw Ecomorphospace (separate file)

All body size and ecological niche data used for calculating ecomorphospace occupation in extinct and extant trapjaw ant taxa. The Excel file has tabs for each genus of trapjaw ants, which were combined for cases where multiple genera were treated as one lineage. “MinHaidomyrmecinae” is collated solely from predictions estimated under a raw trait measurement functional morphology framework, while “MaxHaidomyrmecinae” is collated from predictions estimated using all models.

**Data S3.** Supplementary Data_Random Forest Code (separate file)

R code that may be used to reproduce the Random Forest analyses herein, commented for clarity.

**Data S4.** Supplementary Data_Specimen Images (separate file)

Photomicrographs with species and morphospecies identification for all specimens used in this study that are currently residing in a private collection.
