## Supplementary Data: Specimen Images for "Trait-based paleontological niche prediction demonstrates deep time parallel ecological occupation in specialized ant predators"

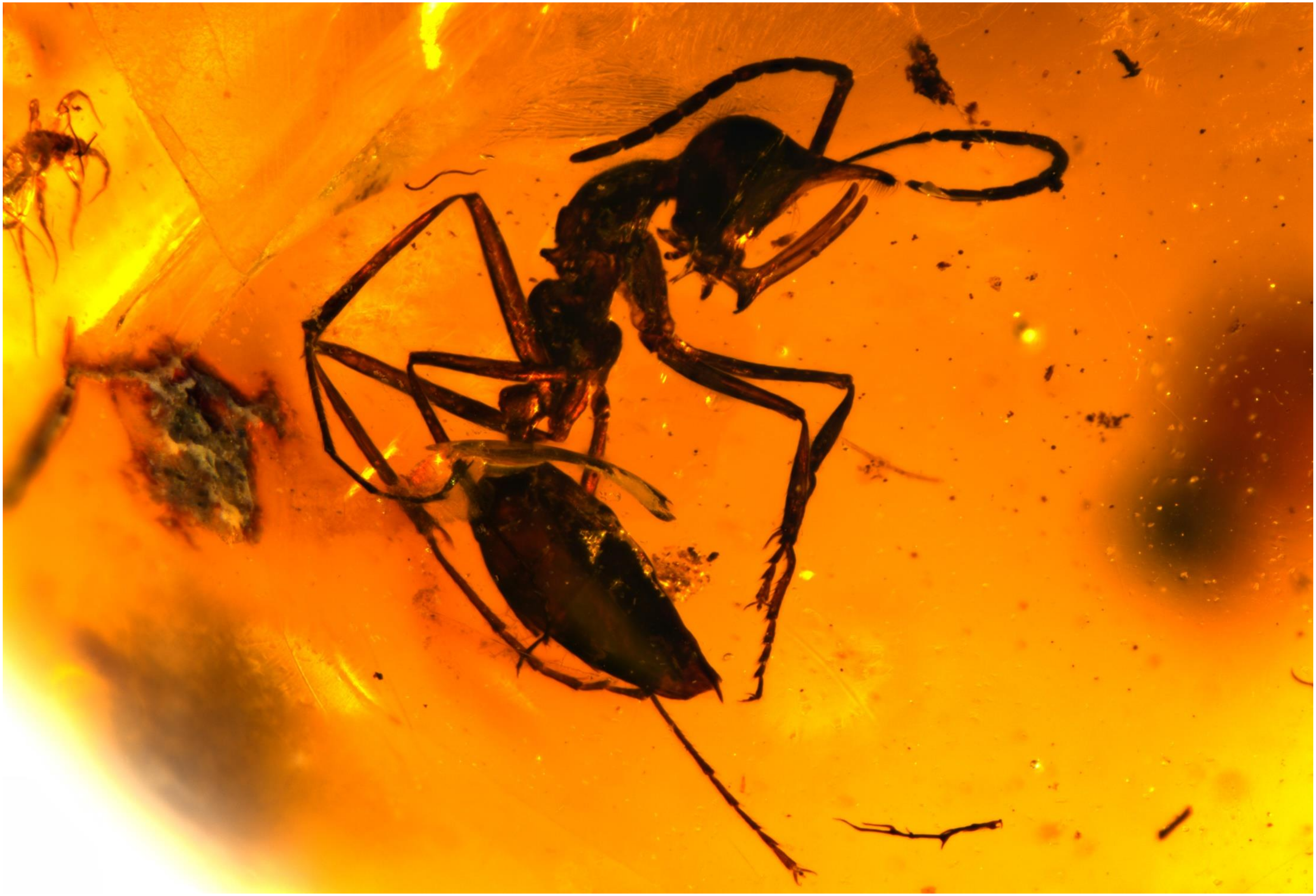

BALTJ_025: *Ceratomyrmex planus*

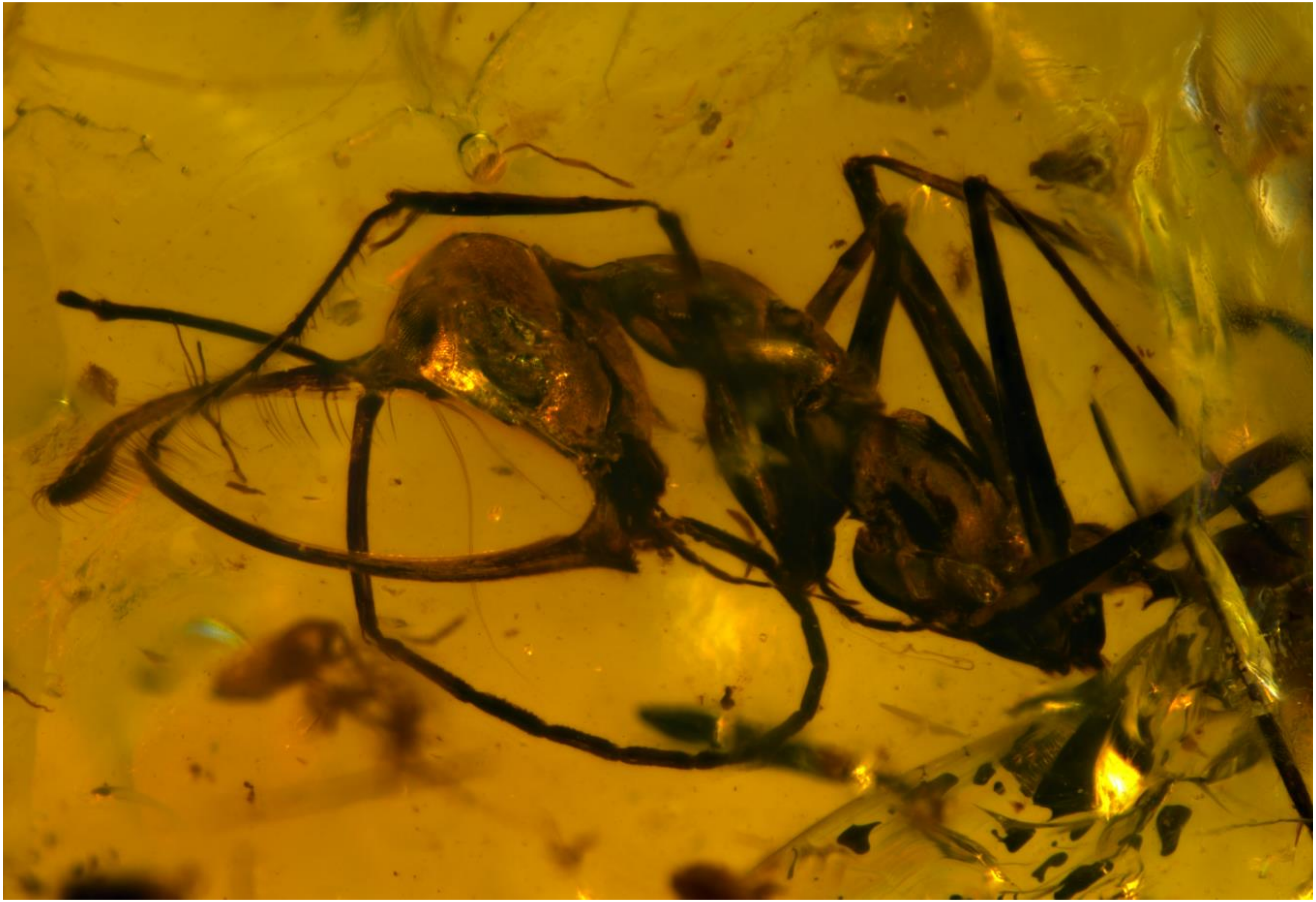

BALTJ_016: *Ceratomyrmex ellenbergeri*.

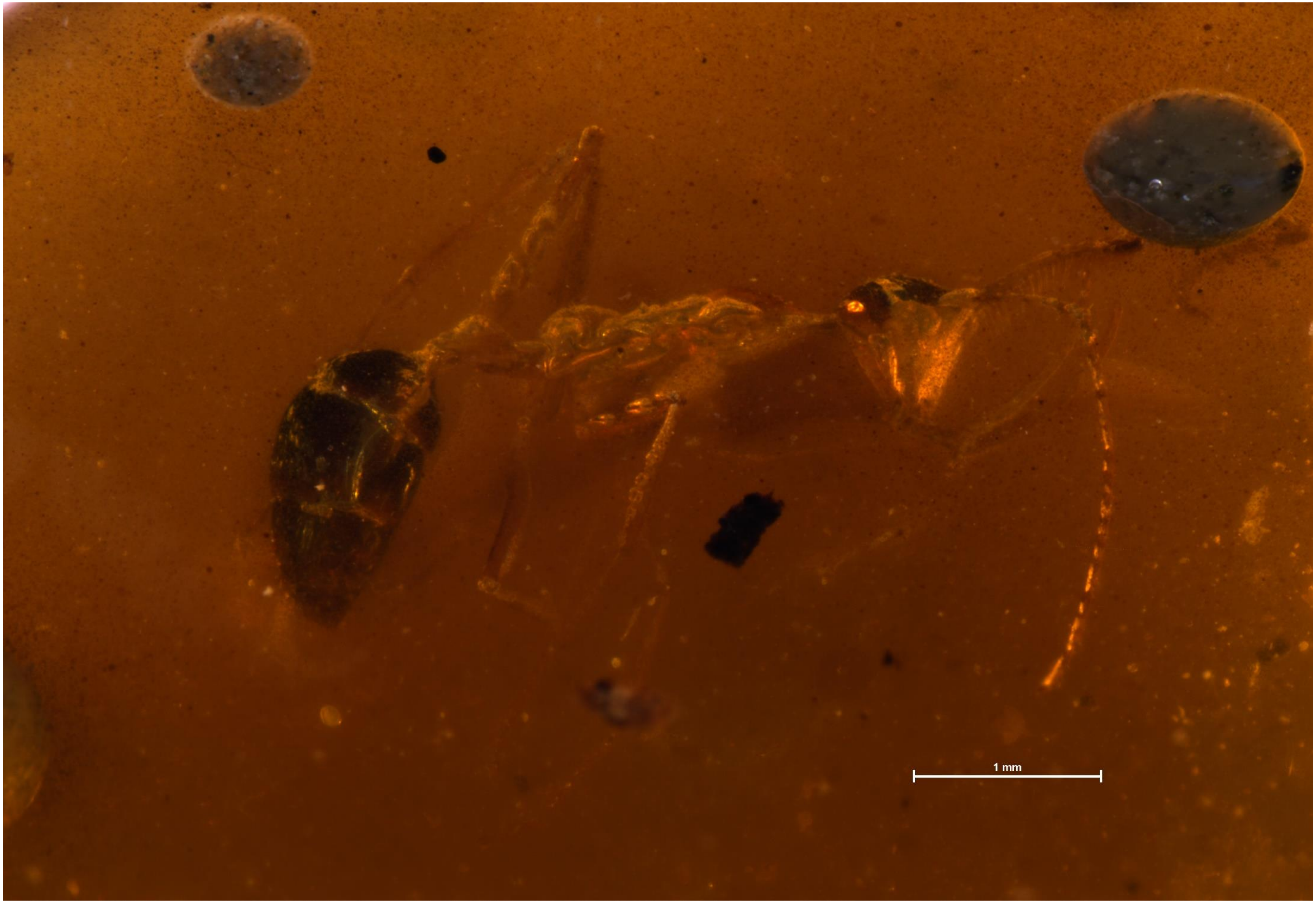


BALTJ_027: *Ceratomyrmex ellenbergeri*

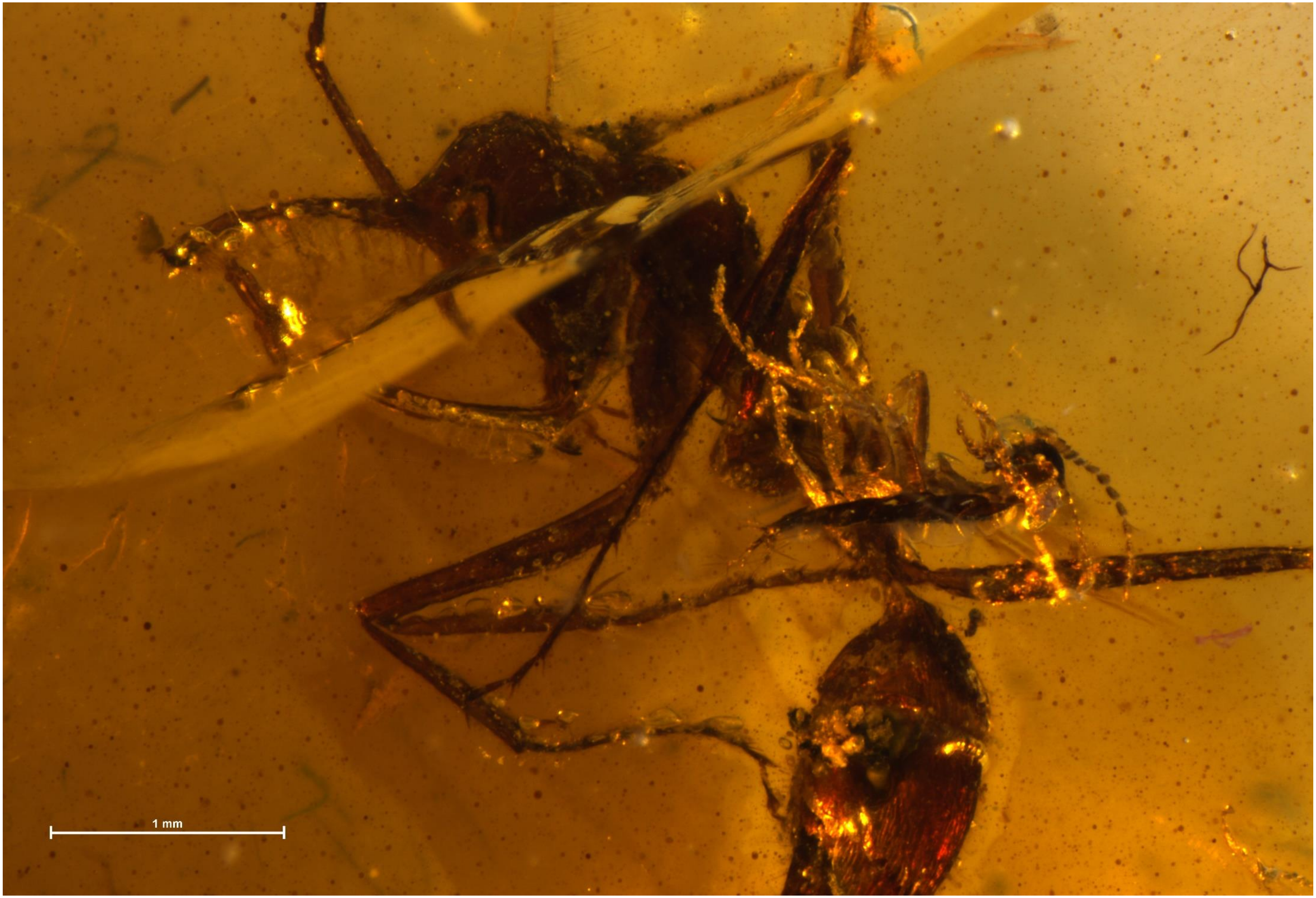


BALTJ_029: *Ceratomyrmex ellenbergeri*

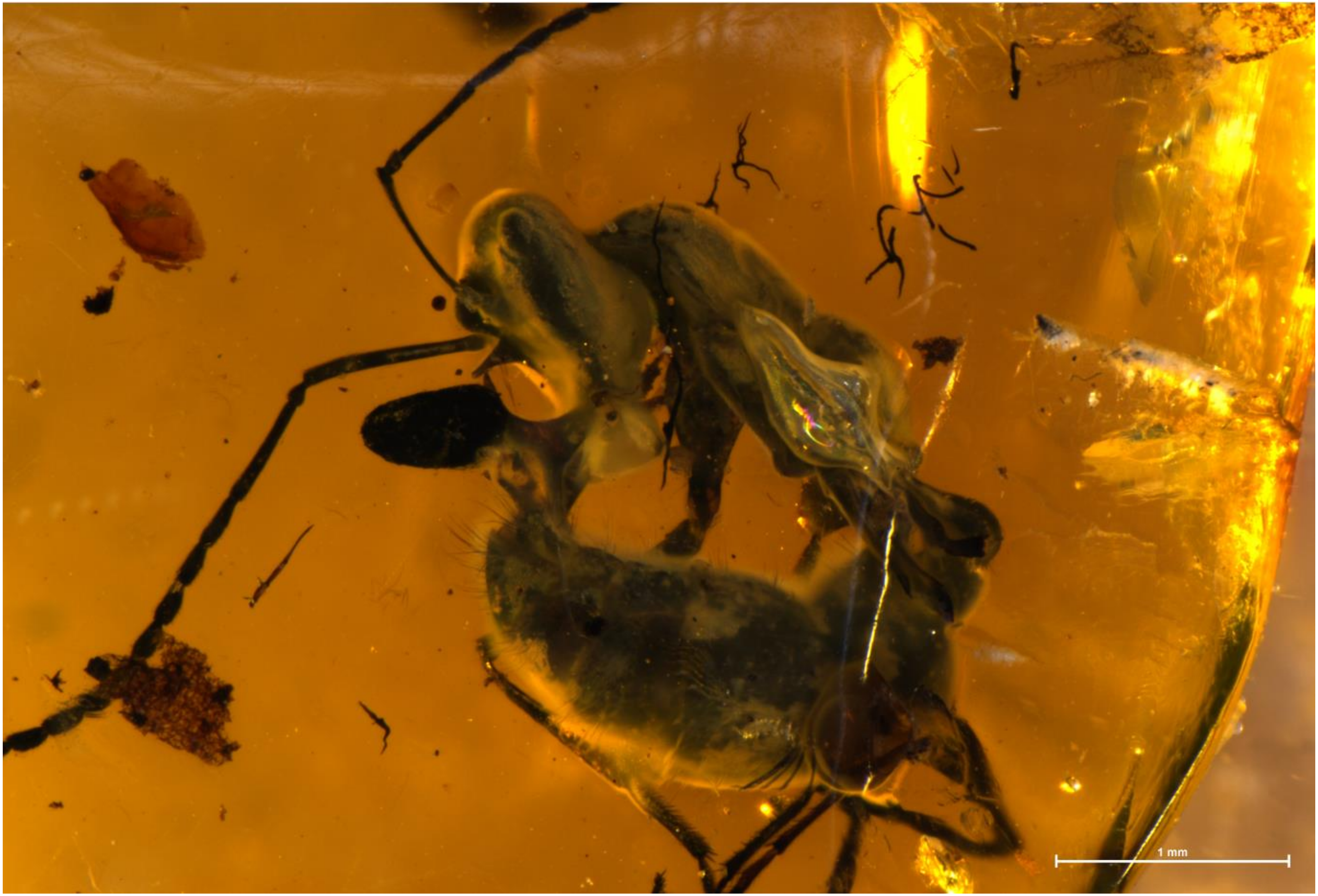


BALTJ_043: *Protoceratomyrmex revelatus*

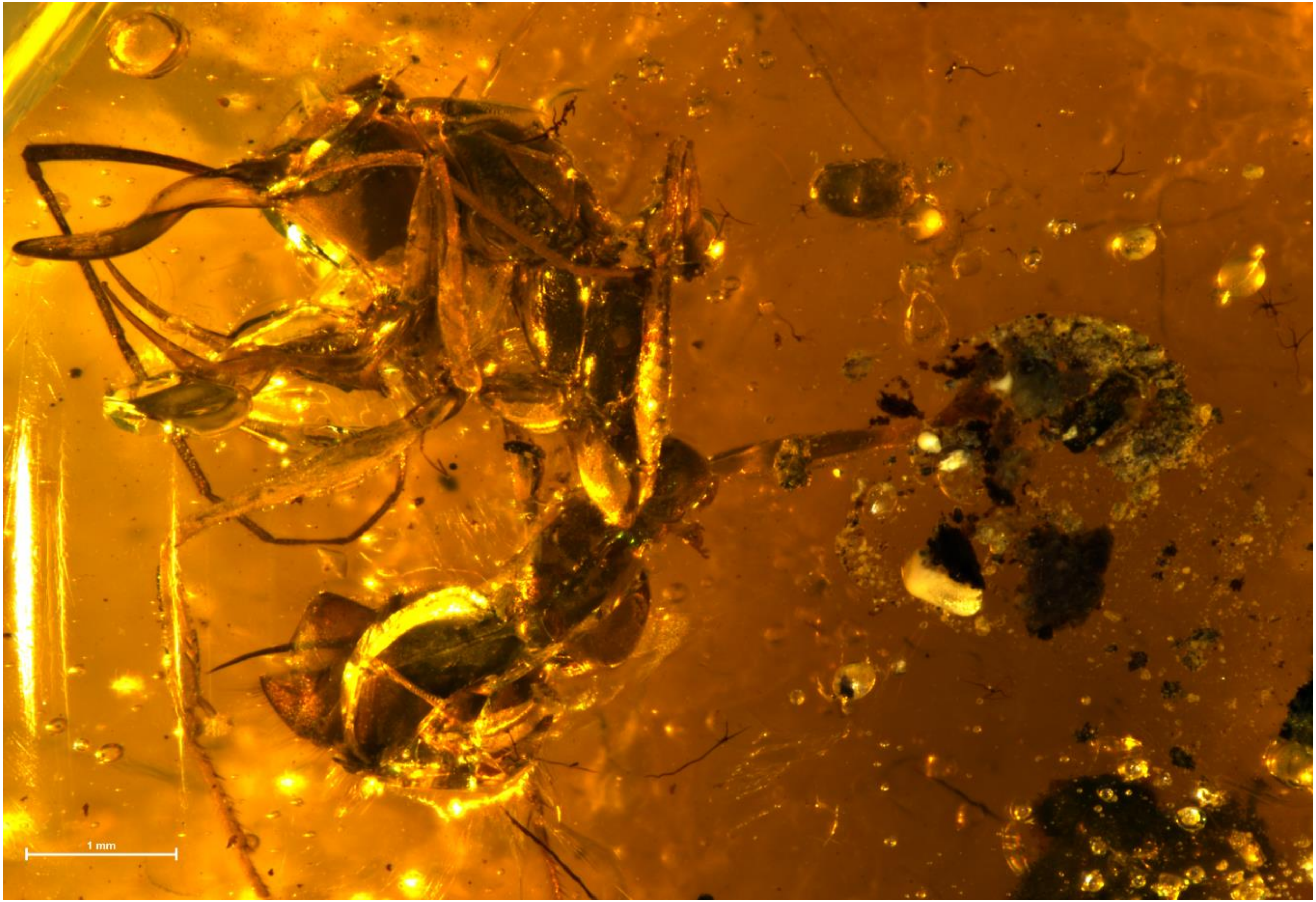


BALTJ_050: *Linguamyrmex sp5*

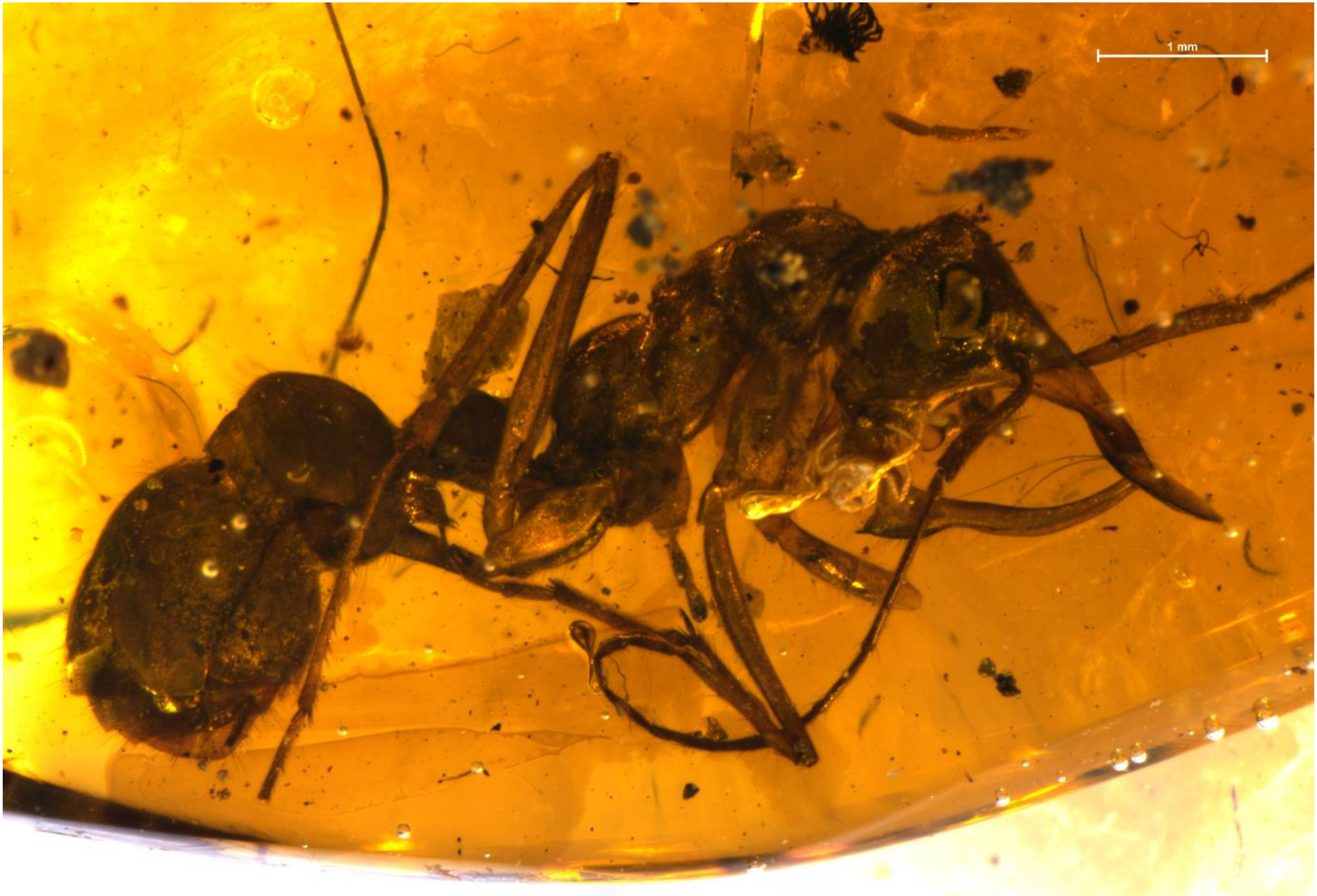


BALTJ_031: *Linguamyrmex sp4.*

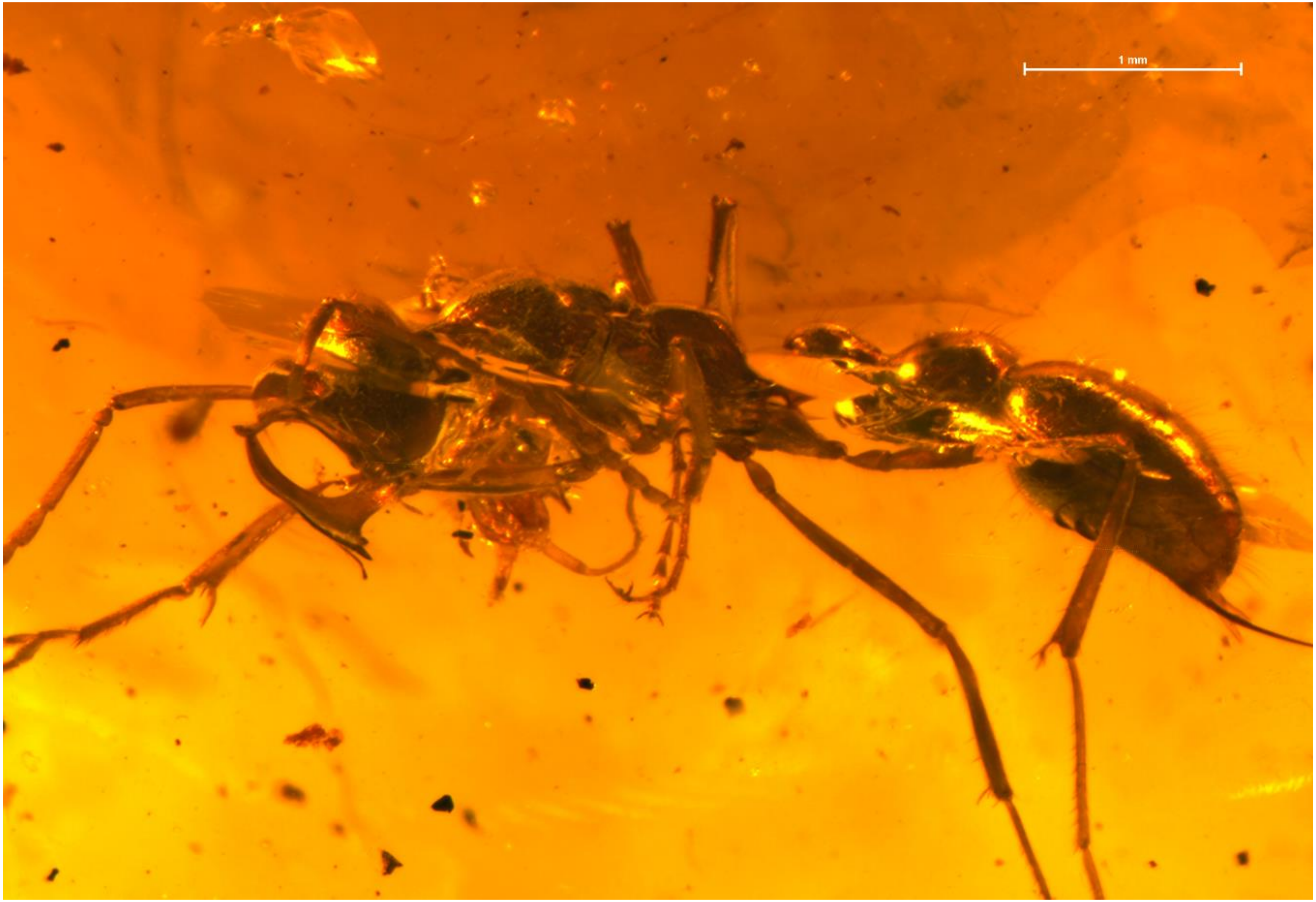


BALTJ_030: *Linguamyrmex brevicornis.*

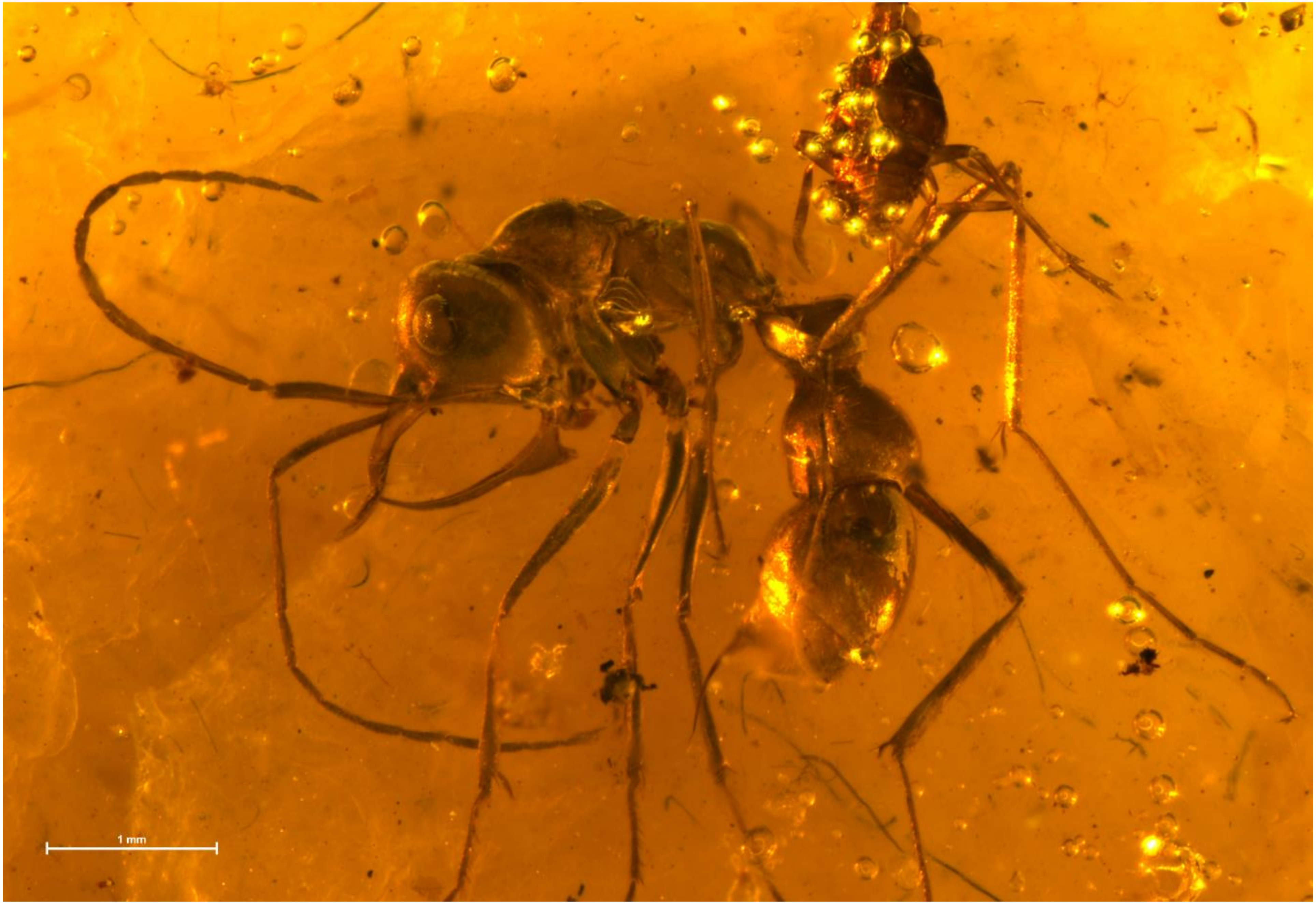


BALTJ_059: *Linguamyrmex sp2*

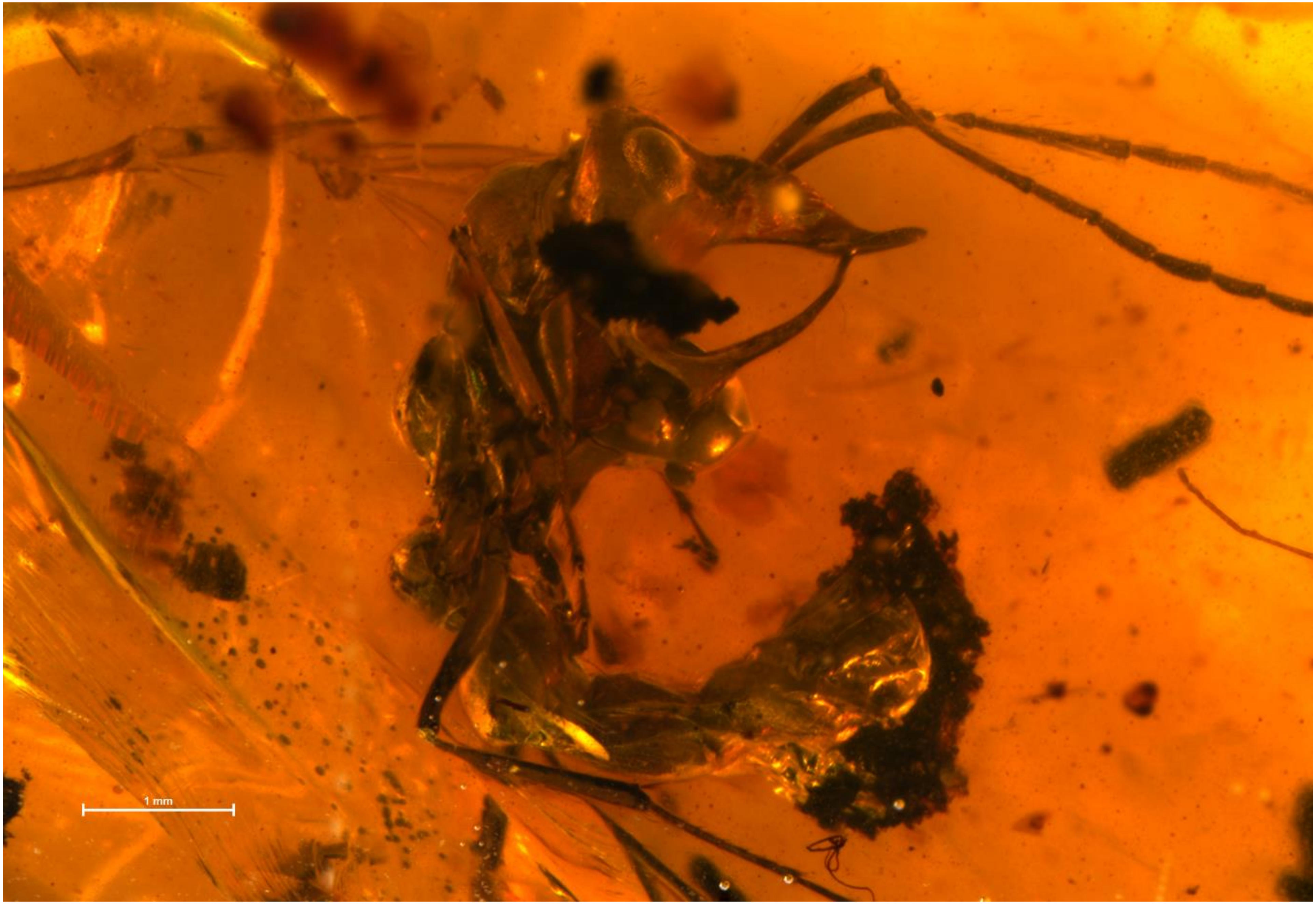


BALTJ_056: *Linguamyrmex sp1*

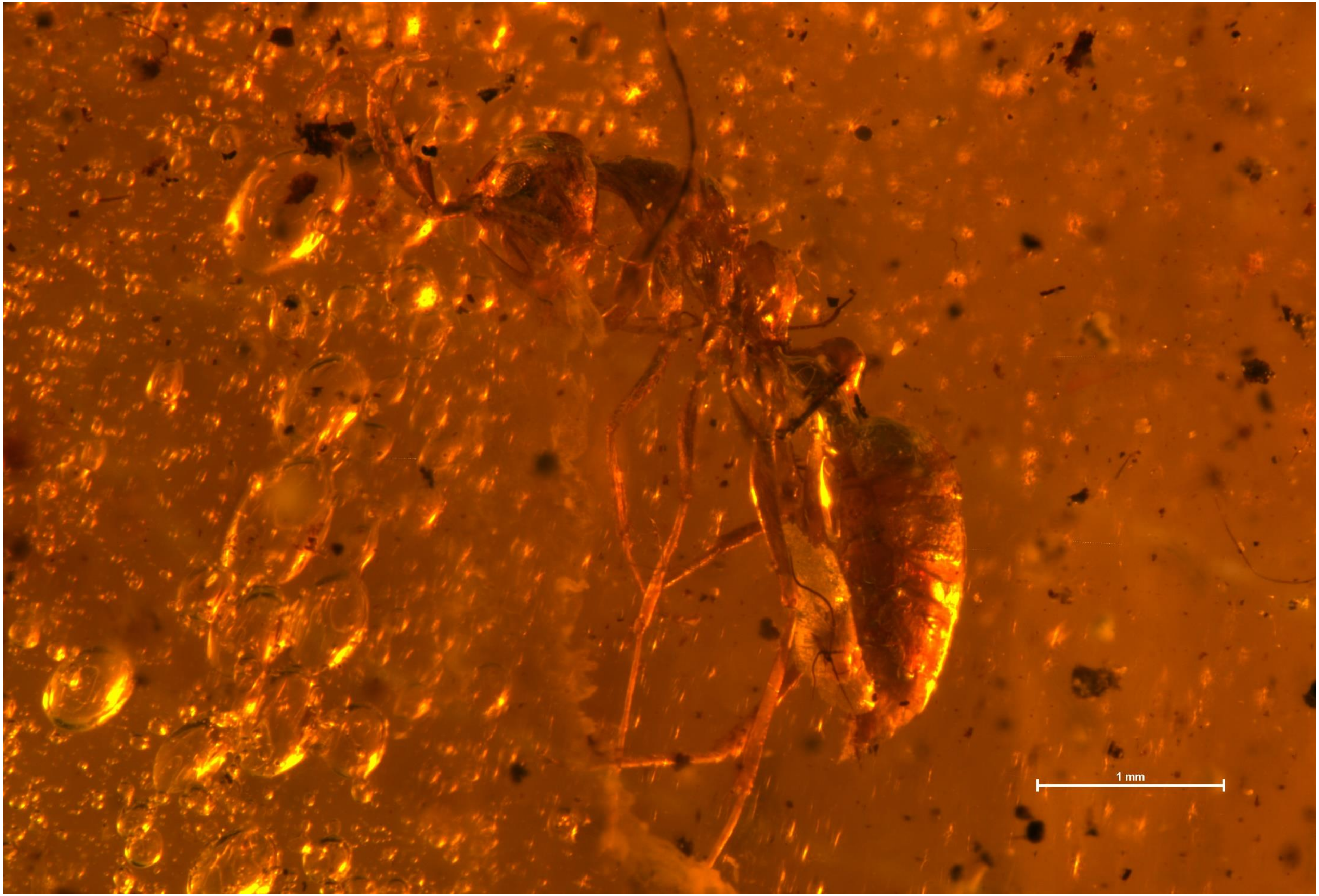


BALTJ_022: *Haidomyrmex sp3*

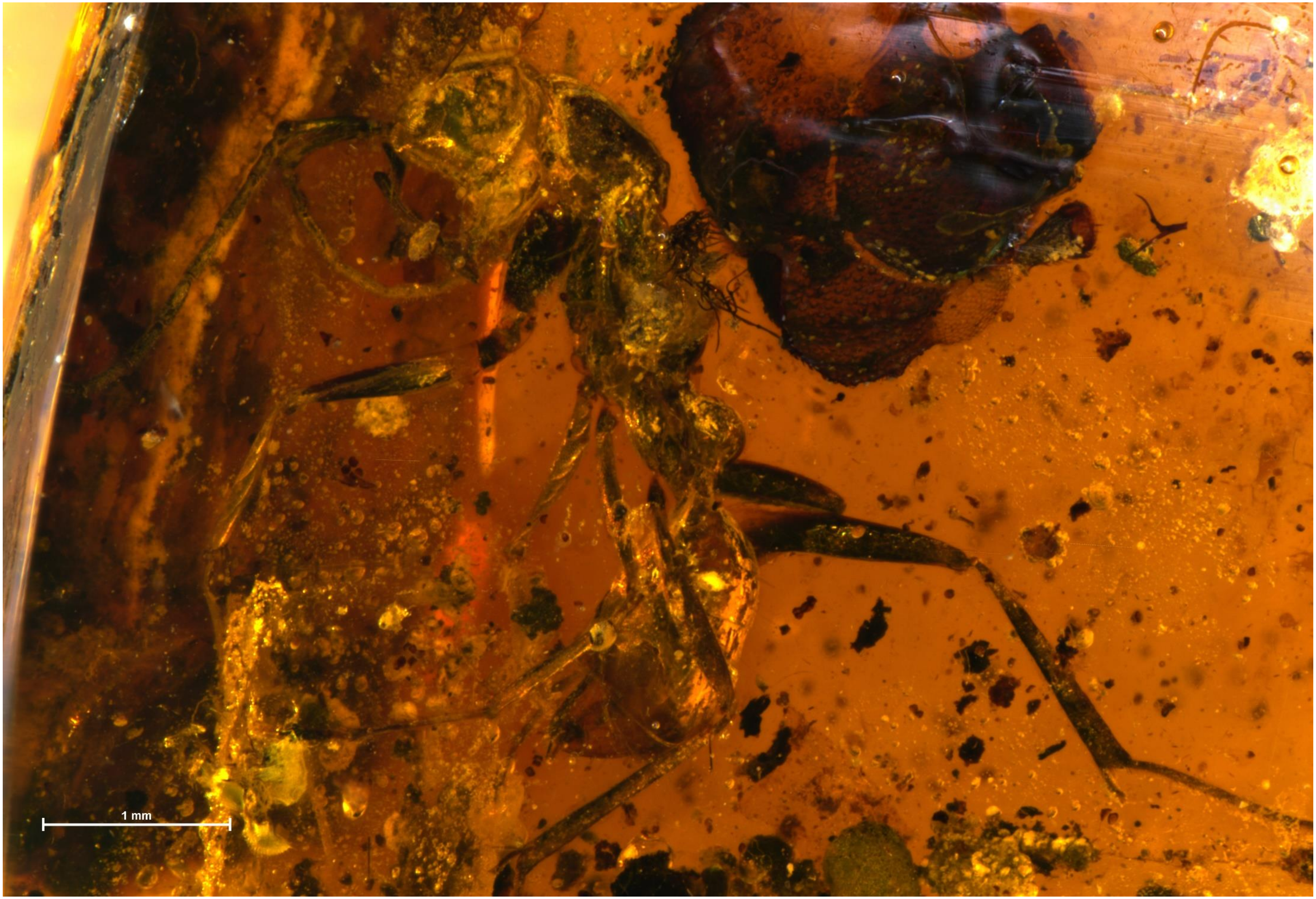


BALTJ_017: *Haidomyrmex sp5*

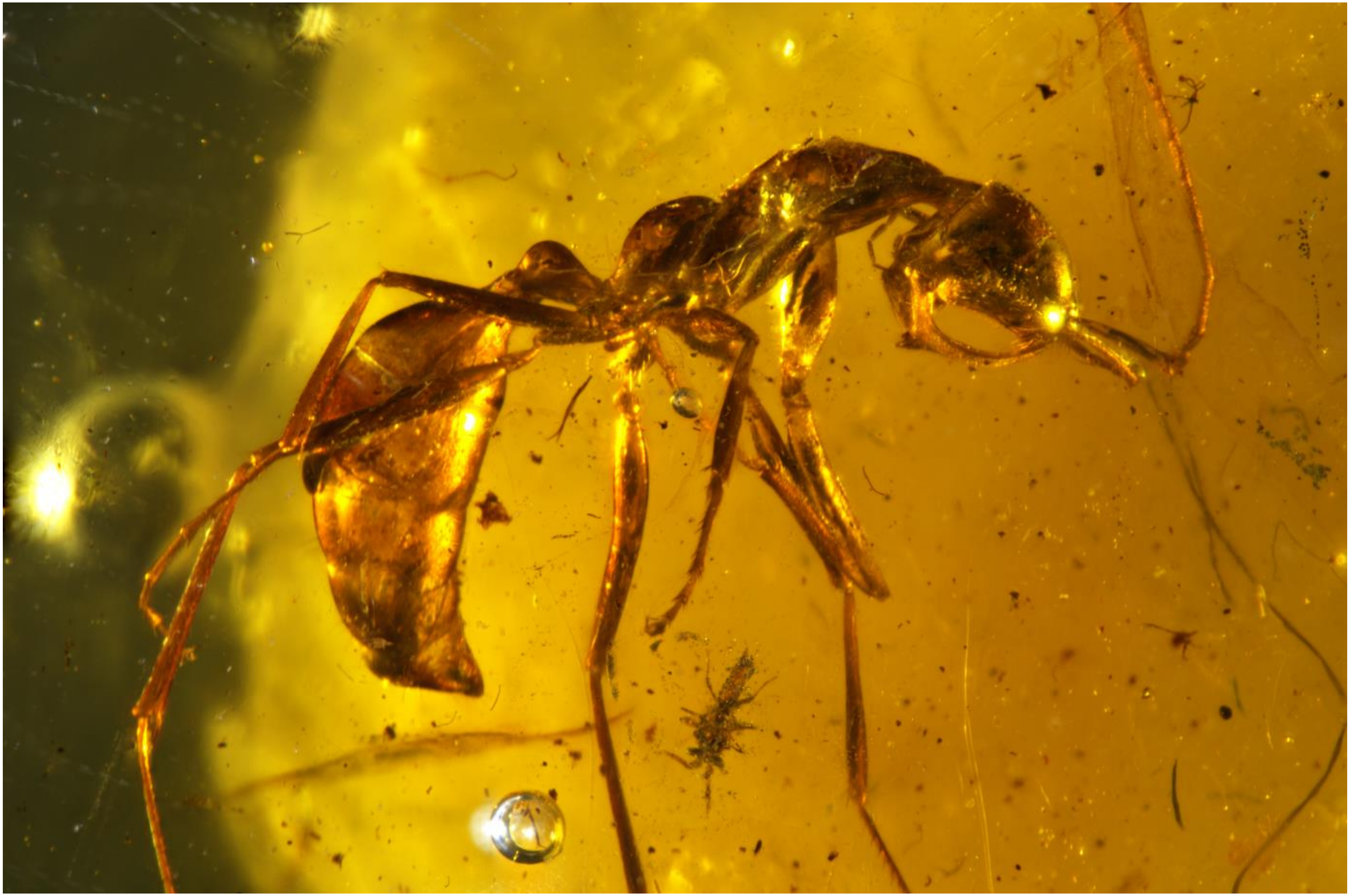

BALTJ_009: *Haidomyrmex sp5*

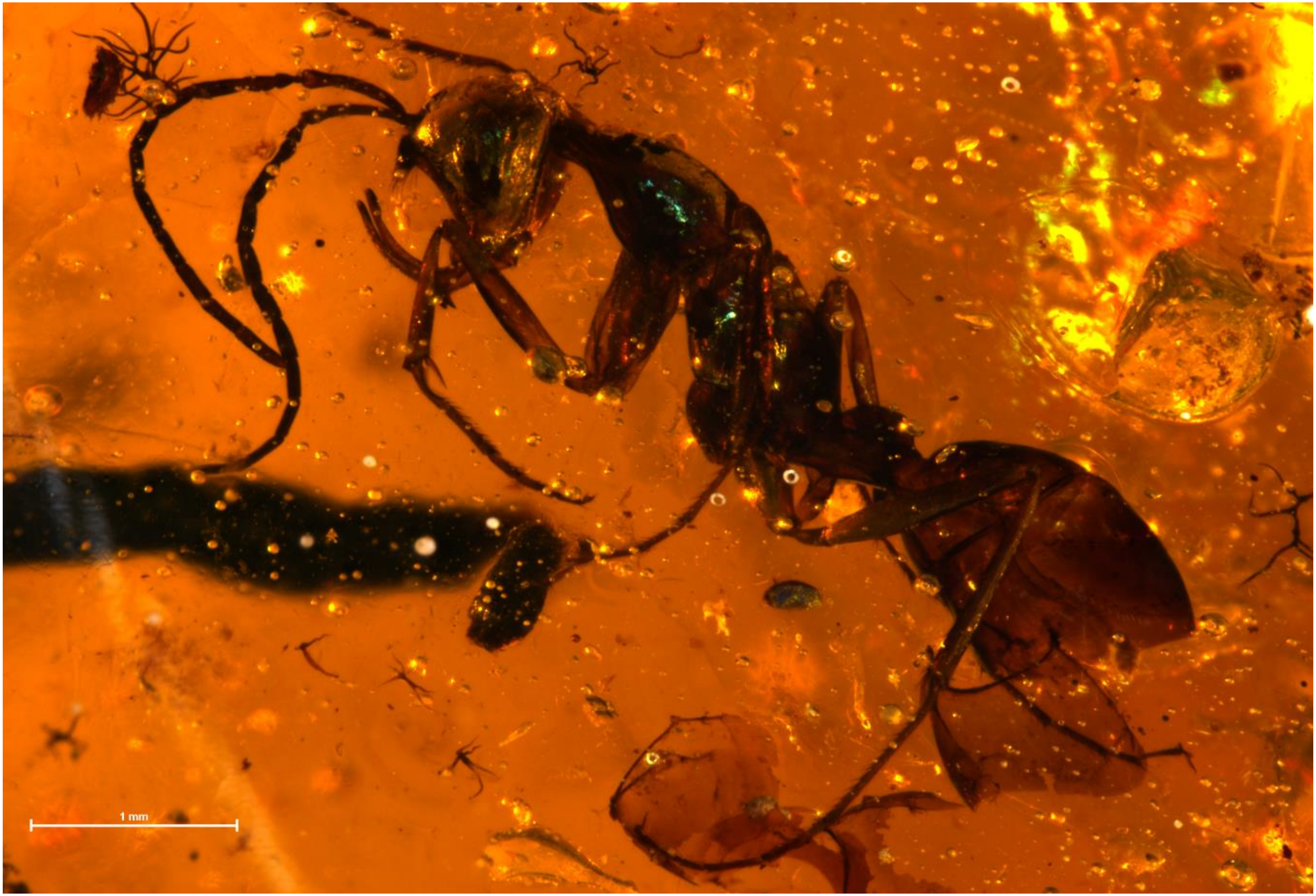


BALTJ_066: *Haidomyrmex sp5*

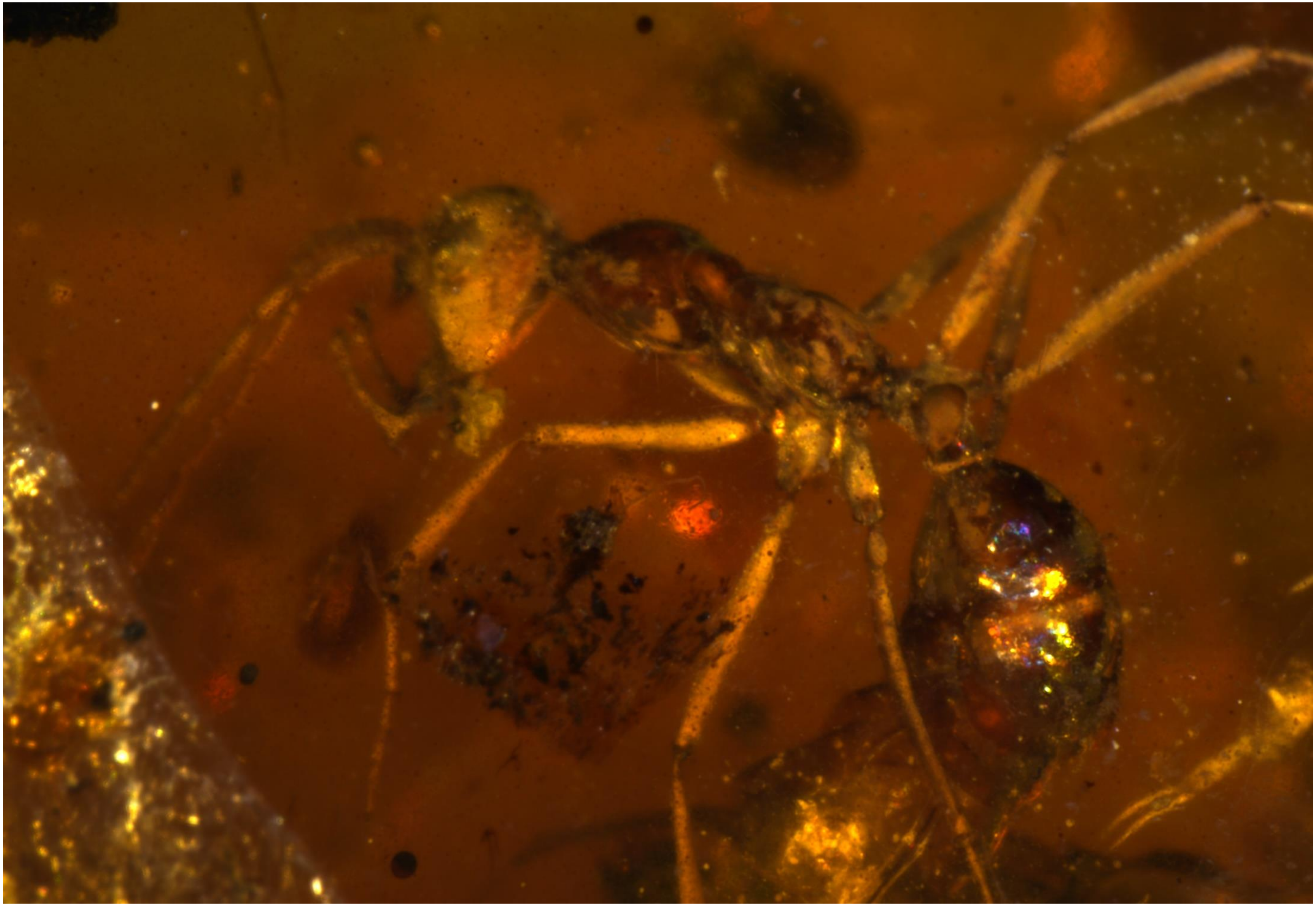

BALTJ_007: *Haidomyrmex sp5*

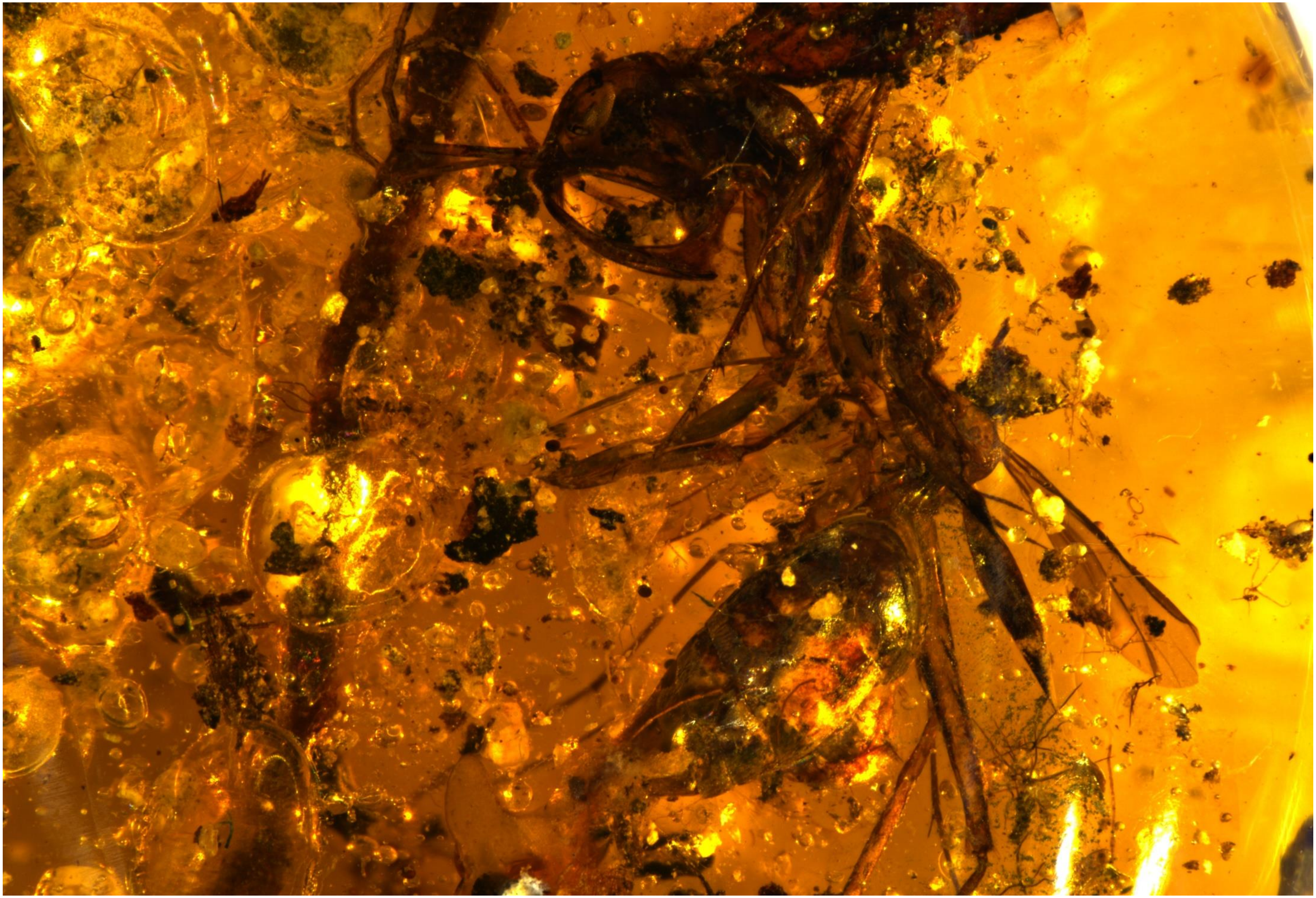

BALTJ_004: *Haidomyrmex scimitarus*

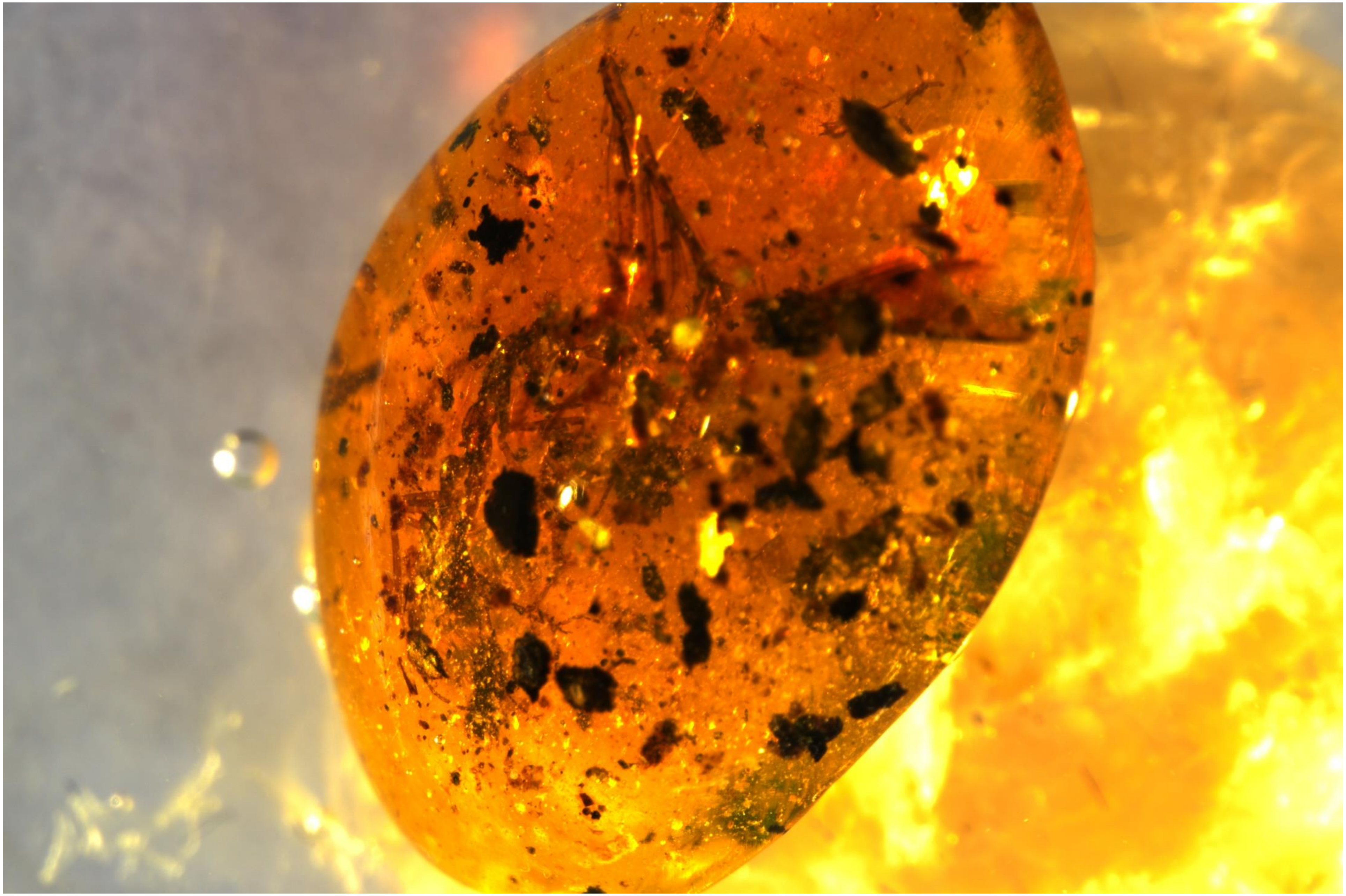

BALTJ_011: *Haidomyrmex sp5*

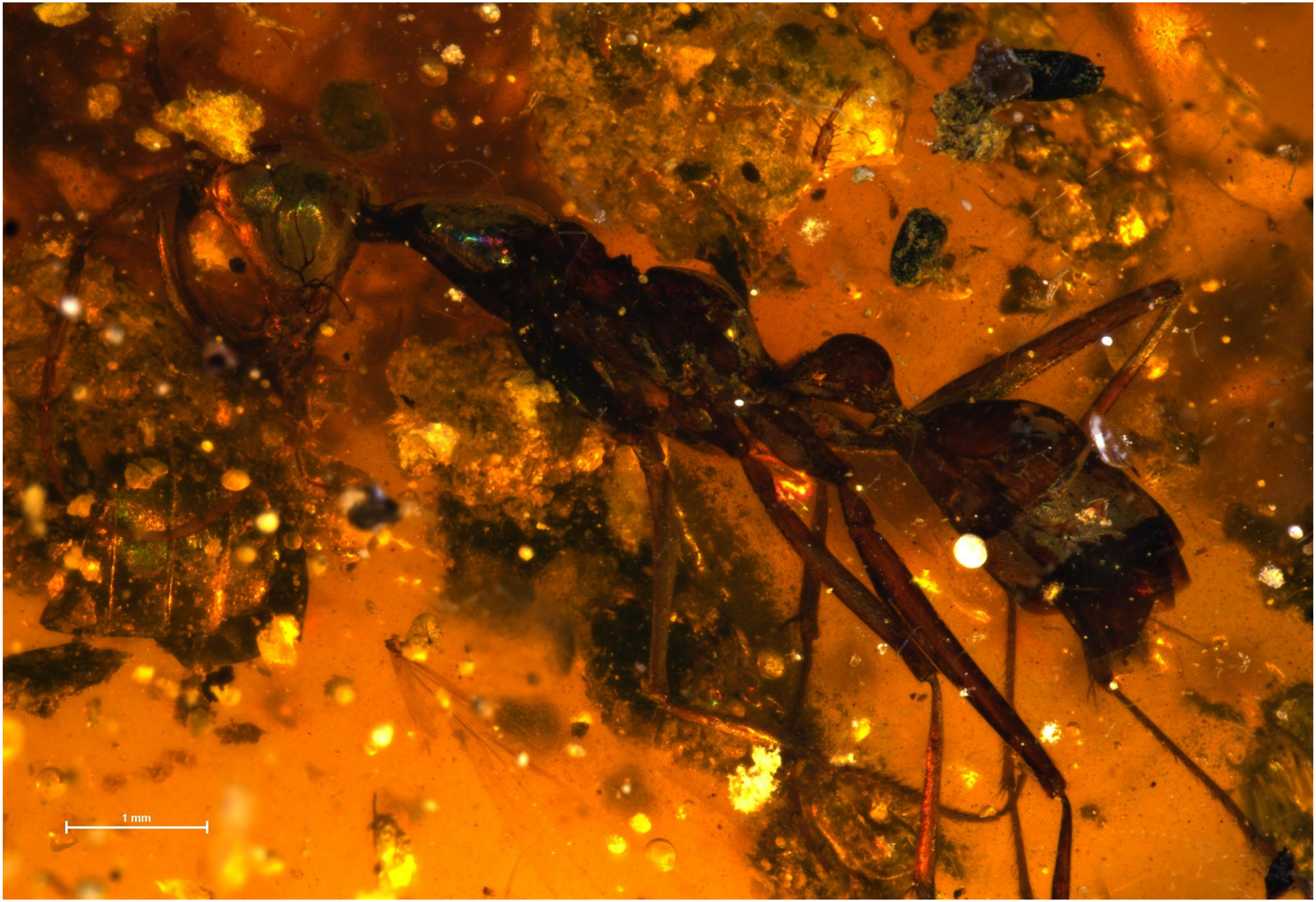


BALTJ_003: *Haidomyrmex sp4*

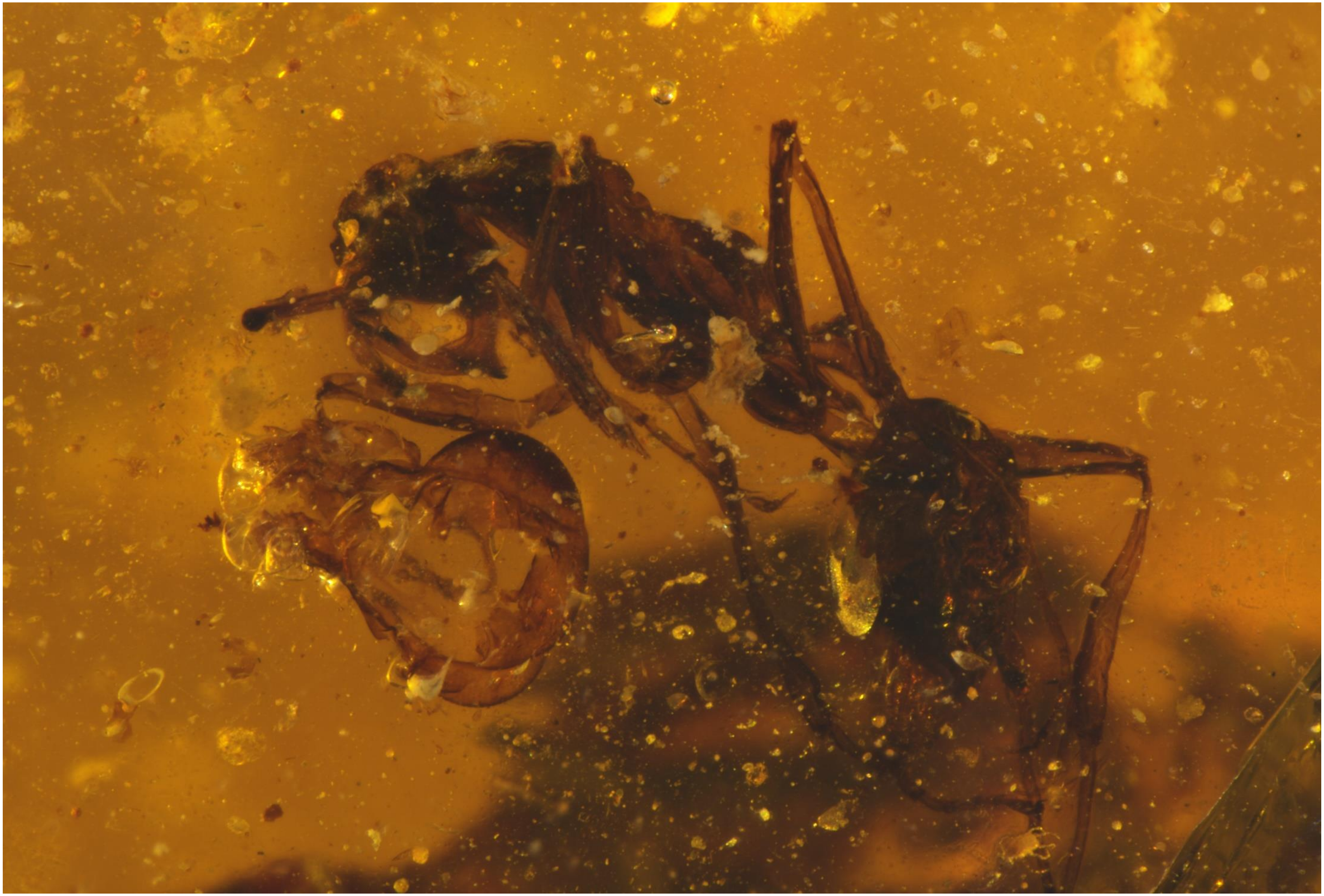

BALTJ_010: *Haidomyrmex indet1*

BALTJ_065: Haidomyrmex indet2
